# capCLIP: a new tool to probe protein synthesis in human cells through capture and identification of the eIF4E-mRNA interactome

**DOI:** 10.1101/2020.04.18.047571

**Authors:** Kirk B. Jensen, B. Kate Dredge, John Toubia, Xin Jin, Valentina Iadevaia, Gregory J. Goodall, Christopher G. Proud

## Abstract

Translation of eukaryotic mRNAs starts with binding of the m^7^G cap to the protein eIF4E followed by recruitment of other translation initiation factors. eIF4E’s essential role in translation suggests the cellular eIF4E-mRNA interactome (or ‘eIF4E cap-ome’) may serve as a faithful proxy of cellular translational activity. Here we describe capCLIP, a novel method to systematically capture and quantify the eIF4E cap-ome. To validate capCLIP, we identified the cap-omes in human cells ± the partial mTORC1 inhibitor rapamycin. As expected, TOP (terminal oligopyrimidine) mRNA representation is systematically reduced in rapamycin-treated cells. capCLIP tag data permits refinement of a 7-nucleotide TOP motif (5′-CUYUYYC-3′). We also apply capCLIP to probe the consequences of phosphorylation of eIF4E, whose function had remained unclear. eIF4E phosphorylation drives an overall reduction in eIF4E-mRNA association; strikingly, mRNAs most sensitive to phosphorylation possess short 5′-UTRs. capCLIP provides a sensitive and comprehensive measure of cellular translational activity. We foresee its application as a high-throughput way to assess translation in contexts not amenable to existing methodologies.

## Introduction

Control of protein synthesis (mRNA translation) is a key regulatory mechanism used by eukaryotic cells to control the composition of the cellular proteome (Proud, 2019). The translational control mechanisms used by mammalian cells are diverse and offer the ability to regulate specific mRNAs or subsets of mRNAs. Such mechanisms act primarily at the initiation phase of translation, where ribosomes are recruited to mRNAs (Proud, 2019). eIF4E (eukaryotic initiation factor 4E) is of central importance both for translation initiation and its regulation (Merrick and Pavitt, 2018). eIF4E binds the 7-methyl-GTP (m^7^GTP) moiety of the 5′-cap structure found on all cellular cytoplasmic mRNAs. This interaction is crucial for efficient translation of almost all mRNAs (Rhoads, 2009). eIF4E binds the scaffold protein eIF4G, along with other eIFs, including eIF3. This eIF4E-eIF4G-eIF3 complex then recruits 40S ribosomal subunits to the 5′ end of the mRNA, permitting scanning of 40S-associated initiation complexes to the AUG, 60S subunit joining, and subsequent translation of the message (Merrick, 2015). While eIF4E^+/−^ mice are viable, eIF4E^+/−^ embryonic fibroblasts (MEFs) are resistant to oncogenic transformation (Truitt et al., 2015), indicating that cellular translational activity is very sensitive to the amount of eIF4E protein.

As the cap-eIF4E interaction is essential for cap-dependent translation, it seems reasonable that identification of the mRNAs bound to eIF4E (the eIF4E ‘cap-ome’) should be a ‘readout’ of cellular translational activity. To test this, we developed a new methodology called capCLIP which complements existing high-throughput approaches such as ribosome profiling. capCLIP is a cross-linking immunoprecipitation (CLIP)-based method (Ule et al., 2005; Ule et al., 2003) that takes advantage of the fact that UV irradiation of living cells can generate a photo-crosslink between eIF4E and the m^7^G cap, thus covalently linking eIF4E to bound, capped mRNAs (Pelletier and Sonenberg, 1985). Immunoprecipitation (IP) and subsequent purification of eIF4E selectively co-purify this crosslinked RNA; cloning and sequencing of this material identifies the cellular eIF4E cap-ome.

To demonstrate capCLIP’s potential as both a novel means to identify a cell’s translationally active mRNAs and as a means to reveal new insights into translational control, we use it to probe how two major signalling pathways shape translational activity in the cell. In the first study, we examine how rapamycin, a partial inhibitor of mTORC1 (mechanistic target of rapamycin, complex 1) impacts the eIF4E cap-ome. mTORC1 is a multi-subunit protein kinase that is activated by amino acids, growth factors and other trophic stimuli. It is a major regulator of translation initiation and ribosome biogenesis (Saxton and Sabatini, 2017). A major consequence of mTORC1 inhibition by rapamycin or drugs which act as direct inhibitors of mTOR kinase activity (‘mTOR-KIs’) is the marked impairment of synthesis of ribosomal proteins (Huo et al., 2012), which are encoded by mRNAs that possess a 5′-terminal tract of oligopyrimidines (5′-TOP) (Meyuhas and Kahan, 2015). While the precise features of a functional TOP motif remain to be established, it is generally agreed that TOP mRNA transcripts always begin with a C nucleotide, followed by 4-15 pyrimidines (Meyuhas and Kahan, 2015).

Using Hela cells as a model, we identify an eIF4E cap-ome of approximately 4,000 unique mRNAs. Upon rapamycin treatment, 86 mRNAs are significantly depleted from the eIF4E cap-ome; 62 (72%) of the 86 mRNAs are TOP mRNAs. We then mapped the transcription start sites (TSS) of 52 TOP mRNAs, which allowed us to determine the precise relationship between a TOP motif’s ‘strength’ (the degree of change in eIF4E-mRNA binding caused by rapamycin) and its sequence. Our analysis indicates that mRNAs exhibiting strong TOP functionality share a highly-conserved, 7-nucleotide pyrimidine consensus sequence. capCLIP thus provides us with a completely novel means to determine the influence of 5′-end sequence on a mRNA’s translational activity, which should prove valuable in a number of experimental contexts. As considerable understanding of the translational consequences of mTORC1 signalling are available from ribosome footprinting/profiling studies, we also compare our eIF4E cap-ome data with the relevant studies (Hsieh et al., 2012; Thoreen et al., 2012). As this analysis affirms that the eIF4E cap-ome does indeed represent translationally active mRNAs, and as capCLIP also provides previously inaccessible insights into individual eIF4E-mRNA interactions, the methodology promises to provide detailed insights into how other events which affect eIF4E-mRNA binding shape translation.

Our second study endeavours to redeem this promise by using capCLIP to probe the poorly-understood translational and biological consequences of phosphorylation of eIF4E itself. eIF4E undergoes a single phosphorylation event on Ser209, catalysed by the MNKs (MAP kinase-interacting kinases; MNK1 and MNK2). MNKs are directly activated by p38 MAP kinase and/or ERK (Proud, 2015). eIF4E is the only known *in vivo* substrate of the MNKs (Xie et al., 2019). There is an association between elevated P-eIF4E levels and cancer (Xie et al., 2019), and interestingly, eIF4E^S209A/S209A^ knock-in mice are resistant high-grade prostate intraepithelial neoplasia (PIN). While a few mRNAs have been identified whose polysomal association appears to *increase* with P-eIF4E, both the mechanistic basis for these differences and the translational consequences of eIF4E Ser209 phosphorylation remain unclear (Furic et al., 2010).

Using the highly-selective MNK inhibitor eFT-508 (Reich et al., 2018) to block eIF4E phosphorylation in human cells, we use capCLIP to identify the effect of blocking eIF4E phosphorylation on the cellular cap-ome. Intriguingly, inhibition of eIF4E phosphorylation leads to an average ~1.7 fold *decrease* in eIF4E-mRNA binding across the 5,000 unique mRNAs of the eIF4E cap-ome, with 256 of these mRNAs exhibiting a statistically significant reduction in eIF4E binding. Strikingly, mRNAs with the most reduced eIF4E association possess significantly shorter than average 5′ UTRs. Our findings are consistent with *in vitro* measurements showing that phosphorylation of eIF4E lowers its affinity for capped mRNA and cap analogues (Scheper et al., 2002; Slepenkov et al., 2006; Zuberek et al., 2003).

In sum, we show that eIF4E capCLIP provides a novel and valid way to characterise cap-dependent translational activity in human cells, and can be used to comprehensively and quantitatively measure the impact of inhibition of two distinct cellular signalling pathways that control the translational machinery. We predict that capCLIP will see widespread use to provide comprehensive data on the translational activity of cells, and potentially in tissue.

## Results and Discussion

### Capture and identification of the eIF4E-mRNA interactome

The 7 steps of the capCLIP method are shown schematically in Figure 1A. The steps are further described in the Star Methods, and the complete protocol is available as a supplemental document. In brief, capCLIP uses short-wavelength (~254 nm) UV irradiation of living cells to create a ‘zero-order’ photocrosslink between the m^7^G cap of an mRNA (Pelletier and Sonenberg, 1985) and an adjacent residue of bound eIF4E. This crosslink (Step 1, Fig. 1A) thus stably captures the *in vivo* protein:RNA interaction; subsequent purification of eIF4E is done so as to co-purify *only* photocrosslinked mRNAs. Steps 2, 3 and 5 (Fig. 1A) illustrate the critical steps of this rigorous purification process: post-crosslinking addition of limiting amounts of RNase (reducing the crosslinked RNA “tags” to fragments of ~50-80 nt), IP of eIF4E (and co-IP of both bound and crosslinked RNA), and a critical gel electrophoresis step to permit the selective recovery of only the crosslinked RNA tags.

**Figure 1.**
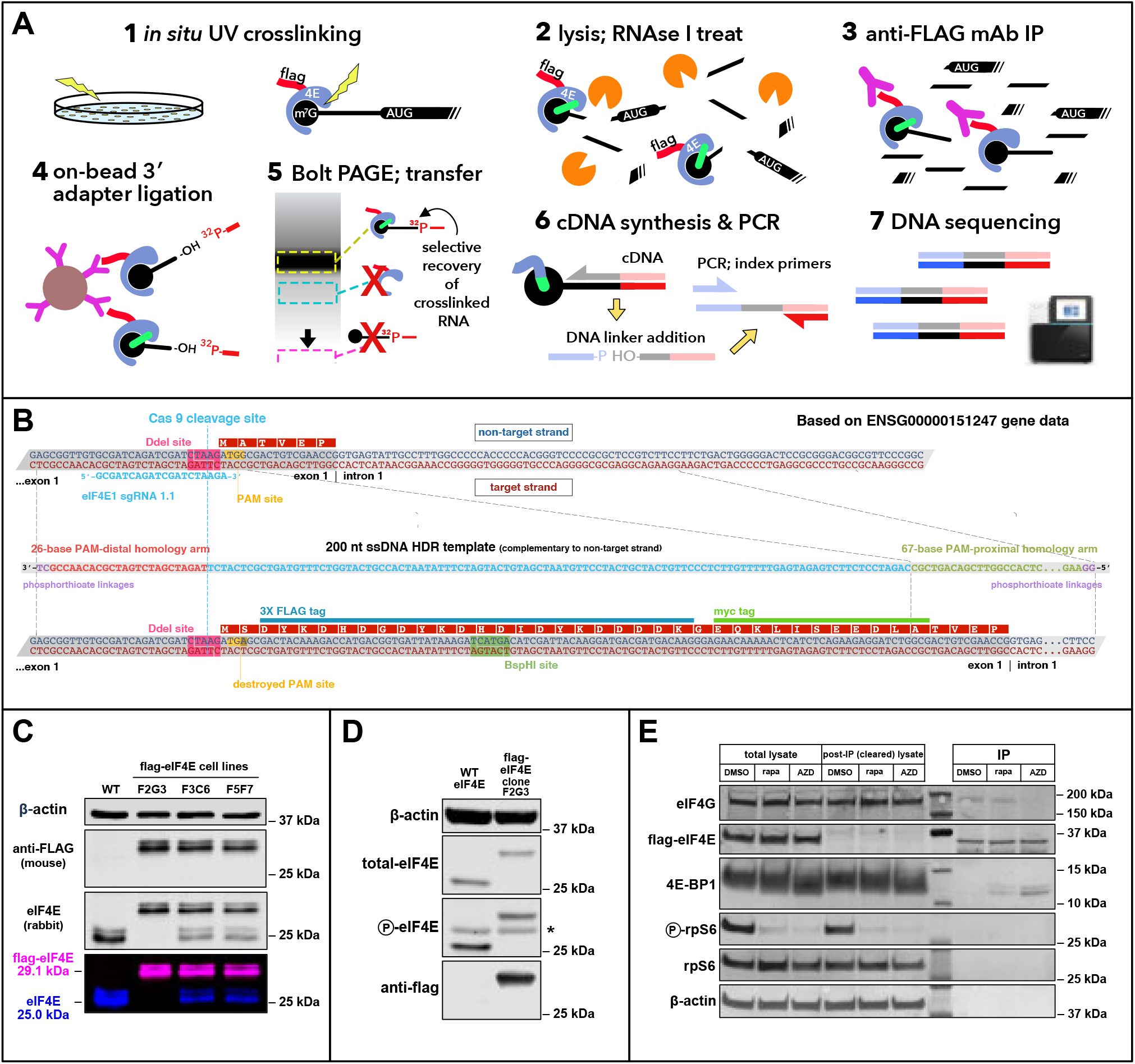
**(A)** Outline of the capCLIP method. See Star Methods for a summary of each of the 7 steps of the figure. **(B)** Scheme for addition of a 3X-flag/1X-myc epitope tag to the N-terminus of human eIF4E using CRISPR/Cas9-mediated gene editing in Hela cells (henceforth identified as ‘flag-eIF4E’). **(C)** Anti-eIF4E/anti-flag westerns of WT and three CRISPR-edited flag-eIF4E Hela cell lines. The 29 kDa flag-eIF4E and the 25 kDa WT-eIF4E bands detected by the anti-eIF4E antibody, and the 29 kDa flag-eIF4E band detected solely by anti-flag can be seen in the both the separate and merged LiCor image channels. **(D)** Anti-total-eIF4E/ P-eIF4E/ flag westerns of WT and 3X-flag-eIF4E Hela cells. The * denotes a non-specific band consistently seen with this anti-P-eIF4E antibody. **(E)** Western blot of IPs of flag-eIF4E from Hela cells using indicated antibodies. Anti-P-rpS6, and rpS6 antibodies were used to evaluate the ability of rapamycin and AZD to inhibit mTORC1 signalling. Molecular weight markers appear between the depleted and IP samples. The relative loading volumes for total lysate, cleared lysate and IP lanes is 1:1:1 (thus the amount of protein used as *input* for each IP reaction is the same as the amount of protein loaded into each total lysate lane).

CapCLIP requires efficient and specific IP of eIF4E. As there are no commercially-available anti-eIF4E antibodies that work in IP, we applied CRISPR/Cas9 genome editing to introduce a 3X-flag epitope tag at the N-termini of the chromosomal copies of the eIF4E gene in Hela cells (Figure 1B and Star Methods). Western analysis of WT Hela cells and 3 clonal lines is shown in Figure 1C. Clone F2G3, which exhibits homozygous expression of the 3X flag-eIF4E, was used for subsequent experiments; further analysis confirmed similar levels of eIF4E expression and phosphorylation (on Ser209) in WT and flag-eIF4E cells (Figure 1D).

### Using capCLIP to probe how the effects of mTORC1 inhibition on translation initiation

A major effect of mTORC1 inhibition is the impairment of translation of TOP mRNAs (Meyuhas and Kahan, 2015). While regulation of TOP mRNA translation by mTORC1 clearly occurs at initiation, there is debate about the mechanism(s) involved. Several translation-related proteins are direct or indirect substrates of mTORC1 including the eIF4E-binding proteins (4E-BPs, which comprise 4E-BP1, -BP2 and -BP3; reviewed (Proud, 2019)) and LARP1 (Fonseca et al., 2015; Hong et al., 2017). Active mTORC1 phosphorylates 4E-BPs at multiple residues; hyper-phosphorylated 4E-BPs cannot bind to eIF4E, allowing eIF4E to bind other partners, including eIF4G, and thereby mediate translation initiation (Gingras et al., 2001). While mTOR-KIs inhibit 4E-BPs phosphorylation and consequently disrupt eIF4E-eIF4G binding (Choo et al., 2008), rapamycin, which binds the immunophilin FKBP12 and indirectly impairs the kinase function of mTORC1(Aylett et al., 2016), generally has little effect on 4E-BP1 phosphorylation, and therefore on association of eIF4E with eIF4G. While inhibition of TOP mRNA translation by mTOR-KIs was thought to be mediated through 4E-BP binding of eIF4E (Thoreen et al., 2012), how TOP mRNA specificity was achieved, or why rapamycin likewise impairs TOP mRNA translation remained unclear (Huo et al., 2012). Growing evidence suggests that TOP mRNA translational inhibition is instead mediated by LARP1. Inhibition of mTORC1 by both mTOR-KIs and rapamycin promotes the hypo-phosphorylation of LARP1, which then competes with eIF4E for binding specifically to TOP mRNAs (Lahr et al., 2017; Philippe et al., 2018). Recently, ribosome profiling of WT and LARP1-deficient human cells using the mTOR-KI Torin1 revealed that LARP1 is essential for translational repression of TOP mRNAs under conditions of mTORC1 inhibition (Philippe et al., 2020).

In preparation for capCLIP, we evaluated the consequences of mTORC1 inhibition on binding of flag-eIF4E to its protein partners in our engineered Hela cells. We treated cells with 100 nM AZD8055 (an mTOR-KI), 100 nM rapamycin or DMSO alone for 2 h followed by IP of flag-eIF4E (Figure 1E) from the resulting cell lysates. Importantly, eIF4E-eIF4G binding was not affected by rapamycin, but was strongly decreased by AZD8055. These data are in agreement with our earlier findings for Hela cells (including pull-downs of endogenous eIF4E on cap resin) which showed that while rapamycin treatment only slightly alters phosphorylation of 4E-BP1 and its binding to eIF4E, AZD8055, in contrast, strongly inhibits 4E-BP1 phosphorylation, increasing binding of 4E-BP1 to eIF4E (Huo et al., 2012). We conclude that normal levels and regulation of eIF4E are preserved in our gene-edited cells.

Next, we demonstrate that flag-eIF4E is quantitatively captured by anti-flag antibody under the high-detergent and high-salt conditions used in CLIP (Figure 2A). Figure 2B demonstrates that short-wavelength UV irradiation is required for co-capture of RNA fragments in the flag-eIF4E IP and that the amount and length of RNA covalently linked to flag-eIF4E is sensitive to the concentration of RNase I. IP from lysates subject to both ‘high’ and ‘low’ RNase (0.10 or 0.02 U/μl of lysate) treatments show an appropriate ‘smear’ of signal from the ^32^P-labelled RNA on the membrane, with the greatest intensity ~37 kDa, ~8 kDa higher than the 29 kDa flag-eIF4E protein. As the low RNase lane has ~3-fold greater RNA ‘signal intensity’ that the high RNase lane, the lower concentration was used for capCLIP.

**Figure 2.**
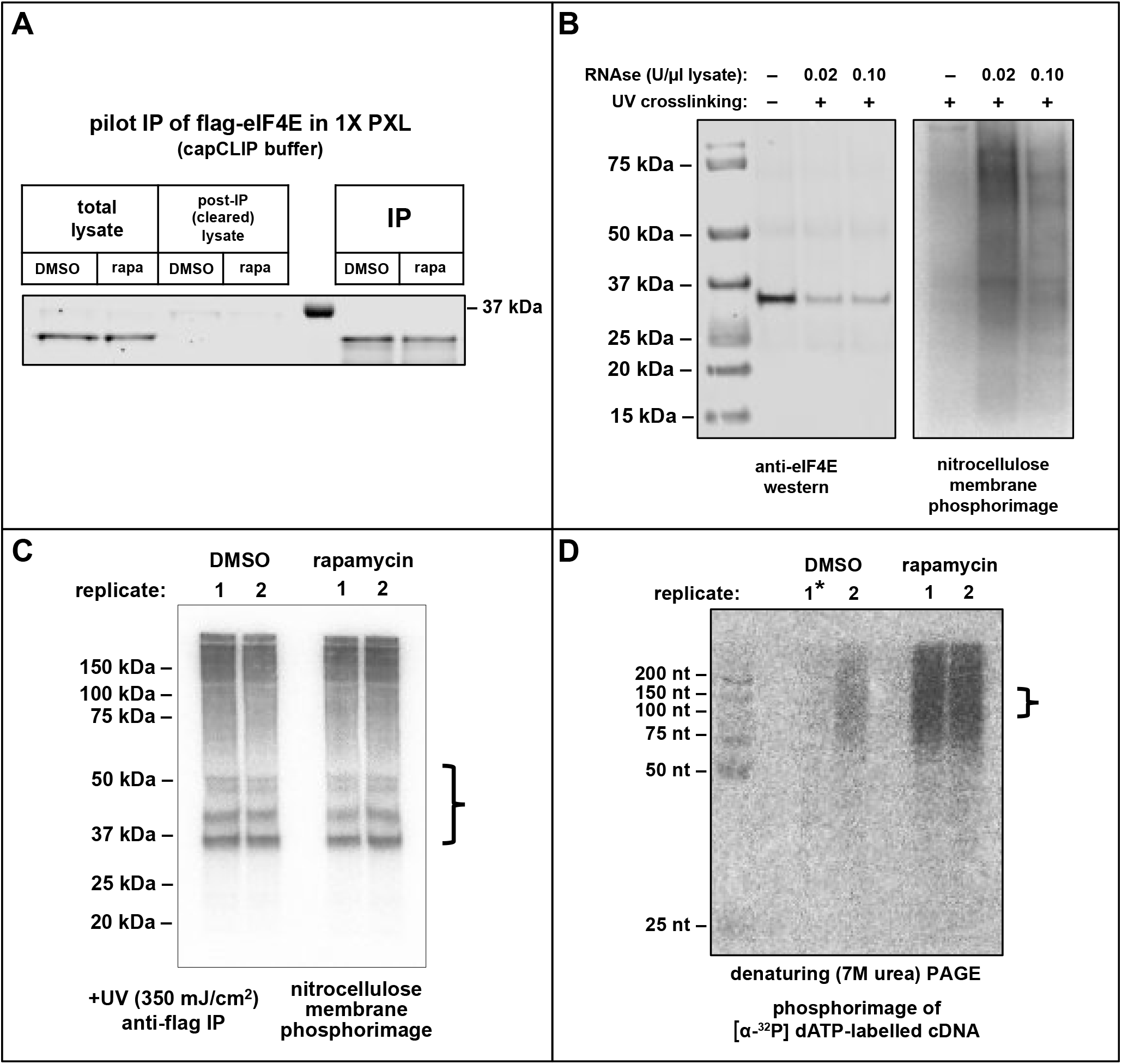
**(A)** Western analysis of flag-eIF4E IPs from untreated (DMSO) and rapamycin-treated (rapa) Hela cells. The relative loading volumes for total lysate, cleared lysate and IP lanes is 1:1:2.5. **(B)** Left: Anti-eIF4E western of flag-eIF4E (29.1 kDa) IPed from Hela cells in a pilot capCLIP experiment to test optimal RNase concentration. **(C & D)**‘Preparative’ capCLIP experiment of flag-eIF4E Hela cells ± rapamycin treatment. **(C)** Phosphorimage of ^32^P-labelled RNA after PAGE separation and transfer to nitrocellulose. Bracket at right indicates the portion of the membrane used for subsequent isolation of RNA. **(D)** Phosphorimage of ^32^P-labelled cDNA on a 7M urea denaturing PAGE. The bracket at right indicates the size range of cDNA taken for subsequent PCR library amplification.

For preparative capCLIP, flag-eIF4E Hela cells were treated for 2 h with 100 nM rapamycin (or DMSO), UV crosslinked on ice, and then processed through the remainder of the capCLIP methodology as detailed in the Star Methods. Two biological replicates per treatment group (2 rapamycin and 2 DMSO) were performed. Flag-eIF4E, along with bound/crosslinked capped-mRNA fragments, was IPed from all 4 samples. A ^32^P-labelled RNA adapter was ligated to the 3′ end of all co-IPed RNA, and the covalently-linked flag-eIF4E:RNA complexes were separated from non-crosslinked RNA using Bis-Tris PAGE, followed by transfer of the gel contents to nitrocellulose. The radioactive RNA was detected by phosphorimaging (Fig. 2C). Comparing the left and right halves of Figure 2C, there is no obvious difference in the amount of crosslinked and IPed RNA upon rapamycin treatment (identical amounts of lysate were IPed and loaded for Bis-Tris PAGE analysis). RNA from flag-eIF4E:RNA complexes migrating at approximately 37-50 kDa in Figure 2C was isolated; subsequent ^32^P-labelled cDNA synthesis products were ligated to a 3′ DNA adapter with UMI (unique molecular identifier), and separated on denaturing PAGE to select for final DNA products containing CLIP tags of ~30-80 nt in length (Fig. 2D). Indexed PCR libraries were prepared from each sample and combined for high-throughput sequencing.

### The eIF4E cap-ome provides a comprehensive picture of cellular translation

Filtered, deduplicated capCLIP tag reads were mapped to the hg19 reference genome. Peak-calling software was then used to create peak data for each replicate. (bioinformatic analysis details in Star Methods). Due to poor recovery of the DMSO replicate 1 RNA for subsequent cDNA synthesis (Fig. 2D), for some analyses two pseudo-replicate datasets were generated from the single DMSO-only control dataset and are termed controls-1A and -1B. The two rapamycin-treated replicates are named rapa-1 and rapa-2. Table 1A displays key peak statistics for the replicates. The majority of peak-calls in all replicates align within processed (spliced) mRNA sequences. The largest peak fraction in each sample (representing 34-47% of all peak calls) maps to 1^st^ exon of the mRNA (the 5′ UTR), consistent with our expectation that UV irradiation generates crosslinks between eIF4E and the m^7^GTP moiety of each mRNA. The ~2-fold higher number of 5′ UTR peaks in the rapamycin samples (compared to the DMSO replicates) is merely a consequence of splitting the single DMSO dataset into two.

**Table 1.** **(1A) rapamycin capCLIP sequencing statistics**. Distribution of peak-calls from the two control pseudo-replicate sequencing datasets and from the two rapamycin-treated replicate sequencing datasets. (**1B) eFT-508 capCLIP sequencing statistics.** Distribution of peak-calls from the two PMA-only sequencing datasets and from the two eFT-508-treated replicate sequencing datasets.

1A. rapamycin capCLIP
| eIF4E-ctrl set 1A | # of peaks | log<sub>2</sub> enrichment | % peaks |
| --- | --- | --- | --- |
| 3' UTR | 109 | 1.68 | 2.6 |
| miRNA | 11 | 5.01 | 0.3 |
| ncRNA | 60 | 2.70 | 1.4 |
| pseudo | 8 | 1.45 | 0.2 |
| exon | 585 | 3.62 | 13.9 |
| intron | 759 | -1.14 | 18.0 |
| intergenic | 610 | -2.00 | 14.4 |
| **5' UTR** | **1941** | **7.51** | **46.0** |
| snoRNA | 129 | 11.17 | 3.1 |
| snRNA | 10 | 10.29 | 0.2 |

| eIF4E-ctrl set 1B | # of peaks | log<sub>2</sub> enrichment | % peaks |
| --- | --- | --- | --- |
| 3' UTR | 121 | 1.81 | 2.8 |
| miRNA | 8 | 4.52 | 0.2 |
| ncRNA | 54 | 2.53 | 1.3 |
| pseudo | 10 | 1.75 | 0.2 |
| exon | 573 | 3.56 | 13.3 |
| intron | 806 | -1.08 | 18.8 |
| intergenic | 627 | -1.99 | 14.6 |
| **5' UTR** | **1955** | **7.49** | **45.5** |
| snoRNA | 130 | 11.15 | 3.0 |
| snRNA | 12 | 10.52 | 0.3 |

| eIF4E-rapa set 1 | # of peaks | log<sub>2</sub> enrichment | % peaks |
| --- | --- | --- | --- |
| 3' UTR | 689 | 2.71 | 5.3 |
| miRNA | 24 | 4.50 | 0.2 |
| ncRNA | 166 | 2.54 | 1.3 |
| pseudo | 38 | 2.07 | 0.3 |
| exon | 3180 | 4.43 | 24.3 |
| intron | 2578 | -1.01 | 19.7 |
| intergenic | 1815 | -2.06 | 13.8 |
| **5' UTR** | **4412** | **7.06** | **33.7** |
| snoRNA | 187 | 10.07 | 1.4 |
| snRNA | 18 | 9.50 | 0.1 |

| eIF4E-rapa set 2 | # of peaks | log<sub>2</sub> enrichment | % peaks |
| --- | --- | --- | --- |
| 3' UTR | 713 | 2.82 | 5.7 |
| miRNA | 19 | 4.23 | 0.2 |
| ncRNA | 182 | 2.74 | 1.5 |
| pseudo | 43 | 2.31 | 0.3 |
| exon | 3045 | 4.43 | 24.3 |
| intron | 2395 | -1.05 | 19.1 |
| intergenic | 1705 | -2.09 | 13.6 |
| **5' UTR** | **4193** | **7.05** | **33.5** |
| snoRNA | 194 | 10.19 | 1.6 |
| snRNA | 20 | 9.72 | 0.2 |

1B. eFT-508 capCLIP
| eIF4E-PMA set 1A | # of peaks | log<sub>2</sub> enrichment | % peaks |
| --- | --- | --- | --- |
| 3' UTR | 442 | 2.55 | 4.7 |
| miRNA | 19 | 4.64 | 0.2 |
| ncRNA | 150 | 2.87 | 1.6 |
| pseudo | 22 | 1.76 | 0.2 |
| exon | 2964 | 4.77 | 31.6 |
| intron | 1769 | -1.07 | 18.8 |
| intergenic | 1071 | -2.34 | 11.4 |
| **5' UTR** | **2764** | **7.26** | **29.4** |
| snoRNA | 170 | 10.41 | 1.8 |
| snRNA | 19 | 10.06 | 0.2 |

| eIF4E-PMA set 1B | # of peaks | log<sub>2</sub> enrichment | % peaks |
| --- | --- | --- | --- |
| 3' UTR | 433 | 2.52 | 4.6 |
| miRNA | 19 | 4.64 | 0.2 |
| ncRNA | 156 | 2.93 | 1.7 |
| pseudo | 27 | 2.06 | 0.3 |
| exon | 2891 | 4.73 | 30.9 |
| intron | 1781 | -1.06 | 19.0 |
| intergenic | 1074 | -1.06 | 19.0 |
| **5' UTR** | **2806** | **7.28** | **29.9** |
| snoRNA | 166 | 10.38 | 1.8 |
| snRNA | 17 | 9.90 | 0.2 |

| eIF4E-eFT set 1 | # of peaks | log<sub>2</sub> enrichment | % peaks |
| --- | --- | --- | --- |
| 3' UTR | 503 | 2.23 | 3.8 |
| miRNA | 16 | 3.89 | 0.1 |
| ncRNA | 198 | 2.77 | 1.5 |
| pseudo | 37 | 2.01 | 0.3% |
| exon | 3657 | 4.57 | 27.6 |
| intron | 2941 | -0.84 | 22.2 |
| intergenic | 1538 | -2.32 | 11.6 |
| **5' UTR** | **4202** | **7.36** | **31.7** |
| snoRNA | 158 | 9.81 | 1.2 |
| snRNA | 14 | 9.12 | 0.1% |

**Table 1. (1A) rapamycin capCLIP sequencing statistics.**
| eIF4E-eFT set 2 | # of peaks | log<sub>2</sub> enrichment | % peaks |
| --- | --- | --- | --- |
| 3' UTR | 562 | 2.51 | 4.6 |
| miRNA | 19 | 4.26 | 0.2 |
| ncRNA | 173 | 2.70 | 1.4 |
| pseudo | 37 | 2.13 | 0.3% |
| exon | 3193 | 4.50 | 26.1 |
| intron | 2836 | -0.77 | 23.2 |
| intergenic | 1483 | -2.25 | 12.1 |
| **5' UTR** | **3735** | **7.31** | **30.6** |
| snoRNA | 159 | 9.94 | 1.3 |
| snRNA | 17 | 9.52 | 0.1 |

PCA (principal component analysis) of the four replicates (Figure 3A) indicates that rapamycin leads to a significant and consistent change in the composition of the peak data. To measure the impact of rapamycin treatment upon eIF4E-mRNA association, and to see if these alterations reflect a mRNA’s translational activity, DiffBind (Ross-Innes et al., 2012) was used to identify those mRNAs whose association with eIF4E is significantly altered by rapamycin. In total, 5′ UTR peak data for the replicates specify the eIF4E cap-ome as a collection of 3,372 individual mRNAs. Figure 3B plots each mRNA as a function of its mRNA abundance (log_2_ of peak area; x-axis) and the log_2_ fold change of its binding to eIF4E in rapamycin- vs. DMSO-treated cells (y-axis). As a population (dashed yellow line; log_2_ fold change = 0.02) the eIF4E cap-ome exhibits essentially unchanged binding to eIF4E upon rapamycin treatment, and the same is true for the vast majority of individual mRNAs (each mRNA depicted as a grey spot). However, 86 mRNAs (red spots) do exhibit significantly altered binding to eIF4E upon rapamycin treatment (log_2_ fold-change ≶ than 0.50; *p* value < 0.01; FDR ≤ 0.05), with all 86 of them *decreasing*. About two-thirds of the 86 mRNAs were previously identified in ribosome profiling screens (Hsieh et al., 2012; Thoreen et al., 2012) as ones whose translation is impaired by mTOR-KI-mediated inhibition of mTORC1. Additional detail on these mRNAs is available in **Supplemental Table 1**.

**Figure 3.**
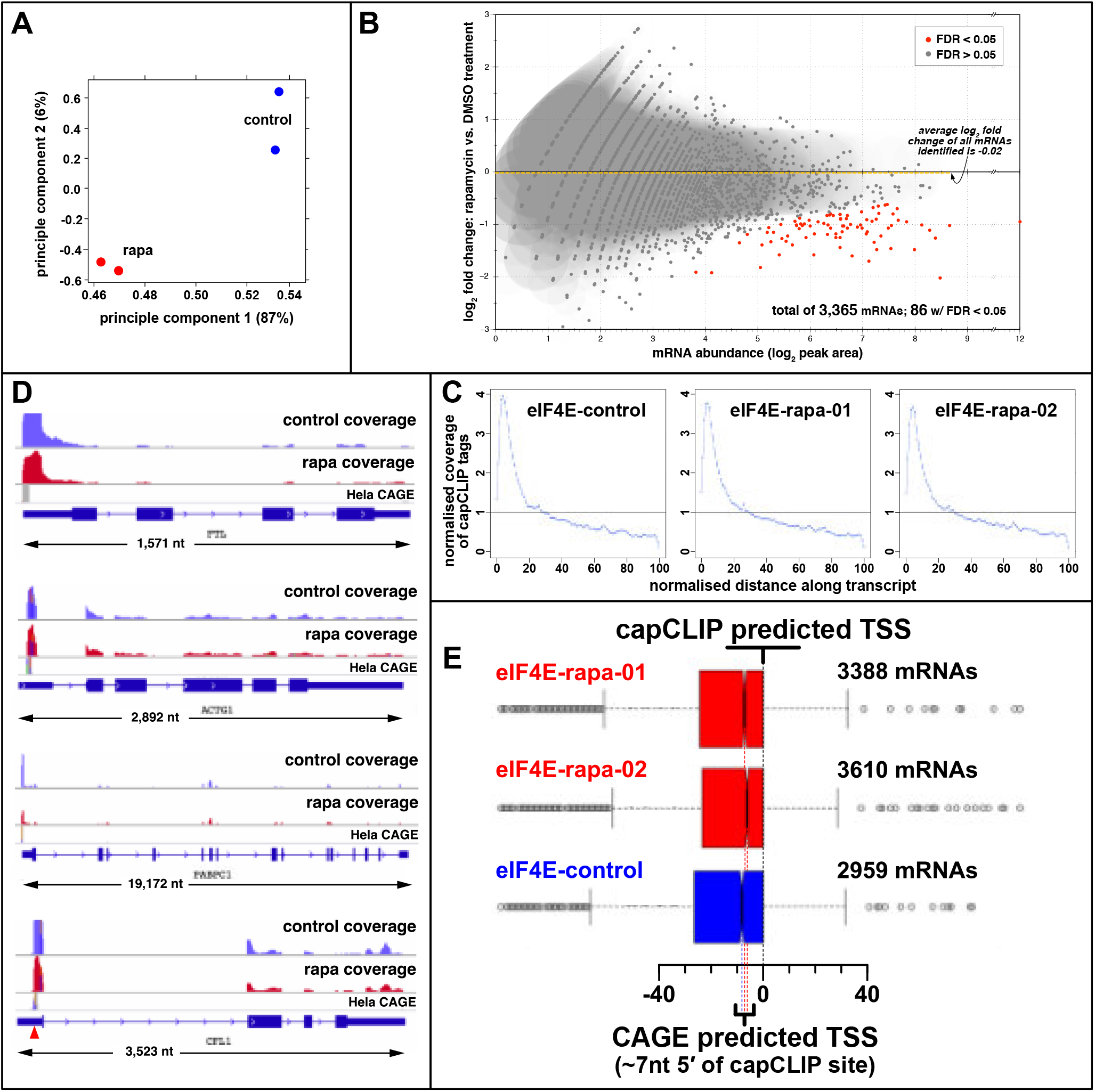
**(A)** Principal component analysis (PCA) of the two control and two rapamycin sequencing datasets. **(B)** log_2_ fold change in eIF4E affinity for 3,365 individual mRNAs vs. mRNA abundance (log_2_ of peak area). 86 mRNAs show a statistically significant log_2_ fold-change in eIF4E binding (p >0.01; FDR>0.05). Average log_2_ fold-change in eIF4E binding of the total mRNA population is −0.02. **(C)** Distribution of capCLIP tags as a function of the normalised distance along the transcript for the single, unsplit control (DMSO) replicate (see text) and two rapamycin (rapa) replicates. **(D)** IGV gene-level views, reformatted to run 5′ to 3′, left to right, to aid visualisation. Control (blue) and rapamycin (red) capCLIP tag alignments, and Hela CAGE data (grey) shown above each gene. **(E)** Comparison of transcription start sites (TSSs) predictions using either Hela CAGE or capCLIP peak-call data. The box-whisker plot data is aligned using the capCLIP peak data-predicted TSSs, which begin at the 0 position on the scale below. The two red (rapa) and one blue (control) descending dashed lines denote the calculated median TSSs predicted by Hela CAGE data.

In Figure 3C, capCLIP tag coverage from the single control and two rapamycin sequence datasets for all tags aligning to processed (spliced) mRNA is plotted over a normalised mRNA ‘length-space’; all 3 plots show the robust enrichment of tags at the extreme 5′ end of the mRNA, as suggested by the mapping data in Table 1. Figure 3D shows capCLIP tag coverage on gene-level IGV (Integrative Genomics Viewer (Robinson et al., 2011)) representations for *FTL*, *PABPC1*, *CFL1* and *ACTG1* mRNAs, along with histograms of the control and rapamycin-treated capCLIP tags data mapping to them. These views again clearly show that the vast majority of capCLIP tags which map to a particular gene align at the extreme 5′ end of its transcript. Hela CAGE data (cap analysis of gene expression (Kodzius et al., 2006; Takahashi et al., 2012)) is also shown for each mRNA in Figure 3D. While close inspection of the alignment between each gene’s capCLIP tag data and the Hela CAGE data is difficult at the scale of the figure, the 5′ boundary of each gene’s capCLIP tag data aligns with the CAGE-identified TSS for each transcript.

### Mapping TSS sites with capCLIP and CAGE data

We next systematically plotted both the 5′ boundary of each capCLIP peak and the TSS as determined from Hela CAGE data (Figure 3E) for each mRNA of the eIF4E cap-ome. We find that the capCLIP *peak* data consistently predicts a TSS site for each mRNA that is 7-8 nucleotides 3′ of the ‘authentic’ TSS site identified by CAGE. This discrepancy is almost certainly driven by a 7-8 nucleotide-long ‘decay’ in capCLIP *tag* coverage at the 5′ end of each transcript, which can be seen in the nucleotide-level IGV images of Figure 4D, which depict six individual mRNA 5′ ends. This loss of tag coverage is likely due to obstruction of the reverse transcriptase as it approaches the cap structure during synthesis of cDNA copies of each original mRNA fragment (the m^7^G cap likely still remains crosslinked to a small eIF4E-derived peptide when the purified RNA tag is treated with protease to digest away eIF4E). This missing tag coverage at the 5′ end of the mRNA leads the MACS2 peak-caller software to consistently define the 5′ boundary of mRNA cap peaks 7-8 nucleotides downstream of the authentic 5′ end of the transcript (as measured by the Hela CAGE data). The consistency of this 7-8 nt offset across the 5′ UTR peaks of the eIF4E cap-ome (clearly apparent in Figure 3E) suggests that, by compensating for this characteristic mis-location of the 5′ capCLIP peak boundary, capCLIP can predict an mRNA’s TSS independently of the TSS estimate provided by CAGE. Thus, *together* capCLIP and CAGE allow us to attain a significantly greater level of confidence in the active TSS site (or sites) used in a particular cell or tissue than either method alone. Most importantly, this high-confidence TSS prediction can be exploited to determine the relationship between a mRNA’s 5′ end sequence and the degree of loss of association with eIF4E elicited by rapamycin. Such an analysis was possible before the development of capCLIP.

**Figure 4.**
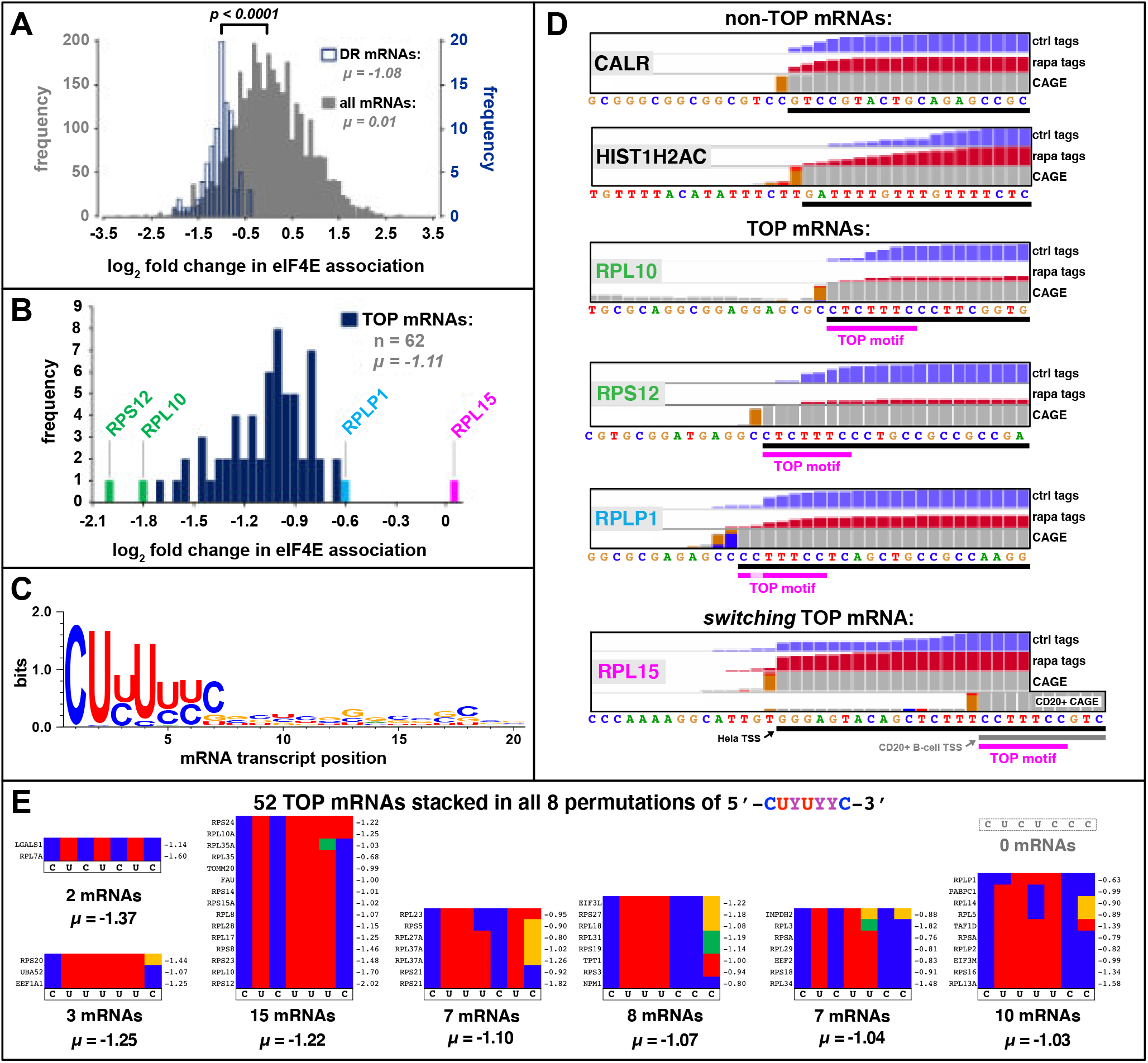
**(A)** Frequency plot of the log_2_ fold-change in eIF4E association upon treatment with rapamycin. All peak-called mRNAs (grey) and the 86 mRNAs (blue) which show a statistically significant change with drug (fold change <−0.5 or >0.5; FDR<0.05; p<0.01). **(B)** log_2_ fold-change in eIF4E association for the 62 TOP mRNAs identified by capCLIP. **(C)** Weblogo analysis of the first 20 nucleotides of 52 capCLIP-identified TOP mRNAs. **(D)** IGV plots of the extreme mRNA 5′ ends of six mRNAs; views are reformatted to run 5′ to 3′, left to right, to aid visualisation. The mRNA (black bar) is depicted below the sequence (the 5′-most nucleotide is the TSS). Normalised histogram plots of control (blue) and rapamycin treated (red) capCLIP tag data, along with Hela CAGE data (grey & orange) are shown above the sequence. In general, the first grey bar 5′ of an orange bar in the CAGE histogram denotes the TSS (Kodzius et al., 2006). Below each TOP mRNA is the 7-nt TOP consensus motif (magenta). **(E)** Sorting of the 52 TOP mRNAs into 7 of 8 possible sequence permutations of the 7-nucleotide TOP consensus motif. Individual sequences are depicted as coloured strips (red=U, blue=C, green=A and yellow=G). Gene name shown left of each strip; log_2_ fold change value (μ) upon rapamycin treatment shown at right.

### TOP mRNAs are specifically depleted from the eIF4E cap-ome with rapamycin treatment

Figure 4A shows a frequency plot depicting the 86 mRNAs whose eIF4E association is significantly altered by rapamycin (which we termed ‘differentially regulated mRNAs’ or ‘DR RNAs’, in blue) and the set of all mRNAs identified by capCLIP (in grey), arranged according to their log_2_ fold-change in eIF4E association in response to rapamycin. While the overall shift in eIF4E association of the total population of mRNAs is essentially 0 (μ = 0.01), there is an average two-fold decrease in eIF4E association amongst the DR mRNAs (μ = −1.08; p<0.0001). Using both capCLIP and CAGE data to generate high-confidence predictions of the TSS for the 86 statistically significant mRNAs, we conclude that 62 of the 86 mRNAs (72%) possess TOPs (an initial C, followed by 5-14 pyrimidines) (Meyuhas and Kahan, 2015); an additional 13 (15%) appear to have multiple TSSs, including a TOP TSS and at least one non-TOP TSS. The remaining 11 of the 86 mRNAs (13%) lack *any* identifiable TOP motif according to our capCLIP/CAGE TSS predictions. While the majority of our 62 TOP mRNAs encode, as expected (Iadevaia et al., 2008), ribosomal proteins or other components of the translational machinery, we identified two novel TOP mRNAs, *TAF1D*, and *LGALS1*. *TAF1D* is intriguing, as it is an essential component of the SL1 complex that regulates pol I-catalysed transcription of the major ribosomal RNAs (Gorski et al., 2007), which has long been known to be driven by mTORC1, and would provide a further way in which mTORC1 signalling promotes ribosome biogenesis (Iadevaia et al., 2014). *LGALS1* (galectin-1) is a secreted lectin with potential roles in apoptosis and cell growth; we are not aware of any link between *LGALS1* and mTORC1 signalling.

Overall, the data clearly support our original hypothesis that capCLIP should provide a sensitive and robust measurement of translational activity. The *specificity* of the changes and the *magnitude* of the effects of rapamycin that we see in the eIF4E cap-ome are strikingly similar to those obtained from ribosome-profiling experiments measuring the impact of the mTOR-KIs PP242 and Torin1 on translational efficiency (Hsieh et al., 2012; Thoreen et al., 2012). As such sucrose gradient centrifugation techniques to assess association of mRNAs with ribosomes in polysomes are confounded by association of mRNAs with other high-molecular weight, but non-translating, complexes, or with translationally-stalled ribosomes (Thermann and Hentze, 2007), capCLIP has the potential to substitute for, or complement, ribosome profiling methods; it will be interesting to test the two methods side-by-side.

### mRNAs sensitive to rapamycin possess a strikingly well-defined TOP motif

In Figure 4B the 62 TOP mRNAs are displayed according to their log_2_ change in association with eIF4E under rapamycin treatment. While the TOP mRNA population shifts significantly upon mTORC1 inhibition, there are significant differences in the magnitude of the response of individual mRNAs. We next asked if further analysis of these mRNAs could yield a deeper understanding of the relationship between TOP motif sequence and TOP ‘strength;’ i.e., the size of the decrease shown by a TOP mRNA in the eIF4E cap-ome from rapamycin-treated cells. Ten of the 62 mRNAs were excluded, as we were unable to use CAGE and capCLIP data to unambiguously identify the their TSSs (CAGE data for the 10 is uncharacteristically broad and two or more (usually overlapping) TSSs may be used).

We extracted the first 20 nt of each of the 52 TOP mRNAs and input this sequence data into **weblogo3** (Crooks et al., 2004). The resulting consensus sequence, which in essence defines the ‘functional’ TOP motif as defined by capCLIP, is shown in Figure 4C. While the weblogo consensus is consistent with earlier work demonstrating that functional TOP mRNAs harbor an invariant C base at position one of their transcripts, the analysis also surprisingly suggest that optimal TOP motifs exhibit additional sequence properties. Firstly, and unexpectedly, there appears to be a near-essential requirement for U in position 2; indeed, only one of the 52 functional TOP mRNAs lacks a U here. That mRNA, *RPLP1*, has C at this position (Figure 4D) and, interestingly, out of the 52 TOP mRNAs showing a significant drop in eIF4E association, it is the TOP mRNA *least* affected by rapamycin (μ = −0.63, Figure 4B). Our analysis also indicate that functional TOPs also show a very strong preference for U in position 4 and a significant preference for C in position 7. Secondly, TOP motifs are (at most) 7 nucleotides long and must be positioned directly at the mRNA’s 5′-end, with absolutely no preference for pyrimidines in positions 8-20. Thus, the capCLIP data suggest that optimal TOP motifs are defined by the sequence 5′-CUYUYYC-3′; this 7-nt motif is significantly shorter than previously suggested (Meyuhas and Kahan, 2015) (Figure 4C).

A graphical summary of our TOP motif analysis can be seen in the Figure 4D, which shows nucleotide-level views of the capCLIP data for six selected mRNAs: two ‘non-TOP’ mRNAs, *CALR* & *HIST1H2AC*; three TOP mRNAs, *RPL10*, *RPS12* & *RPLP1*; and one so-called *switching*-TOP mRNA (see below), *RPL15* (Figure 4D). The specific effect of rapamycin on TOP mRNAs can been seen by comparing the relative heights of the normalised control and rapamycin capCLIP tag histograms for each mRNA. Interestingly, the two mRNAs whose association with eIF4E is the most strongly affected by rapamycin, *RPL10* and *RPS12* (see Figure 4B), harbor identical 7-nucleotide TOP motifs: 5′-CUCUUUC-3′. As this sequence is 1 of 8 possible individual permutations of the our TOP consensus 5′-CUYUYYC-3′, we sorted all 52 TOP mRNAs by permutation group to ascertain if any individual permutation possesses unique properties (Figure 4E). Interestingly, the 5′-CUCUUUC-3′ group, containing the highest responding *RPL10* and *RPS12* mRNAs, is the most abundant permutation group by a significant margin, accommodating 15 of the 52 TOP mRNAs (29%). There is no significant difference in each group’s mean log_2_ fold change in eIF4E binding with rapamycin treatment, suggesting that each of the 8 permutations are equally functional as TOP motifs. Indeed, Figure 4E strikingly illustrates that the only permissible nucleotide alteration to the TOP consensus motif (with just 2 exceptions) is in the 7^th^ and last position of the motif. No TOP mRNAs lie in the 5′-CUCUCCC-3′ group, suggesting that this sequence may not function as a TOP, or be subject to other negative selection pressures. The correlation between an mRNA’s extent of homology with our TOP consensus motif 5′-CUYUYYC-3′ and the degree to which rapamycin alters its binding to eIF4E give us confidence that the weblog analysis does indeed identify salient features of TOP motifs.

The last mRNA in Figure 4D, *RPL15*, encodes a core protein of the 60S ribosomal subunit, and would thus be expected to behave as a TOP. However, it does not, with essentially no change in eIF4E association upon rapamycin treatment (μ = 0.03; Figure 4B). Examination of the capCLIP tag and CAGE data for *RPL15* indicates the TSS begins 5′-GGGAGTA…-3′, i.e., a non-TOP sequence. However, a search of human ENCODE CAGE data reveals that CD20+ B-lymphocytes utilise a TSS which begins 16 nt 5′ of the Hela TSS; in these cells the first 7nt of the mRNA is 5′-CCUUUCC-3′ (Figure 4D). As CD20+ B-cells transcribe *RPL15* with a TOP, we term *RPL15* a *switching*-TOP mRNA. Alternative TSS usage would permit cells to find-tune which mRNAs are subject to mTORC1-mediated translational regulation.

A consensus is now emerging that mTORC1 regulates TOP mRNA translation through the cap- and TOP mRNA-specific binding ability of LARP1, and this hypothesis is further strengthened by recent ribosome profiling data demonstrating that LARP1 is essential for translational repression of TOP mRNAs in response to mTOR inhibition (Philippe et al., 2020). As capCLIP serves as a *direct* readout of the change in eIF4E association for mRNAs (rather than inferring the sequestration of mRNAs from eIF4E, e.g., by LARP1, as ribosome profiling does), and as we find that TOP mRNAs are selectively lost from the eIF4E cap-ome through a 4E-BP-independent mechanism, our results are wholly consistent with the LARP1 TOP mRNA regulatory model. Interestingly, a LARP1:RNA 3D structure, incorporating the 8-nucleotide synthetic TOP motif 5′-CUUUUCCG-3′ surprisingly found that the 8^th^ nucleotide of the RNA did not contact the LARP1 protein fragment which bound the first 7 nucleotides of the TOP motif, but instead interacted with a second LARP1 complex in an adjacent unit cell (Lahr et al., 2017). This too is consistent both with our determination of a 7-nucleotide functional TOP motif, and with a recent study showing that LARP1 is insensitive to the nucleotides at position 8 and beyond (Philippe et al., 2020).

### TOP mRNA discovery: benchmarking of capCLIP to ribosome profiling

There is striking similarity between the set of rapamycin-responsive mRNAs identified by capCLIP and findings from previous work identifying mRNAs whose translation is regulated by mTORC1 signalling (Hsieh et al., 2012; Huo et al., 2012; Thoreen et al., 2012). This overlap strongly suggests that changes in an mRNA’s binding to eIF4E, the ‘output’ of capCLIP, is a reliable substitute for methods that more directly assess an mRNA’s translational activity (at least for mRNAs regulated by mTORC1 signalling). To more rigorously probe this hypothesis, we compared our data to the pioneering ribosome profiling studies that provided the first transcriptome-wide identification of mRNAs translationally regulated by mTOR signalling (Hsieh et al., 2012; Thoreen et al., 2012). Ribosome profiling data is most often employed to compute a mRNA’s translational efficiency (T_E_), which relates the number of ribosomes occupying a mRNAs open reading-frame (ORF) to the mRNA’s expression level (Ingolia, 2016). The Hsieh and Thoreen studies both showed that mTOR-KIs cause statistically significant reductions in T_E_ (ranging in general from 2-4 fold) for the ~90 known TOP mRNAs. Hsieh *et al.* also report T_E_s for individual TOP mRNAs upon rapamycin treatment (average T_E_ reduction of ~1.8-fold); however, only ~20% of these appear to have an FDR value less than 0.05 (and these 20% are not identified).

Nevertheless, we sought to determine the degree of correlation between the changes in eIF4E binding (capCLIP) and the changes in T_E_ (ribosome profiling) upon rapamycin treatment. Figure 5A plots the log_2_ fold change in eIF4E-cap binding from capCLIP vs. the log_2_ fold change in T_E_ for the 45 TOP mRNAs common to our differentially-regulated capCLIP target list (p<0.01; FDR<0.05) and the rapamycin ribosomal profiling T_E_ data of Hsieh *et al.* (FDR and p-values unknown). A Spearman *r-value* calculation shows a significant correlation (r = 0.3216; p = 0.031) between eIF4E binding and T_E_. This correlation is certainly encouraging, though we note that direct and comprehensive comparisons of the two methods (identical cell lines/tissues, identical conditions) will be necessary to fully validate capCLIP as a quantitative indicator of translational activity.

**Figure 5.**
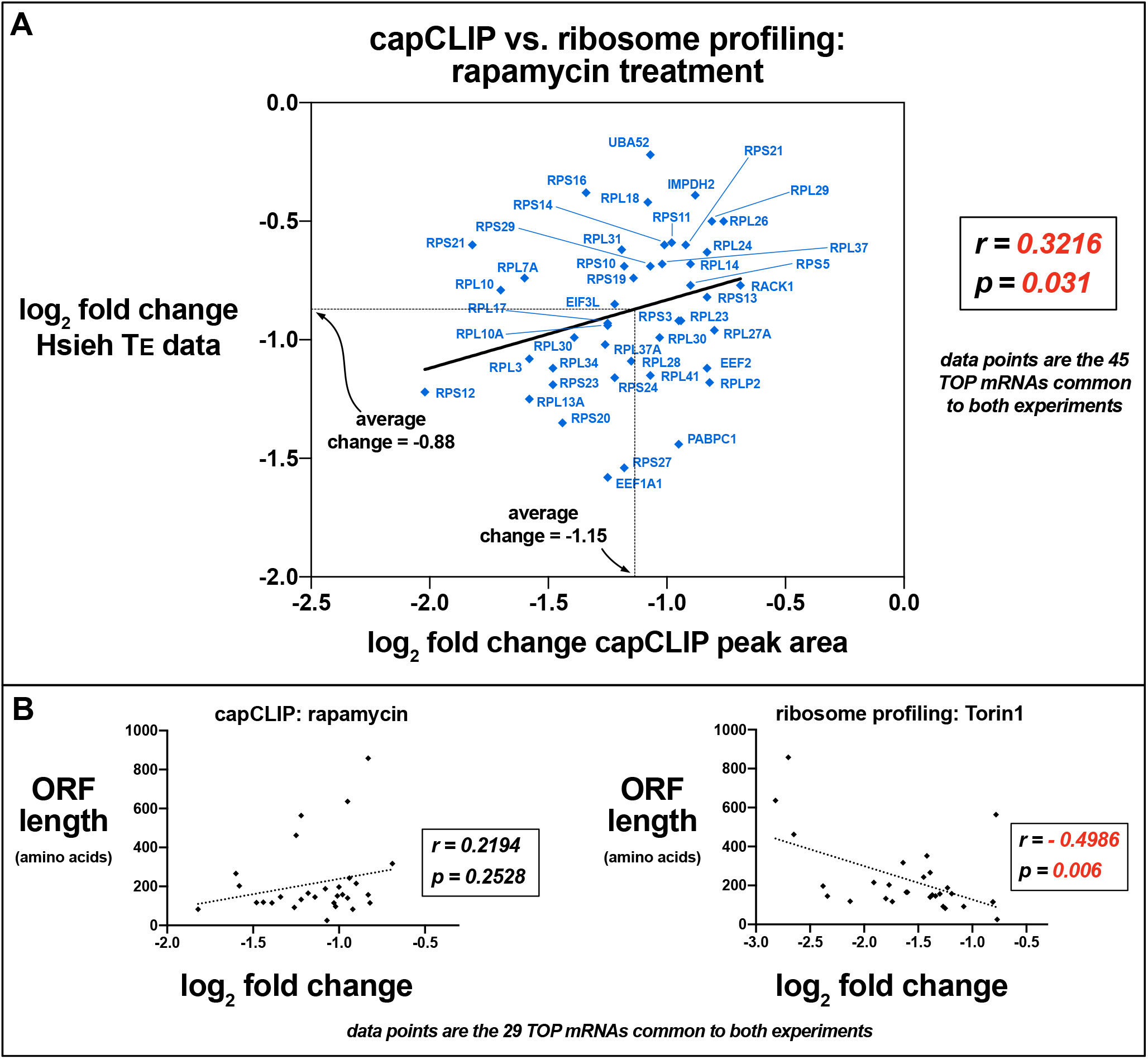
**(A)** Comparison of the 45 TOP mRNAs identified by both capCLIP and the Hsieh *et al.* ribosome profiling experiments, with log_2_ fold change of eIF4E binding (peak area) *vs.* log_2_ fold change in T_E_. **(B) Left:** log_2_ fold change of eIF4E binding *vs*. ORF length for 29 TOP mRNAs from capCLIP and Thoreen *et al.* ribosome profiling data (see text). **Right:** log_2_ fold change of T_E_ ±Torin1 treatment *vs.* ORF length for the same 29 mRNAs.

As both the degree of translational repression and the absolute number of mRNA targets repressed are generally much larger with mTOR-KIs than with rapamycin, we did not attempt a similar comparison of our capCLIP data with either study’s mTOR-KI T_E_ data. However, we noted that a number of TOP mRNAs categorised by these groups’ ribosome profiling experiments as profoundly sensitive to mTORC1 inhibition were not so in the rapamycin capCLIP data. Examination of these mRNAs revealed that most possess significantly longer primary open reading-frames (ORFs) than the majority of TOP mRNAs. As accurate calculations of T_E_ across mRNAs whose ORF lengths differ by ≥5-fold may be difficult to achieve (Ingolia, 2016), we tested for a relationship between ORF length and the degree of translational repression. In Figure 5B, the log_2_ fold change in eIF4E binding or the log_2_ fold change of T_E_ are plotted against mRNA ORF length for the 29 TOP mRNAs common to the rapamycin capCLIP and Torin1 ribosome profiling dataset of Thoreen *et al.* No significant relationship was found for ORF length and eIF4E binding (r = 0.2194; p = 0.253), but there is a significant negative correlation between ORF length and T_E_ (r = −0.4986; p = 0.006). We are unaware of any rigorous studies that have experimentally tested how ORF length impacts T_E_ calculations using ribosome profiling data. However, as ribosome footprint densities likely vary along any ORF, the uncertainty in determining the ‘correct’ ribosome density will tend to rise with ORF length. As capCLIP is not influenced by ORF length, it may, particularly when combined with RNA-seq data to assess changes in steady-state mRNA levels, provide a highly accurate picture of cap-dependent translation.

### capCLIP reveals significant differences in the eIF4E and P-eIF4E ‘cap-omes’

As our initial capCLIP study demonstrates the utility of the method for probing translational activity, we next sought to use the technique to address a long-standing translational mystery: the biological role played by phosphorylation of eIF4E by MNKs. Surprisingly, even after >30 years of work, the functional consequences of phosphorylation of eIF4E on Ser209 remain obscure. The flag-eIF4E Hela cell line, like WT cells, exhibits moderate levels of P-eIF4E under normal growth conditions. Serum starvation for 12 h reduced P-eIF4E levels to well below basal levels (lane 1, Figure 6A). Serum-starved cells treated with phorbol myristate acetate (PMA), a potent activator of ERK signalling and thus of MNK activity, promoted robust phosphorylation of eIF4E (Wang et al., 1998). This increase in P-eIF4E was seen as early as 15 min. post-PMA treatment (lanes 2-6, Figure 6A). To obtain cells with undetectable P-eIF4E to serve as a control, we used the highly-specific MNK inhibitor eFT-508 (Reich et al., 2018). A dose-response experiment (Figure 6B). showed that the lowest dose of eFT-508 to substantially inhibit eIF4E phosphorylation was 0.1 μM.

**Figure 6.**
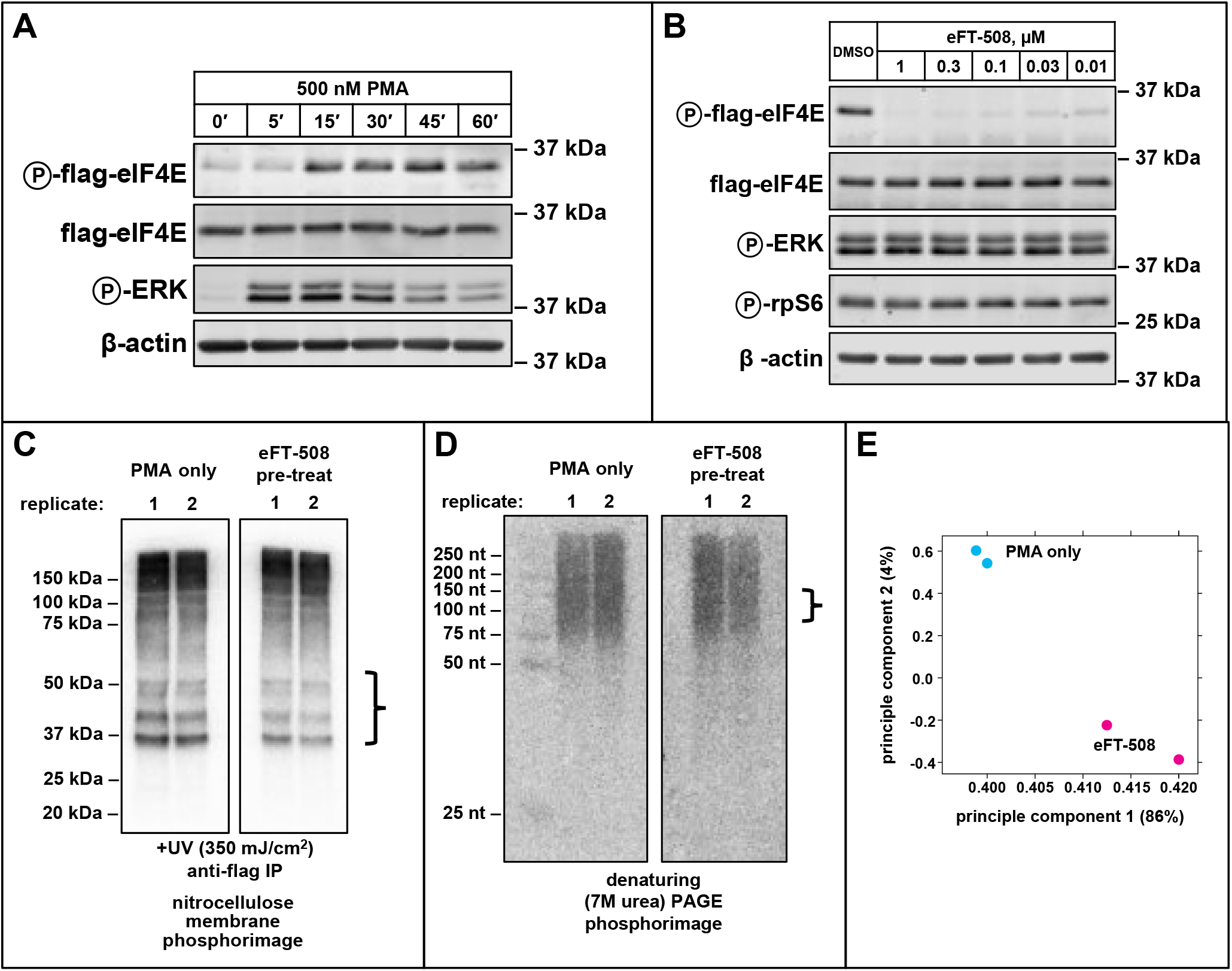
**(A)** Western analysis of P-flag-eIF4E stimulation in flag-eIF4E Hela cells. **(B)** eFT-508 dose-response of P-eIF4E levels in flag-eIF4E Hela cells. Continued activation of MAPK/ERK signalling was confirmed with anti-P-ERK; mTORC1 activity assayed with anti-P-rps6. **(C & D)**‘Preparative’ capCLIP experiment of flag-eIF4E Hela cells ±eFT-508-mediated inhibition of eIF4E phosphorylation. **(C)** Phosphorimage of ^32^P-labelled RNA after PAGE separation and transfer to nitrocellulose. Bracket at right indicates the portion of the membrane used for subsequent isolation of RNA. **(D)** Phosphorimage of ^32^P-labelled cDNA on a 7M urea denaturing PAGE. The bracket at right indicates the size range of cDNA taken for subsequent PCR library amplification. **(E)** PCA of the 2 control (PMA-only) DNA sequencing datasets and 2 eFT-508 pre-treated (eFT-508) DNA sequencing datasets.

For capCLIP, serum-starved flag-eIF4E Hela cells were treated with 0.1 μM eFT-508 or DMSO only for 1 h, followed by 30 min treatment with 500 nM PMA (2 replicates for each treatment). Following an essentially identical work-up to that of the rapamycin capCLIP study, ^32^P-labelled covalently-linked flag-eIF4E:RNA complexes were separated from non-crosslinked RNA using Bis-Tris PAGE, followed by transfer of the gel contents to nitrocellulose (Figure 6C). RNA from flag-eIF4E:RNA complexes was isolated and used for cDNA synthesis; ^32^P-labelled cDNA products were ligated to a 3′ DNA adapter with UMI, and separated on denaturing PAGE to select for final DNA products containing CLIP tags of ~30-80 nt in length (Figure 6D). Table 1B displays key peak statistics. Approximately 30% of all peak-calls in the 4 replicates map to 1^st^ exon of the mRNA (the 5′ UTR). PCA of the 4 datasets (Figure 6E) indicates that the PMA-only and eFT-508 treatments are clearly differentiable on the two components of the analysis.

Of the 4,965 mRNAs identified by peak-calling in this capCLIP experiment, 256 exhibited statistically-significant differences in eIF4E binding upon eFT-508 treatment (FDR<0.05). Additional detail on these 256 mRNAs is provided in **Supplemental Table 2**. Figure 7A plots each mRNA’s abundance vs. the change in eIF4E binding upon eIF4E (de)phosphorylation (left y-axis). The right y-axis depicts the same data but with respect to the underlying PMA/eFT-508 treatment paradigm, and, hence, has the opposite sign. Surprisingly, all 256 of these statistically-significant changes are *reductions* in eIF4E binding upon stimulating its phosphorylation. Furthermore, upon stimulation of phosphorylation the 4,965 mRNA population *as a whole* exhibits a reduction in eIF4E binding upon stimulation, with a log_2_ fold change in eIF4E affinity of −0.75 (Figure 7A, dashed yellow line). This population behaviour is significantly different than what was seen with rapamycin treatment, where mRNA population as a whole was essentially unchanged by the inhibitor treatment. The plot in Figure 7A also shows that a small fraction of the cap-ome mRNAs display an *increase* in binding to eIF4E upon its phosphorylation, although none of the mRNAs within this ‘increasing’ subpopulation exhibits statistically significant differences between the two treatment conditions; greater capCLIP library depth will be required to resolve the significance of this fraction.

**Figure 7.**
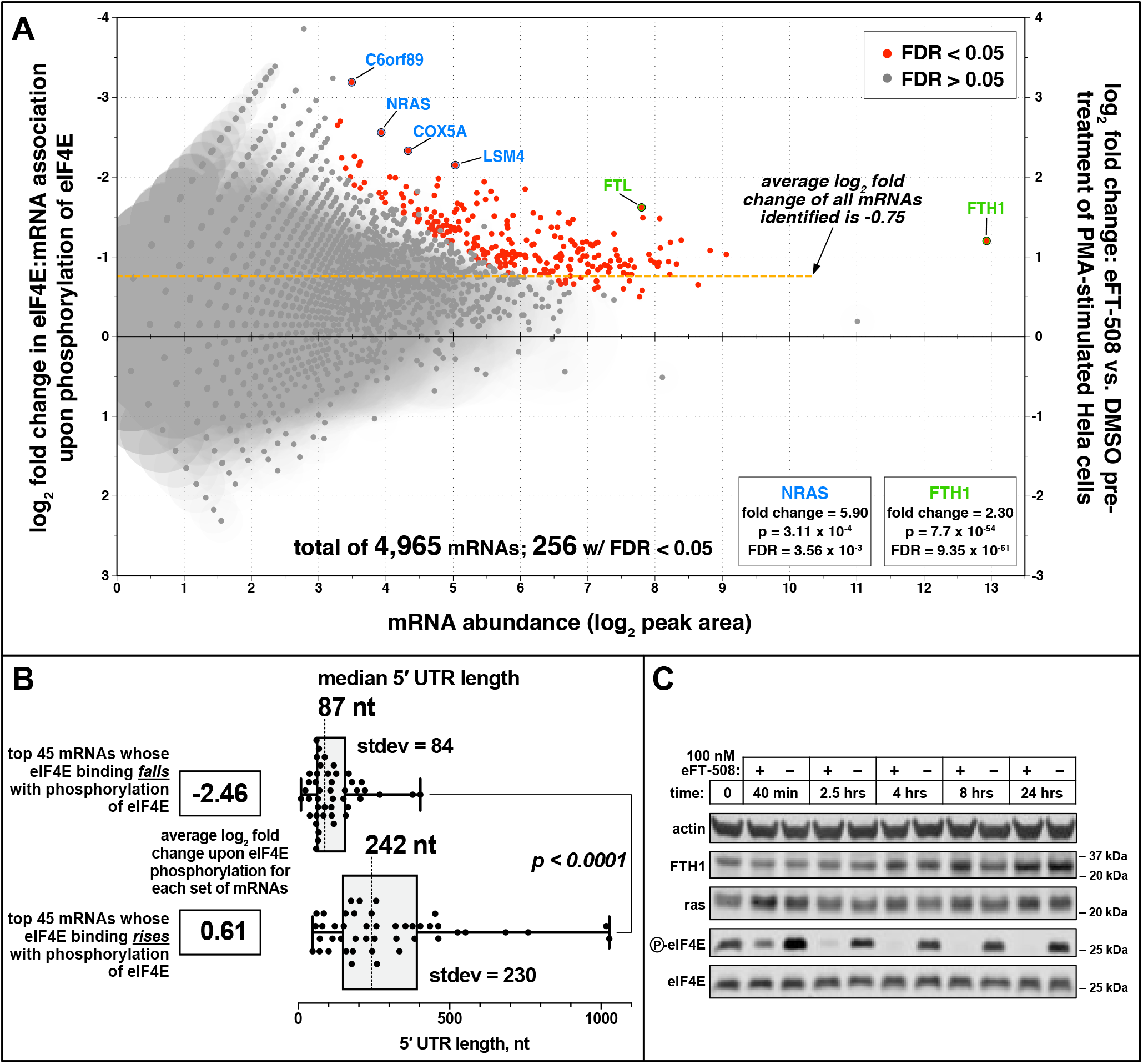
**(A)** Plot of mRNA abundance (x-axis; log_2_ of the capCLIP peak area) vs. the log_2_ fold change in eIF4E binding upon eIF4E phosphorylation (left y-axis). Right y-axis expresses the results with respect to the underlying PMA/eFT-508 treatment paradigm. Total of 4,965 mRNAs; 256 exhibit a significant change in log_2_ fold change binding to eIF4E upon phosphorylation (p >0.01; FDR>0.05). **(B)** 5′ UTR length distribution for the 45 mRNAs with the greatest *reduction* (top) or the greatest apparent *increase* (bottom) in eIF4E binding upon eIF4E phosphorylation. **(C)** Western blot of protein levels in WT Hela cells treated with DMSO only or 100 nM eFT-508 for indicated times. Inhibition of eIF4E phosphorylation by eFT-508 was monitored with anti-P-eIF4E antibody.

Our data therefore indicate that overall, *phosphorylation decreases the intrinsic affinity of eIF4E for capped mRNAs*. This finding is intriguing, as the only unbiased screen for effects of eIF4E phosphorylation (Furic et al., 2010) concluded that phosphorylation of eIF4E *promoted* translation of certain mRNAs. For their study, Furic *et al.* substituted Ser209 in eIF4E with a non-phosphorylatable alanine in mouse embryonic fibroblasts, and employed sucrose density gradient centrifugation to identify polysome-associated mRNAs by microarray. As this substitution may not accurately mirror the absence of a phosphate at this position (Beggs et al., 2015) and, as the study did not directly analyse translational activity, or crucially, eIF4E-mRNA interactions, it is difficult to compare those data with our findings. Notably, our capCLIP data is wholly consistent with *in vitro* biophysical studies demonstrating that phosphorylation of eIF4E lowers its affinity for the cap (Scheper et al., 2002; Slepenkov et al., 2006; Zuberek et al., 2003). Phosphorylation of eIF4E may do so due to electrostatic repulsion between the negatively-charged phosphate on Ser209 and similarly charged groups on the first nucleotides of the bound mRNA.

We next asked if we could identify characteristics (primarily length and base composition) that are shared by the 5′ UTRs of mRNAs which exhibit the largest reductions in eIF4E binding upon phosphorylation, and which differentiate these mRNAs from those whose eIF4E association appears to rise with eIF4E phosphorylation. In total, 90 mRNAs were selected: the 45 with the greatest *reduction* in eIF4E binding upon eIF4E phosphorylation and the 45 with the greatest (relative) *increase* in eIF4E binding upon eIF4E phosphorylation. 5′ UTR sequences were sourced from Ensembl Biomart and the NCBI RefSeq collection for all 90 mRNAs. We found a striking difference in the median 5′ UTR length for the two sets of mRNAs (Figure 7B). The 45 mRNAs whose binding to eIF4E falls with eIF4E phosphorylation have a median 5′ UTR length of 87 nt (SD=84nt), significantly shorter than the average for human 5′ UTRs (estimated to be 218 nt (NCBI RefSeq data, (Leppek et al., 2018)) or 293 nt (SD=27) (UTRdb, (Grillo et al., 2010)). Strikingly, the average log_2_ fold decrease in eIF4E binding upon phosphorylation of these 45 mRNAs is 2.46. In contrast, the median 5′ UTR length of the 45 mRNAs whose binding to eIF4E rises with eIF4E phosphorylation is 242 nt (SD=230). Notably, and in reference to the discussion above of our uncertainty regarding their apparent increase in binding to P-eIF4E, the average log_2_ fold increase in eIF4E binding of this group of mRNAs is an very modest 0.61. We also examined nucleotide composition for the two groups of 5’ UTRs. While average GC content for human 5’ UTRs is 58% (UTRdb, (Grillo et al., 2010)), it is 68% for the mRNAs which lose binding to eIF4E upon phosphorylation, and 58% for the group which gain binding. In sum, mRNAs whose eIF4E binding falls the most in response to eIF4E phosphorylation have very short, GC-rich 5′ UTRs. It is mechanistically unclear why eIF4E binding by mRNAs with these 5′ UTR features are the most sensitive to eIF4E phosphorylation.

In Figure 7, the 4 mRNAs with the largest reductions in eIF4E binding upon phosphorylation (~5 fold) are highlighted in blue. Highlighted in green are two ferritin mRNAs, which exhibit more modest changes in response to eIF4E phosphorylation but reach levels of particularly high significance. Analysis of the protein levels of NRAS, a Ras family member, and FTH1 (ferritin heavy chain), whose mRNA showed a more modest fold-decrease upon eIF4E phosphorylation, was conducted after inhibiting eIF4E phosphorylation in Hela cells over a 24-h time course (Figure 7C). Interestingly, we saw small (~1.8-fold) increases in NRAS or FTH1 steady-state protein levels at 4 and 8 h after MNK inhibition; however, further validation of these and other candidate mRNAs whose translation is potentially regulated by eIF4E phosphorylation will require methods that directly measure *real-time de novo* synthesis of the corresponding proteins.

Further validation of our data will require analysis of whether alterations in binding of eIF4E to a given mRNA affects the rate of synthesis of the encoded protein. This would require approaches that allow quantification of *de novo* rates of synthesis of specific polypeptides such as ‘BONCAT’ (Landgraf et al., 2015), probably coupled with pulsed stable-isotope labelling, as has been applied to study rapid changes in protein synthesis in otherwise hard-to-study cells such as primary neurons or cardiomyocytes (Kenney et al., 2016; Liu et al., 2016). These techniques will likely be invaluable for validating our leading eFT-508-capCLIP candidate mRNAs. In addition, as such methods potentially serve as potentially the most direct validation of a mRNA’s translational activity *in vivo*, we are eager to use them to validate capCLIP more broadly, as we hypothesise that capCLIP could serve as a general method for measuring the translational activity of individual mRNA species.

### The way forward for capCLIP and future prospects

We envision that capCLIP can be developed into a general technique to acquire deep, quantitative information on a cell or tissue’s present translational output, the eIF4E cap-ome. As the methodology evolves, increasing capCLIP tag library depth will permit capCLIP to be employed as a general technique for the reliable and quantitative detection of changes in translational activity even for mRNAs expressed at very low levels. We foresee using capCLIP to identify the eIF4E cap-omes of individual tumours, especially in mouse models of tumorigenesis using patient-derived cancer cell lines which can be edited to express flag-eIF4E. Especially promising is the ability, using flag-eIF4E tumour cells, to evaluate the translational status of mRNAs in metastatic cancer cells without the need to remove surrounding tumour stroma, as the host tissue containing WT eIF4E cannot ‘contaminate’ the metastatic samples. As there is little comprehensive data on metastatic proteomes/translatomes, and as such an environment is not amenable to either ribosome profiling or direct proteomic approaches, capCLIP offers a novel way to significantly expand our understanding of translational regulation in cancer.

There are further applications for capCLIP where existing technologies are not suitable, e.g., to elucidate the specific biological role(s) of other cellular cap-binding proteins, particularly eIF4E2 (4EHP) and eIF4E3, the other two cytoplasmic cap-binding proteins in mammals. While eIF4E2 is generally thought to play a role in translational repression (see, for example (Jafarnejad et al., 2018; Peter et al., 2017)), other work suggests eIF4E2 promotes mRNA translation in certain contexts (Uniacke et al., 2012). As capCLIP makes no assumptions about the cellular role of a cap-binding protein, it is ideally suited to elucidating eIF4E2 function. Likewise, capCLIP of LARP1, the nuclear cap-binding protein NCBP2, or of proteins with proposed cap-binding properties has the potential to enlarge our understanding of how dynamic networks of cap-protein interactions modulate RNA processing events throughout the cell.

## Acknowledgements

We gratefully acknowledge funding from the National Health & Medical Research Council [GNT1089167 and Research Fellowship GNT1118170 to GJG] and SAHMRI (to CGP).

## Author Contributions

Conceptualization, K.B.J. & C,G.P.; Methodology, K.B.J., B.K.D. & J.T.; Software, J.T.; Formal Analysis, J.T. & K.B.J.; Investigation, K.B.J., B.K.D., X.J., & V.I.; Writing – Original Draft, K.B.J. & C,G.P; Writing – Review & Editing, K.B.J., C,G.P, B.K.D., J.T., & G.J.G.; Supervision, K.B.J, C,G.P. & G.J.G.; Funding Acquisition, C,G.P. & G.J.G.

## Declaration of Interests

The authors declare no competing interests.

## STAR Methods

### CRISPR/Cas9-based gene editing

To introduce a 3X flag epitope tag at the N-terminus of the endogenous eIF4E gene in HeLa cells, an sgRNA guide sequence targeting the first exon of human eIF4E was designed, with the Cas9 endonuclease cut site lying 3 bases upstream of the AUG of the eIF4E open-reading frame (scheme is shown in Figure 1B). A 200 nucleotide asymmetric, single-stranded homology-directed repair (HDR) DNA template (Richardson et al., 2016) containing two modified phosphorthioate linkages at each end (Renaud et al., 2016) was synthesised to permit the introduction of a 3X-flag/1X-myc epitope tag at the extreme N-terminus of the eIF4E protein. The HDR template, along with a GeneArt CRISPR Nuclease vector (Life Technologies) expressing the eIF4E guide RNA, and a Cas9 nuclease-human CD4 pre-protein, was introduced into Hela cells using nucleofection (Lonza). 48 h post-nucleofection, Cas9-expressing cells were isolated using anti-CD4 Dynabeads, and the purified cells were plated at 1 cell/well in 4 96-well TC dishes. Following approximately 7 days of growth, wells containing clonal cell colonies were replica-plated, to allow further growth of each colony and provide cells for genomic DNA analysis. To check for insertion of the epitope tag, PCR products spanning the editing region of eIF4E were screened for length and for digestion with BspHI, a restriction site provided by the HDR template. Colonies positive in these screens were cultured further, and genomic DNA was again isolated for additional PCR amplification of the 5′-region on the eIF4E coding sequence, followed by forward and reverse Sanger sequencing of the entire edited region of eIF4E. Clones passing sequencing verification were further analysed by western blot, using both anti-flag and anti-eIF4E antibodies (Figure 1C).

### capCLIP

A brief account of the pilot capCLIP experiments and ‘preparative’ capCLIP experiments is provided here. A full account of capCLIP reagents and methodology, along with detailed commentary of aspects of the method have been included as a **Supplemental Method**. capCLIP was initially piloted on untreated Hela cells endogenously expressing flag-eIF4E. Cells were exposed to 350 mJ/cm^2^ of short-wavelength (~254 nm) UV irradiation, followed by cell lysis in 1X PXL [1X PBS supplemented with 0.1% SDS, 0.5% deoxycholate and 0.5% NP-40]. Both a ‘high’ [0.10 U/μl lysate] and ‘low’ [0.02 U/μl lysate] concentration of Ambion RNase I was tested to determine the optimal RNase I amount for generation of RNA tags of length ~30-80 nt. Anti-flag mAb bound to protein G Dynabeads was used to IP flag-eIF4E and crosslinked mRNAs, followed by three washes in 1X PXL and one wash using 5X PXL (1X PXL with 5X PBS). [γ-^32^P]ATP was used to 5′ end-label the RNA molecules, followed by a final wash, elution and Bis-Tris PAGE size separation of the labelled eIF4E:RNA complexes (Fig. 2A). Based on the initial RNase concentration experiments, an RNase I concentration of 0.02 U/μl of cell lysate was used for all subsequent capCLIP experiments.

For the preparative capCLIP experiments ± rapamycin, Hela cells endogenously expressing flag-eIF4E were treated for 2 h with 100 nM rapamycin, or with the equivalent amount of DMSO vehicle, and then UV irradiated as above. For the eFT-508 capCLIP experiment, flag-eIF4E Hela cells that had been serum-starved overnight were pre-treated with either DMSO only or with 0.1 μM eFT-508 for 1 h. Cells were subsequently stimulated with 500 nM PMA for 30 min immediately prior to UV irradiation. Cell pellets for all samples were lysed in 1X PXL supplemented with 0.02 U of RNase I per μl of cell lysate and DNAse I. Cell lysates were subject to anti-flag IP and washed, as above. A 5′-32P-labelled RNA adapter was ligated to the 3′ end of the eIF4E-crosslinked RNA ‘on-bead,’ followed by a final wash, elution and Bis-Tris PAGE size separation of the 32P-labelled eIF4E-RNA complexes. The material resolved by Bis-Tris PAGE was electrophoretically transferred to a nitrocellulose membrane, and the radioactive RNA detected using phosphorimaging (Fig. 2C). The ~37-60 kDa region of the membrane, corresponding to eIF4E:RNA complexes with RNAs approximately 30-80 nt in length, was excised, protease treated, and the liberated RNA precipitated. cDNA synthesis, followed by UMI (unique molecular identifier)-containing DNA adapter ligation to the 3′ end of the cDNA were conducted, and the resulting material was purified and size-selected on denaturing PAGE (Fig. 2D). Gel slices corresponding to ssDNAs of approximately 80-130 nt were cut from the gel, and the cDNA was eluted and precipitated. This DNA was subsequently amplified by PCR, and individual capCLIP samples were barcoded prior to sequencing. Following a final size-purification of the amplified PCR product, and QC checks of library concentration and length, all sample libraries were subject to high-throughput single-end sequencing on the Nextseq 500 with a read length of 80.

### Bioinformatic analysis

Raw reads were adaptor trimmed and filtered for short sequences during base-calling with bcl2fastq2 (--find-adapters-with-sliding-window --adapter-stringency 0.9 --mask-short-adapter-reads 35 --minimum-trimmed-read-length 35). The resulting FASTQ files, averaging 45 million reads per sample, were analysed and quality checked using FastQC (www.bioinformatics.babraham.ac.uk/projects/fastqc) and Picard (<u>broadinstitute.github.io/picard</u>). The filtered reads were mapped against the human reference genome (hg19) using the STAR alignment algorithm (Dobin et al., 2013) (version 2.5.3a with default parameters and --chimSegmentMin 20, --quantMode GeneCounts) returning an average unique alignment rate of 30%. UMItools (Smith et al., 2017) was used to deduplicate reads in each sample using unique molecular identifiers (UMI’s) (default settings with --method adjacency --edit-distance-threshold 1). Enriched regions of the genome were identified separately for each strand with the MACS2 peak-caller (Wang et al., 2008) (version 2.1.1.20160309) using default parameters and reporting only peaks with an FDR cut-off (-q) < 0.05. The resulting peak files from each strand were merged. Differential binding analysis was performed using R (version 3.4.3) and the DiffBind package (Ross-Innes et al., 2012). Alignments were visualised and interrogated using the Integrative Genomics Viewer v2.3.80 (Robinson et al., 2011).

### CAGE data

The CAGE datasets used for analysis of mRNA TSSs are ENCSR000CJJ (Hela S3 cells) and ENCSR000CKD (CD20+ B cells) from the ENCODE portal (www.encodeproject.org) (Davis et al., 2018).

### capCLIP sequence datasets

Raw sequencing files for the rapamycin capCLIP and eFT-508 capCLIP samples, along with processed peak, .bed, and .tsv files for each sample, have been uploaded to the Gene Expression Omnibus (GEO) repository at the NCBI under the accession number GSE138473. Public access will be provided upon final publication of the manuscript.

### Antibodies

For western analysis of Hela cell lysates and IPs, the following antibodies were used: anti-β-actin, mouse mAb, Sigma Aldrich A2228; anti-flag, mouse mAb (clone M2) Sigma Aldrich F3165; anti-eIF4E, rabbit polyclonal, Cell Signaling Technology 9742; anti-P-eIF4E, rabbit polyclonal, ThermoFisher 44-528G; anti-eIF4G, rabbit polyclonal, Cell Signaling Technology 2498; anti-4E-BP1, rabbit polyclonal, Cell Signaling Technology 9452; anti-rpS6, goat polyclonal, Santa Cruz Biotechnology sc-13007; anti-P-rpS6 (Ser240/244), rabbit polyclonal, Cell Signaling Technology 2215; anti-P-ERK (Thr202/Try204), rabbit polyclonal, Cell Signaling Technology 4370; anti-FTH1, rabbit polyclonal, Cell Signaling Technology 3998; anti-ras, rabbit polyclonal, Cell Signaling Technology 3965.

### Analysis of 5′ UTR length and nucleotide composition for eFT-508 capCLIP data

In total 90 mRNAs were selected: the 45 mRNAs with the greatest *reduction* in eIF4E binding upon eIF4E phosphorylation, and correspondingly, the 45 mRNAs with the greatest relative *increase* in eIF4E binding upon eIF4E phosphorylation. All selected mRNAs were subject to a minimum mRNA abundance cut-off of 2.5 to omit low-confidence mRNAs (low tag coverage) from the analysis. Ensembl Biomart and the NCBI RefSeq collection were used to source the 5′ UTR sequence of all 90 mRNAs selected. For mRNAs where the database queries returned multiple mRNA isoforms (often due to the presence of multiple TSSs and/or alternative splicing in the 5′ UTR regions of these mRNAs), eFT-508 capCLIP tag data and Hela CAGE data were viewed in IGV to determine which sequence best fit the observed capCLIP and CAGE data. When capCLIP data did not help to identify the precise pattern of 5′ UTR alternative exon usage in Hela cells, all potential 5′ UTR isoforms were included and the average 5′ UTR length used for subsequent calculations.

## Notes

### Competing Interest Statement

The authors have declared no competing interest.

### Summary of Updates

The paper has been edited for clarity and brevity.

