## Supplemental Information for "capCLIP: a new tool to probe protein synthesis in human cells through capture and identification of the eIF4E-mRNA interactome"

**Supplemental Table 1.** List of the 86 mRNAs with statistically significant changes to capCLIP peak volumes at the mRNA 5′ end ± rapamycin (p >0.01; FDR>0.05). DiffBind analysis of the alteration in capCLIP peak volumes in Hela cells treated with control (DMSO carrier only) or with 100 nM rapamycin for 2 h. The table is divided into three categories: TOP (or TOP-like) mRNAs, potential TOP mRNAs (most often these mRNAs show evidence of multiple TSSs; one with TOP-like characteristics and one without), and non-TOP mRNAs (these mRNAs show no evidence of a TOP-like motif at the TSS derived from the capCLIP and Hela CAGE data).

**Supplemental Table 2.** List of the 256 mRNAs with statistically significant changes to capCLIP peak volumes at the mRNA 5′ end ± eFT-508 pre-treatment and subsequent PMA stimulation (p >0.01; FDR>0.05). mRNAs identified through DiffBind analysis of the alteration in capCLIP peak volumes in Hela cells serum-starved overnight, pre-treated for 60 min with 30 nM eFT-508 (or a DMSO carrier alone), and then stimulated with x µM PMA for 30 min.

**Supplemental Method.** This supplement, capCLIP protocol v1.0, contains a detailed capCLIP protocol and list of reagents and core equipment necessary to carry out the method.

### Supplementary Table 1

| chr | start | end | width | Conc RAPA | Conc CTRL | log2 fold change rapa/ctrl | p value | FDR | Deatiled Annotation | gene | 1st 20 nt of mRNA | notes |
|---|---|---|---|---|---|---|---|---|---|---|---|---|
| **TOP mRNAs** | | | | | | | | | | | | |
| chr6 | 133135708 | 133135769 | 62 | 7.14 | 9.17 | -2.02 | 2.19E-40 | 1.95E-37 | 5' UTR (NM_001016, exon 1 of 6) | RPS12 | cucuuuccagccagcgccga | |
| chr20 | 60962380 | 60962455 | 76 | 3.87 | 5.69 | -1.82 | 5.77E-06 | 1.72E-04 | 5' UTR (NM_001024, exon 2 of 6) | RPS21 | cuuuucccacccccuagcgc | |
| chrX | 153626691 | 153626782 | 92 | 6.03 | 7.73 | -1.70 | 1.66E-14 | 2.64E-12 | 5' UTR (NM_006013, exon 1 of 7) | RPL10 | cuuuccggcggugacgaccu | |
| chr9 | 136215074 | 136215134 | 61 | 5.79 | 7.39 | -1.60 | 1.40E-11 | 1.38E-09 | 5' UTR (NM_000972, exon 1 of 8) | RPL7A | cuuucuguguuccuggcccg | |
| chr22 | 39715573 | 39715628 | 56 | 5.07 | 6.65 | -1.58 | 1.54E-07 | 6.53E-06 | 5' UTR (NM_001033853, exon 1 of 10) | RPL3 | cuuuccaacuuggacgcugc | |
| chr19 | 49990866 | 49990917 | 52 | 4.42 | 5.99 | -1.58 | 1.47E-05 | 3.52E-04 | 5' UTR (NM_001270491, exon 1 of 7) | RPL13A | cuuucugggcucggacctag | |
| chr5 | 81574098 | 81574218 | 121 | 7.07 | 8.55 | -1.48 | 4.78E-19 | 1.42E-16 | 5' UTR (NM_001025, exon 1 of 4) | RPS23 | cuuuuccgugcuaccugcag | |
| chr4 | 109541724 | 109541800 | 77 | 6.35 | 7.83 | -1.48 | 1.59E-11 | 1.41E-09 | 5' UTR (NM_033625, exon 1 of 6) | RPL34 | cucuuccuucucgccuaac | |
| chr1 | 45241214 | 45241304 | 91 | 6.63 | 8.09 | -1.46 | 1.79E-14 | 2.64E-12 | 5' UTR (NM_001012, exon 1 of 6) | RPS8 | cucugcggcgcgguccucgg | |
| chr8 | 56986963 | 56987066 | 104 | 5.68 | 7.12 | -1.44 | 1.59E-08 | 9.43E-07 | 5' UTR (NM_001023, exon 1 of 4) | RPS20 | cucuuccgguucuaggcgcu | |
| chr8 | 99057552 | 99057626 | 75 | 4.80 | 6.19 | -1.39 | 2.35E-05 | 5.34E-04 | 5' UTR (NM_000989, exon 2 of 5) | RPL30 | multiple TSSs | |
| chr11 | 93474575 | 93474659 | 85 | 4.22 | 5.62 | -1.39 | 2.66E-04 | 4.46E-03 | 5' UTR (NM_024116, exon 1 of 6) | TAF1D | cucuuaccgccatcttggcu | NEW TOP mRNA |
| chr19 | 39926479 | 39926591 | 113 | 6.40 | 7.74 | -1.34 | 1.41E-10 | 1.14E-08 | 5' UTR (NM_001020, exon 1 of 5) | RPS16 | cuucccucuuccggggacguu | |
| chr17 | 27047014 | 27047070 | 57 | 4.73 | 6.05 | -1.32 | 9.69E-05 | 1.83E-03 | 5' UTR (NM_000984, exon 1 of 5) | RPL23A | cuuucuggucucggccgcag | |
| chr2 | 217363546 | 217363640 | 95 | 7.62 | 8.88 | -1.26 | 6.42E-18 | 1.43E-15 | 5' UTR (NM_000998, exon 1 of 4) | RPL37A | cucuuucgcucaggccgugg | |
| chr18 | 47018781 | 47018852 | 72 | 5.82 | 7.07 | -1.25 | 1.26E-06 | 4.67E-05 | 5' UTR (NM_001199356, exon 1 of 7) | RPL17 | cucuuuccggugugg agucu | |
| chr6 | 35436162 | 35436248 | 87 | 5.74 | 6.98 | -1.25 | 1.57E-06 | 5.56E-05 | 5' UTR (NM_007104, exon 1 of 6) | RPL10A | cuuucccuggugugauuccg | |
| chr6 | 74230688 | 74230755 | 68 | 5.17 | 6.41 | -1.25 | 9.50E-05 | 1.83E-03 | 5' UTR (NM_001402, exon 1 of 8) | EEF1A1 | cucucucucuuucacugcaa | |
| chr10 | 79793619 | 79793699 | 81 | 6.61 | 7.83 | -1.22 | 1.03E-09 | 6.51E-08 | 5' UTR (NM_001142282, exon 1 of 6) | RPS24 | cucuuccacaggaggccuac | |
| chr22 | 38245419 | 38245527 | 109 | 4.83 | 6.05 | -1.22 | 2.60E-04 | 4.43E-03 | 5' UTR (NM_016091, exon 1 of 13) | EIF3L | multiple TSSs | |
| chr2 | 101618730 | 101618815 | 86 | 5.62 | 6.81 | -1.19 | 1.63E-05 | 3.80E-04 | 5' UTR (NM_000993, exon 1 of 5) | RPL31 | cucuuuuccgguuagcgcgg | |
| chr6 | 34393753 | 34393876 | 124 | 6.36 | 7.54 | -1.18 | 5.50E-08 | 2.87E-06 | 5' UTR (NM_001202470, exon 1 of 9) | RPS10 | cuuuuucgcaacggguuugc | |
| chr1 | 153963238 | 153963301 | 64 | 5.78 | 6.96 | -1.18 | 5.80E-06 | 1.72E-04 | 5' UTR (NM_001030, exon 1 of 4) | RPS27 | cucuuucccugccgccgccg | |
| chr19 | 55897300 | 55897374 | 75 | 4.49 | 5.64 | -1.15 | 1.91E-03 | 2.12E-02 | 5' UTR (NM_001136137, exon 1 of 4) | RPL28 | cuuucccuuucggauucccg | |
| chr22 | 38071643 | 38071758 | 116 | 6.41 | 7.55 | -1.14 | 1.07E-07 | 5.01E-06 | 5' UTR (NM_002305, exon 1 of 4) | LGALS1 | cuuuuugaggaagacgcggu | NEW TOP mRNA |
| chr19 | 42364284 | 42364401 | 118 | 5.93 | 7.07 | -1.14 | 7.21E-06 | 2.00E-04 | 5' UTR (NM_001022, exon 1 of 6) | RPS19 | multiple TSSs | |
| chr19 | 49122368 | 49122440 | 73 | 5.15 | 6.23 | -1.08 | 7.44E-04 | 1.05E-02 | 5' UTR (NM_000979, exon 1 of 7) | RPL18 | cucucccccggcgguuagug | |
| chr8 | 146017707 | 146017803 | 97 | 6.86 | 7.93 | -1.07 | 2.06E-07 | 8.30E-06 | 5' UTR (NM_000973, exon 1 of 6) | RPL8 | cucuuucggccgcgcuggug | |
| chr12 | 56510397 | 56510626 | 230 | 6.36 | 7.43 | -1.07 | 3.47E-06 | 1.18E-04 | 5' UTR (NM_001035267, exon 1 of 3) | RPL41 | multiple TSSs | |
| chr19 | 18682641 | 18682750 | 110 | 6.08 | 7.15 | -1.07 | 1.30E-05 | 3.30E-04 | 5' UTR (NM_003333, exon 1 of 5) | UBA52 | cucucuccucccgccgccca | |
| chr14 | 50052969 | 50053125 | 157 | 5.76 | 6.83 | -1.07 | 5.36E-05 | 1.08E-03 | 5' UTR (NM_001032, exon 1 of 3) | RPS29 | cucuuuuccuccuuggcugu | |
| chr8 | 99057690 | 99057787 | 98 | 6.04 | 7.07 | -1.03 | 3.03E-05 | 6.72E-04 | 5' UTR (NM_000989, exon 1 of 5) | RPL30 | cuuuuucucccugucgccac | |
| chr3 | 197677056 | 197677138 | 83 | 6.02 | 7.04 | -1.03 | 5.03E-05 | 1.04E-03 | 5' UTR (NM_000996, exon 1 of 5) | RPL35A | cuuuuucgucugggcugcca | |
| chr5 | 40835241 | 40835342 | 102 | 8.07 | 9.09 | -1.02 | 3.74E-13 | 4.74E-11 | 5' UTR (NM_000997, exon 1 of 4) | RPL37 | multiple TSSs | |
| chr16 | 18801576 | 18801659 | 84 | 6.95 | 7.97 | -1.02 | 1.51E-07 | 6.53E-06 | 5' UTR (NM_001030009, exon 1 of 5) | RPS15A | cuuuuccggucggcgugguc | |
| chr5 | 149829214 | 149829317 | 104 | 7.54 | 8.55 | -1.01 | 4.23E-10 | 2.89E-08 | 5' UTR (NM_005617, exon 1 of 5) | RPS14 | cucuuucucgacuccaucuu | |
| chr17 | 72199814 | 72199921 | 108 | 6.37 | 7.38 | -1.01 | 6.29E-06 | 1.80E-04 | 5' UTR (NM_001035258, exon 1 of 5) | RPL38 | cuuuccuuucagcggagcgc | |
| chr11 | 64889530 | 64889660 | 131 | 6.18 | 7.18 | -1.00 | 3.26E-05 | 7.07E-04 | 5' UTR (NM_001997, exon 1 of 5) | FAU | cuuuucuucccgcacgguug | |
| chr13 | 45915163 | 45915317 | 155 | 4.85 | 5.85 | -1.00 | 3.88E-03 | 3.87E-02 | 5' UTR (NM_003295, exon 1 of 6) | TPT1 | multiple TSSs | |
| chr1 | 235291958 | 235292151 | 194 | 5.31 | 6.31 | -0.99 | 1.20E-03 | 1.56E-02 | 5' UTR (NM_014765, exon 1 of 5) | TOMM20 | cuuuuccgcccgcucccccc | |
| chr11 | 32605362 | 32605500 | 139 | 5.24 | 6.23 | -0.99 | 1.74E-03 | 1.96E-02 | 5' UTR (NM_001307929, exon 1 of 8) | EIF3M | multiple TSSs | |
| chr19 | 49999683 | 49999751 | 69 | 5.68 | 6.67 | -0.98 | 4.28E-04 | 6.67E-03 | 5' UTR (NM_001015, exon 1 of 5) | RPS11 | ccuuuccucagcugccgcca | |
| chr17 | 37009914 | 37009987 | 74 | 7.29 | 8.24 | -0.95 | 6.49E-08 | 3.20E-06 | 5' UTR (NM_000978, exon 1 of 5) | RPL23 | cucuuucgccaucuuuccg | |
| chr8 | 101734094 | 101734316 | 223 | 5.60 | 6.55 | -0.95 | 9.07E-04 | 1.26E-02 | 5' UTR (NM_002568, exon 1 of 15) | PABPC1 | cucuucccuuuugcggccau | |
| chr11 | 75110560 | 75110636 | 77 | 6.64 | 7.57 | -0.94 | 9.29E-06 | 2.50E-04 | 5' UTR (NM_001256802, exon 1 of 7) | RPS3 | multiple TSSs | |
| chr20 | 60962121 | 60962233 | 113 | 7.15 | 8.06 | -0.92 | 3.60E-07 | 1.39E-05 | 5' UTR (NM_001024, exon 1 of 6) | RPS21 | cuuccuuuuuucuuuuuucc | |
| chr6 | 33239852 | 33239936 | 85 | 6.55 | 7.46 | -0.91 | 3.64E-05 | 7.70E-04 | 5' UTR (NM_022551, exon 1 of 6) | RPS18 | multiple TSSs | |
| chr19 | 58898636 | 58898731 | 96 | 6.74 | 7.64 | -0.90 | 1.03E-05 | 2.69E-04 | 5' UTR (NM_001009, exon 1 of 6) | RPS5 | cucuuucccuaagcagccug | |
| chr3 | 40498817 | 40498888 | 72 | 6.06 | 6.96 | -0.90 | 5.09E-04 | 7.79E-03 | 5' UTR (NM_001034996, exon 1 of 6) | RPL14 | cucuuccgccgucgucgccg | |
| chr1 | 93297594 | 93297697 | 104 | 6.07 | 6.95 | -0.89 | 5.74E-04 | 8.35E-03 | 5' UTR (NM_000969, exon 1 of 8) | RPL5 | cuuuuucuucagcgaggcgg | |
| chr13 | 49066713 | 49066834 | 122 | 5.30 | 6.18 | -0.88 | 4.66E-03 | 4.40E-02 | 5' UTR (NM_000884, exon 1 of 14) | IMPDH2 | cuuuuccgguugcggcgccg | |
| chr3 | 101405487 | 101405561 | 75 | 6.24 | 7.07 | -0.83 | 5.73E-04 | 8.35E-03 | 5' UTR (NM_000986, exon 1 of 6) | RPL24 | cuuccccuggctggcagca | |
| chr19 | 3985354 | 3985461 | 108 | 5.96 | 6.79 | -0.83 | 1.71E-03 | 1.95E-02 | 5' UTR (NM_001961, exon 1 of 15) | EEF2 | cuuuccggaccuggccgagc | |
| chr11 | 17099137 | 17099253 | 117 | 5.85 | 6.68 | -0.83 | 2.11E-03 | 2.29E-02 | 5' UTR (NM_001017, exon 1 of 6) | RPS13 | cuuuuccaagcggcugccga | |
| chr11 | 809954 | 810062 | 109 | 5.90 | 6.73 | -0.82 | 2.08E-03 | 2.28E-02 | 5' UTR (NM_001004, exon 1 of 5) | RPLP2 | multiple TSSs | |
| chr3 | 52029840 | 52029956 | 117 | 6.11 | 6.92 | -0.81 | 1.28E-03 | 1.61E-02 | 5' UTR (NM_000992, exon 1 of 4) | RPL29 | cucuuuccgucucaggucgc | |
| chr5 | 170814854 | 170814940 | 87 | 7.12 | 7.93 | -0.80 | 1.40E-05 | 3.45E-04 | 5' UTR (NM_001037738, exon 1 of 10) | NPM1 | ctcttcctgtcuguaccagg | |
| chr11 | 8704317 | 8704404 | 88 | 5.93 | 6.74 | -0.80 | 2.38E-03 | 2.51E-02 | 5' UTR (NM_000990, exon 1 of 5) | RPL27A | cuuucucucucgcgcgcggu | |
| chr3 | 39448212 | 39448267 | 56 | 6.86 | 7.65 | -0.79 | 9.66E-05 | 1.83E-03 | 5' UTR (NM_002295, exon 1 of 7) | RPSA | cucucucgggugg agucuuc | |
| chr17 | 8286443 | 8286528 | 86 | 6.55 | 7.31 | -0.76 | 6.76E-04 | 9.69E-03 | 5' UTR (NM_000987, exon 1 of 4) | RPL26 | cuuuccggcggugcucgcaa | |
| chr9 | 127624150 | 127624251 | 102 | 6.82 | 7.50 | -0.68 | 1.16E-03 | 1.53E-02 | 5' UTR (NM_007209, exon 1 of 4) | RPL35 | cucuaccggcgggauuugau | |
| chr15 | 69745157 | 69745307 | 151 | 7.08 | 7.71 | -0.63 | 1.13E-03 | 1.52E-02 | 5' UTR (NM_001003, exon 1 of 4) | RPLP1 | cucuuucccuucggugugcc | |
| **potential TOP mRNAs** | | | | | | | | | | | | |
| chr10 | 71993033 | 71993179 | 147 | 3.82 | 5.18 | -1.35 | 1.71E-03 | 1.95E-02 | 5' UTR (NM_021129, exon 1 of 11) | PPA1 | | |
| chr6 | 160210429 | 160210694 | 266 | 4.44 | 5.76 | -1.33 | 2.81E-04 | 4.61E-03 | 5' UTR (NM_001008897, exon 1 of 11) | TCP1 | | |
| chr12 | 56498356 | 56498521 | 166 | 4.08 | 5.28 | -1.19 | 3.98E-03 | 3.93E-02 | 5' UTR (NM_006191, exon 1 of 13) | PA2G4 | | |
| chr7 | 26241352 | 26241478 | 127 | 5.37 | 6.55 | -1.18 | 1.45E-04 | 2.63E-03 | 5' UTR (NM_016587, exon 1 of 6) | CBX3 | | |
| chr14 | 69864921 | 69865051 | 131 | 4.61 | 5.76 | -1.15 | 1.67E-03 | 1.95E-02 | 5' UTR (NM_004450, exon 1 of 4) | ERH | | |
| chr5 | 68665265 | 68665367 | 103 | 4.59 | 5.64 | -1.06 | 4.58E-03 | 4.38E-02 | 5' UTR (NM_003187, exon 1 of 3) | TAF9 | | |
| chr12 | 57119006 | 57119122 | 117 | 5.39 | 6.43 | -1.03 | 5.40E-04 | 8.13E-03 | 5' UTR (NM_001113203, exon 1 of 11) | NACA | | |
| chr1 | 46769337 | 46769531 | 195 | 5.25 | 6.28 | -1.02 | 1.47E-03 | 1.75E-02 | 5' UTR (NM_006004, exon 1 of 4) | UQCRH | | |
| chr6 | 27775950 | 27776034 | 85 | 5.15 | 6.16 | -1.01 | 1.48E-03 | 1.75E-02 | 5' UTR (NM_003509, exon 1 of 1) | HIST1H2AI | | |
| chr6 | 41755406 | 41755554 | 149 | 5.16 | 6.11 | -0.94 | 3.06E-03 | 3.20E-02 | 5' UTR (NM_001134493, exon 1 of 3) | TOMM6 | | |
| chr17 | 49230920 | 49231047 | 128 | 5.45 | 6.37 | -0.92 | 3.27E-03 | 3.38E-02 | 5' UTR (NM_000269, exon 1 of 5) | NME1 | | |
| chr2 | 198364490 | 198364604 | 115 | 6.11 | 6.91 | -0.81 | 1.29E-03 | 1.61E-02 | 5' UTR (NM_002156, exon 1 of 12) | HSPD1 | | |
| chr6 | 30688122 | 30688328 | 207 | 6.94 | 7.59 | -0.65 | 1.36E-03 | 1.66E-02 | 5' UTR (NM_178014, exon 1 of 4).7 | TUBB | | |
| **non-TOP mRNAs** | | | | | | | | | | | | |
| chr1 | 40506411 | 40506506 | 96 | 2.86 | 4.78 | -1.92 | 3.40E-04 | 5.40E-03 | 5' UTR (NM_006367, exon 1 of 13) | CAP1 | | |
| chr15 | 79165190 | 79165402 | 213 | 2.57 | 4.48 | -1.91 | 2.28E-03 | 2.44E-02 | 5' UTR (NM_006791, exon 1 of 12) | MORF4L1 | | |
| chr6 | 111279904 | 111280027 | 124 | 4.69 | 5.95 | -1.27 | 2.52E-04 | 4.39E-03 | 5' UTR (NM_138408, exon 1 of 6) | GTF3C6 | | |
| chr12 | 49525055 | 49525180 | 126 | 7.62 | 8.73 | -1.11 | 4.69E-13 | 5.21E-11 | 5' UTR (NM_006082, exon 1 of 4) | TUBA1B | | |
| chr7 | 25164807 | 25164884 | 78 | 6.28 | 7.33 | -1.05 | 3.99E-06 | 1.31E-04 | 5' UTR (NM_018947, exon 1 of 3) | CYCS | | |
| chr7 | 5632444 | 5632564 | 121 | 5.14 | 6.16 | -1.02 | 1.22E-03 | 1.58E-02 | 5' UTR (NM_003088, exon 1 of 5) | FSCN1 | | |
| chr17 | 7210877 | 7211071 | 195 | 5.75 | 6.76 | -1.01 | 1.65E-04 | 2.93E-03 | 5' UTR (NM_001970, exon 1 of 6) | EIF5A | | |
| chr11 | 61734986 | 61735104 | 119 | 11.52 | 12.47 | -0.95 | 3.93E-34 | 1.74E-31 | 5' UTR (NM_002032, exon 1 of 4) | FTH1 | | |
| chr12 | 56552141 | 56552225 | 85 | 6.18 | 7.07 | -0.89 | 2.97E-04 | 4.79E-03 | 5' UTR (NM_021019, exon 1 of 7) | MYL6 | | |
| chr9 | 113018694 | 113018859 | 166 | 7.27 | 7.97 | -0.70 | 1.36E-04 | 2.52E-03 | 5' UTR (NM_003329, exon 1 of 5) | TXN | | |
| chr7 | 26240194 | 26240393 | 200 | 7.13 | 7.75 | -0.62 | 1.36E-03 | 1.66E-02 | 5' UTR (NM_031243, exon 1 of 12) | HNRNPA2B1 | | |

FDR is 0.05 or less
p value is 0.01 or less
fold change is -0.50 or less, or 0.50 or more

### Supplementary Table 2

| chr | start | end | width | Conc eFT508 | Conc PMA | log2 fold change eFT508/DMSO | p value | FDR | Deatiled Annotation | Gene |
|---|---|---|---|---|---|---|---|---|---|---|
| **mRNAs altered with eFT-508 treatment (FDR < 0.05)** | | | | | | | | | | |
| chr11 | 61734986 | 61735104 | 119 | 13.41 | 12.21 | 1.20 | 7.71E-54 | 9.35E-51 | 5UTR (NM_002032, exon 1 of 4, FTH1) | FTH1 |
| chr6 | 133135708 | 133135769 | 62 | 8.63 | 7.14 | 1.48 | 1.87E-17 | 1.13E-14 | 5UTR (NM_001016, exon 1 of 6, RPS12) | RPS12 |
| chr19 | 49468564 | 49468803 | 240 | 8.40 | 6.78 | 1.62 | 5.44E-17 | 2.20E-14 | 5UTR (NM_000146, exon 1 of 4, FTL) | FTL |
| chr11 | 18416108 | 18416219 | 112 | 8.39 | 6.89 | 1.49 | 2.79E-16 | 8.46E-14 | 5UTR (NM_001165416, exon 1 of 7, LDHA) | LDHA |
| chr7 | 44836277 | 44836403 | 127 | 8.88 | 7.67 | 1.21 | 5.55E-14 | 1.13E-11 | 5UTR (NM_001300981, exon 1 of 6, PPIA) | PPIA |
| chr8 | 27472127 | 27472220 | 94 | 9.26 | 8.18 | 1.08 | 5.59E-14 | 1.13E-11 | 5UTR (NM_001831, exon 1 of 9, CLU) | CLU |
| chr9 | 19380158 | 19380238 | 81 | 8.70 | 7.53 | 1.18 | 2.86E-12 | 4.33E-10 | 5UTR (NM_001010, exon 1 of 6, RPS6) | RPS6 |
| chr5 | 40835241 | 40835340 | 100 | 9.49 | 8.46 | 1.03 | 2.76E-12 | 4.33E-10 | 5UTR (NM_000997, exon 1 of 4, RPL37) | RPL37 |
| chr20 | 57607286 | 57607410 | 125 | 8.53 | 7.42 | 1.11 | 1.34E-10 | 1.81E-08 | 5UTR (NM_006886, exon 1 of 3, ATP5E) | ATP5E |
| chr8 | 56986974 | 56987067 | 94 | 7.33 | 5.86 | 1.47 | 1.00E-09 | 1.10E-07 | 5UTR (NM_001023, exon 1 of 4, RPS20) | RPS20 |
| chr6 | 27114531 | 27114634 | 104 | 7.90 | 6.61 | 1.29 | 9.77E-10 | 1.10E-07 | 5UTR (NM_080593, exon 1 of 2, HIST1H2BK) | HIST1H2BK |
| chr15 | 66797092 | 66797183 | 92 | 6.72 | 4.87 | 1.85 | 4.34E-09 | 3.68E-07 | 5UTR (NM_000968, exon 1 of 10, RPL4) | RPL4 |
| chr14 | 102553303 | 102553381 | 79 | 7.14 | 5.60 | 1.55 | 4.54E-09 | 3.68E-07 | 5UTR (NM_005348, exon 1 of 11, HSP90AA1) | HSP90AA1 |
| chr1 | 153963237 | 153963308 | 72 | 7.26 | 5.83 | 1.43 | 4.55E-09 | 3.68E-07 | 5UTR (NM_001030, exon 1 of 4, RPS27) | RPS27 |
| chr20 | 60962121 | 60962233 | 113 | 8.33 | 7.30 | 1.03 | 4.28E-09 | 3.68E-07 | 5UTR (NM_001024, exon 1 of 6, RPS21) | RPS21 |
| chr2 | 3622866 | 3622984 | 119 | 7.27 | 5.85 | 1.42 | 7.43E-09 | 5.63E-07 | 5UTR (NM_001011, exon 1 of 7, RPS7) | RPS7 |
| chr9 | 127624150 | 127624251 | 102 | 7.63 | 6.44 | 1.18 | 3.22E-08 | 2.25E-06 | 5UTR (NM_007209, exon 1 of 4, RPL35) | RPL35 |
| chr6 | 26156555 | 26156669 | 115 | 8.71 | 7.79 | 0.91 | 3.34E-08 | 2.25E-06 | 5UTR (NM_005321, exon 1 of 1, HIST1H1E) | HIST1H1E |
| chr16 | 18801579 | 18801660 | 82 | 8.60 | 7.65 | 0.94 | 5.04E-08 | 3.22E-06 | 5UTR (NM_001019, exon 1 of 5, RPS15A) | RPS15A |
| chr5 | 81574098 | 81574218 | 121 | 8.38 | 7.45 | 0.93 | 5.68E-08 | 3.38E-06 | 5UTR (NM_001025, exon 1 of 4, RPS23) | RPS23 |
| chr5 | 10250357 | 10250537 | 181 | 8.45 | 7.52 | 0.93 | 5.85E-08 | 3.38E-06 | 5UTR (NM_012073, exon 1 of 11, CCT5) | CCT5 |
| chr2 | 101618737 | 101618815 | 79 | 7.46 | 6.27 | 1.19 | 1.56E-07 | 8.58E-06 | 5UTR (NM_001098577, exon 1 of 5, RPL31) | RPL31 |
| chr3 | 52029840 | 52029940 | 101 | 7.53 | 6.39 | 1.14 | 1.80E-07 | 9.48E-06 | 5UTR (NM_000992, exon 1 of 4, RPL29) | RPL29 |
| chr20 | 17550708 | 17550872 | 165 | 7.08 | 5.71 | 1.37 | 2.04E-07 | 1.03E-05 | 5UTR (NM_006870, exon 1 of 4, DSTN) | DSTN |
| chr1 | 93297595 | 93297697 | 103 | 6.99 | 5.62 | 1.37 | 2.13E-07 | 1.03E-05 | 5UTR (NM_000969, exon 1 of 8, RPL5) | RPL5 |
| chr15 | 44084516 | 44084626 | 111 | 6.86 | 5.37 | 1.49 | 2.33E-07 | 1.05E-05 | 5UTR (NM_001018108, exon 1 of 3, SERF2) | SERF2 |
| chr1 | 153508391 | 153508530 | 140 | 7.85 | 6.76 | 1.09 | 2.33E-07 | 1.05E-05 | 5UTR (NM_014624, exon 1 of 3, S100A6) | S100A6 |
| chr17 | 8286424 | 8286512 | 89 | 8.05 | 7.08 | 0.96 | 2.88E-07 | 1.25E-05 | 5UTR (NM_001315530, exon 1 of 4, RPL26) | RPL26 |
| chr15 | 69745157 | 69745318 | 162 | 8.34 | 7.38 | 0.96 | 3.46E-07 | 1.44E-05 | 5UTR (NM_001003, exon 1 of 4, RPLP1) | RPLP1 |
| chr19 | 55897300 | 55897365 | 66 | 6.13 | 4.18 | 1.94 | 4.38E-07 | 1.66E-05 | 5UTR (NM_000991, exon 1 of 5, RPL28) | RPL28 |
| chr18 | 47018785 | 47018856 | 72 | 7.39 | 6.21 | 1.18 | 4.35E-07 | 1.66E-05 | 5UTR (NM_001199342, exon 1 of 8, RPL17) | RPL17 |
| chr11 | 65625528 | 65625684 | 157 | 8.41 | 7.54 | 0.88 | 4.15E-07 | 1.66E-05 | 5UTR (NM_005507, exon 1 of 4, CFL1) | CFL1 |
| chr3 | 40498822 | 40498889 | 68 | 7.23 | 6.02 | 1.21 | 4.75E-07 | 1.69E-05 | 5UTR (NM_003973, exon 1 of 6, RPL14) | RPL14 |
| chr2 | 55459785 | 55459978 | 194 | 7.51 | 6.34 | 1.16 | 4.70E-07 | 1.69E-05 | 5UTR (NM_002954, exon 2 of 6, RPS27A) | RPS27A |
| chr16 | 30077081 | 30077232 | 152 | 6.77 | 5.22 | 1.55 | 6.22E-07 | 2.15E-05 | 5UTR (NM_001243177, exon 1 of 10, ALDOA) | ALDOA |
| chr16 | 89627073 | 89627160 | 88 | 7.25 | 6.06 | 1.19 | 6.45E-07 | 2.17E-05 | 5UTR (NM_001243131, exon 1 of 7, RPL13) | RPL13 |
| chr5 | 159848846 | 159848939 | 94 | 7.09 | 5.85 | 1.24 | 6.80E-07 | 2.23E-05 | 5UTR (NM_004219, exon 1 of 6, PTTG1) | PTTG1 |
| chr16 | 2827198 | 2827286 | 89 | 6.20 | 4.38 | 1.82 | 7.07E-07 | 2.25E-05 | 5UTR (NM_007108, exon 1 of 4, ELOB) | ELOB |
| chr8 | 146017706 | 146017807 | 102 | 7.87 | 6.91 | 0.96 | 8.32E-07 | 2.59E-05 | 5UTR (NM_033301, exon 1 of 6, RPL8) | RPL8 |
| chr19 | 5690242 | 5690382 | 141 | 6.25 | 4.53 | 1.73 | 1.01E-06 | 3.07E-05 | 5UTR (NM_033643, exon 1 of 4, RPL36) | RPL36 |
| chr3 | 12882950 | 12883044 | 95 | 7.36 | 6.26 | 1.10 | 1.12E-06 | 3.30E-05 | 5UTR (NM_001007073, exon 1 of 4, RPL32) | RPL32 |
| chr16 | 87902953 | 87903112 | 160 | 7.01 | 5.75 | 1.26 | 1.27E-06 | 3.67E-05 | 5UTR (NM_003486, exon 1 of 10, SLC7A5) | SLC7A5 |
| chr1 | 45241213 | 45241312 | 100 | 7.85 | 6.84 | 1.02 | 1.54E-06 | 4.34E-05 | 5UTR (NM_001012, exon 1 of 6, RPS8) | RPS8 |
| chr20 | 60718260 | 60718474 | 215 | 6.46 | 4.94 | 1.52 | 2.05E-06 | 5.63E-05 | 5UTR (NM_002792, exon 1 of 7, PSMA7) | PSMA7 |
| chr12 | 98987451 | 98987558 | 108 | 7.32 | 6.22 | 1.10 | 2.25E-06 | 5.92E-05 | 5UTR (NM_213611, exon 1 of 7, SLC25A3) | SLC25A3 |
| chr5 | 170814854 | 170814939 | 86 | 8.06 | 7.19 | 0.87 | 2.25E-06 | 5.92E-05 | 5UTR (NM_002520, exon 1 of 11, NPM1) | NPM1 |
| chr1 | 226250416 | 226250475 | 60 | 7.27 | 6.12 | 1.15 | 2.30E-06 | 5.93E-05 | 5UTR (NM_002107, exon 1 of 4, H3F3A) | H3F3A |
| chr5 | 130500835 | 130500993 | 159 | 6.77 | 5.46 | 1.32 | 2.49E-06 | 6.28E-05 | 5UTR (NM_005340, exon 1 of 3, HINT1) | HINT1 |
| chr19 | 39926488 | 39926590 | 103 | 7.93 | 7.03 | 0.91 | 2.58E-06 | 6.38E-05 | 5UTR (NM_001020, exon 1 of 5, RPS16) | RPS16 |
| chr1 | 155278628 | 155278798 | 171 | 7.58 | 6.56 | 1.02 | 3.02E-06 | 7.17E-05 | 5UTR (NM_001242824, exon 1 of 10, FDPS) | FDPS |
| chr1 | 153518185 | 153518282 | 98 | 7.86 | 6.90 | 0.96 | 3.00E-06 | 7.17E-05 | 5UTR (NM_002961, exon 1 of 3, S100A4) | S100A4 |
| chr19 | 18433792 | 18433926 | 135 | 5.73 | 3.58 | 2.15 | 3.72E-06 | 8.50E-05 | 5UTR (NM_001252129, exon 1 of 4, LSM4) | LSM4 |
| chr4 | 109541723 | 109541830 | 108 | 7.74 | 6.79 | 0.95 | 3.69E-06 | 8.50E-05 | 5UTR (NM_000995, exon 1 of 6, RPL34) | RPL34 |
| chr6 | 34393753 | 34393867 | 115 | 7.51 | 6.52 | 0.99 | 3.81E-06 | 8.56E-05 | 5UTR (NM_001203245, exon 1 of 6, RPS10) | RPS10 |
| chrX | 153626657 | 153626886 | 230 | 7.28 | 6.22 | 1.06 | 4.36E-06 | 9.43E-05 | 5UTR (NM_006013, exon 1 of 7, RPL10) | RPL10 |
| chr19 | 58898636 | 58898734 | 99 | 7.98 | 7.09 | 0.89 | 4.30E-06 | 9.43E-05 | 5UTR (NM_001009, exon 1 of 6, RPS5) | RPS5 |
| chr2 | 120124518 | 120124661 | 144 | 5.99 | 4.18 | 1.80 | 5.17E-06 | 1.09E-04 | 5UTR (NM_001178042, exon 1 of 5, DBI) | DBI |
| chr9 | 130213575 | 130213685 | 111 | 6.10 | 4.38 | 1.72 | 5.21E-06 | 1.09E-04 | 5UTR (NM_000976, exon 1 of 7, RPL12) | RPL12 |
| chr19 | 42364280 | 42364401 | 122 | 7.10 | 5.97 | 1.13 | 6.21E-06 | 1.28E-04 | 5UTR (NM_001022, exon 1 of 6, RPS19) | RPS19 |
| chr11 | 61559992 | 61560085 | 94 | 6.35 | 4.89 | 1.45 | 8.63E-06 | 1.72E-04 | 5UTR (NM_014206, exon 1 of 4, TMEM258) | TMEM258 |
| chr17 | 4851547 | 4851857 | 311 | 7.35 | 6.33 | 1.01 | 8.56E-06 | 1.72E-04 | 5UTR (NM_005022, exon 1 of 3, PFN1) | PFN1 |
| chr5 | 149829227 | 149829313 | 87 | 8.92 | 8.28 | 0.65 | 9.24E-06 | 1.81E-04 | 5UTR (NM_001025071, exon 1 of 5, RPS14) | RPS14 |
| chr12 | 56510397 | 56510627 | 231 | 7.85 | 6.97 | 0.88 | 1.09E-05 | 2.10E-04 | 5UTR (NM_001035267, exon 1 of 3, RPL41) | RPL41 |
| chr6 | 33239853 | 33239937 | 85 | 7.66 | 6.78 | 0.89 | 1.28E-05 | 2.43E-04 | 5UTR (NM_022551, exon 1 of 6, RPS18) | RPS18 |
| chr19 | 3985360 | 3985461 | 102 | 7.39 | 6.38 | 1.01 | 1.90E-05 | 3.54E-04 | 5UTR (NM_001961, exon 1 of 15, EEF2) | EEF2 |
| chr22 | 38245421 | 38245481 | 61 | 6.34 | 4.94 | 1.40 | 1.98E-05 | 3.64E-04 | 5UTR (NM_001242923, exon 1 of 11, EIF3L) | EIF3L |
| chr19 | 49990866 | 49990917 | 52 | 6.17 | 4.66 | 1.51 | 2.07E-05 | 3.74E-04 | 5UTR (NM_001270491, exon 1 of 7, RPL13A) | RPL13A |
| chr7 | 25164795 | 25164901 | 107 | 7.70 | 6.84 | 0.86 | 2.43E-05 | 4.33E-04 | 5UTR (NM_018947, exon 1 of 3, CYCS) | CYCS |
| chr15 | 60690115 | 60690186 | 72 | 6.34 | 4.98 | 1.36 | 2.62E-05 | 4.45E-04 | 5UTR (NM_001002858, exon 1 of 13, ANXA2) | ANXA2 |
| chr5 | 32585638 | 32585759 | 122 | 7.03 | 5.99 | 1.04 | 2.63E-05 | 4.45E-04 | 5UTR (NM_006713, exon 1 of 5, SUB1) | SUB1 |
| chr2 | 85132777 | 85132870 | 94 | 7.64 | 6.73 | 0.90 | 2.56E-05 | 4.45E-04 | 5UTR (NM_021103, exon 1 of 3, TMSB10) | TMSB10 |
| chr3 | 39448211 | 39448275 | 65 | 7.99 | 7.20 | 0.78 | 2.64E-05 | 4.45E-04 | 5UTR (NM_001304288, exon 1 of 7, RPSA) | RPSA |
| chr1 | 8938621 | 8938770 | 150 | 7.56 | 6.69 | 0.87 | 2.94E-05 | 4.87E-04 | 5UTR (NM_001428, exon 1 of 12, ENO1) | ENO1 |
| chr9 | 86595406 | 86595542 | 137 | 5.96 | 4.29 | 1.67 | 3.08E-05 | 5.04E-04 | 5UTR (NM_031262, exon 1 of 17, HNRNPK) | HNRNPK |
| chr11 | 75110561 | 75110655 | 95 | 7.88 | 7.08 | 0.80 | 3.17E-05 | 5.13E-04 | 5UTR (NM_001260507, exon 1 of 6, RPS3) | RPS3 |
| chr19 | 18682640 | 18682749 | 110 | 7.39 | 6.48 | 0.91 | 3.45E-05 | 5.49E-04 | 5UTR (NM_001033930, exon 1 of 5, UBA52) | UBA52 |
| chr3 | 197677061 | 197677139 | 79 | 7.29 | 6.33 | 0.96 | 3.88E-05 | 6.10E-04 | 5UTR (NM_000996, exon 1 of 5, RPL35A) | RPL35A |
| chr22 | 24236568 | 24236752 | 185 | 7.15 | 6.18 | 0.97 | 4.01E-05 | 6.15E-04 | 5UTR (NM_002415, exon 1 of 3, MIF) | MIF |
| chr9 | 113018689 | 113018868 | 180 | 7.92 | 7.14 | 0.78 | 3.99E-05 | 6.15E-04 | 5UTR (NM_003329, exon 1 of 5, TXN) | TXN |
| chr1 | 26232848 | 26232946 | 99 | 6.64 | 5.49 | 1.15 | 4.73E-05 | 7.17E-04 | 5UTR (NM_005563, exon 1 of 5, STMN1) | STMN1 |
| chr6 | 87804733 | 87804830 | 98 | 7.02 | 6.02 | 1.01 | 5.16E-05 | 7.72E-04 | 5UTR (NM_001252383, exon 1 of 5, CGA) | CGA |
| chr12 | 56552132 | 56552225 | 94 | 6.99 | 5.98 | 1.01 | 5.25E-05 | 7.76E-04 | 5UTR (NM_079423, exon 1 of 6, MYL6) | MYL6 |
| chr2 | 189839085 | 189839275 | 191 | 7.45 | 6.56 | 0.88 | 6.48E-05 | 9.46E-04 | 5UTR (NM_000090, exon 1 of 51, COL3A1) | COL3A1 |
| chr12 | 49525043 | 49525180 | 138 | 7.50 | 6.67 | 0.84 | 7.54E-05 | 1.09E-03 | 5UTR (NM_006082, exon 1 of 4, TUBA1B) | TUBA1B |
| chr17 | 73042990 | 73043074 | 85 | 5.45 | 3.47 | 1.98 | 8.22E-05 | 1.17E-03 | 5UTR (NM_001003785, exon 1 of 5, ATP5H) | ATP5H |
| chr1 | 8021725 | 8021850 | 126 | 6.58 | 5.37 | 1.21 | 8.29E-05 | 1.17E-03 | 5UTR (NM_007262, exon 1 of 7, PARK7) | PARK7 |
| chr1 | 149783858 | 149783929 | 72 | 6.66 | 5.50 | 1.16 | 1.03E-04 | 1.43E-03 | 5UTR (NM_001024599, exon 1 of 1, HIST2H2BF) | HIST2H2BF |
| chr11 | 809941 | 810062 | 122 | 6.94 | 5.95 | 0.99 | 1.04E-04 | 1.43E-03 | 5UTR (NM_001004, exon 1 of 5, RPLP2) | RPLP2 |

### Supplementary Table 2 (cont.)

| chr | start | end | width | Conc eFT508 | Conc PMA | log2 fold change eFT508/DMSO | p value | FDR | Deatiled Annotation | Gene |
|---|---|---|---|---|---|---|---|---|---|---|
| chr9 | 91926110 | 91926258 | 149 | 7.66 | 6.85 | 0.81 | 1.12E-04 | 1.52E-03 | 5UTR (NM_001827, exon 1 of 3, CKS2) | CKS2 |
| chr17 | 49243758 | 49243903 | 146 | 5.72 | 4.07 | 1.65 | 1.20E-04 | 1.59E-03 | 5UTR (NM_001198682, exon 1 of 4, NME2) | NME2 |
| chr11 | 17099136 | 17099260 | 125 | 7.00 | 6.04 | 0.96 | 1.21E-04 | 1.60E-03 | 5UTR (NM_001017, exon 1 of 6, RPS13) | RPS13 |
| chr10 | 33247054 | 33247194 | 141 | 5.67 | 4.00 | 1.67 | 1.27E-04 | 1.65E-03 | 5UTR (NM_002211, exon 1 of 16, ITGB1) | ITGB1 |
| chr19 | 1438365 | 1438459 | 95 | 6.23 | 4.94 | 1.29 | 1.29E-04 | 1.67E-03 | 5UTR (NM_001018, exon 1 of 4, RPS15) | RPS15 |
| chr7 | 128379390 | 128379512 | 123 | 6.32 | 5.09 | 1.22 | 1.33E-04 | 1.70E-03 | 5UTR (NM_001219, exon 1 of 7, CALU) | CALU |
| chr7 | 26240198 | 26240393 | 196 | 7.93 | 7.23 | 0.70 | 1.36E-04 | 1.72E-03 | 5UTR (NM_031243, exon 1 of 12, HNRNPA2B1) | HNRNPA2B1 |
| chr15 | 75230287 | 75230473 | 187 | 5.06 | 2.73 | 2.33 | 1.64E-04 | 2.04E-03 | 5UTR (NM_004255, exon 1 of 5, COX5A) | COX5A |
| chr1 | 28562623 | 28562755 | 133 | 5.86 | 4.35 | 1.51 | 1.65E-04 | 2.04E-03 | 5UTR (NM_178190, exon 1 of 3, ATPIF1) | ATPIF1 |
| chr12 | 48357367 | 48357457 | 91 | 5.74 | 4.15 | 1.60 | 1.72E-04 | 2.10E-03 | 5UTR (NM_024056, exon 1 of 8, TMEM106C) | TMEM106C |
| chr10 | 17271276 | 17271508 | 233 | 6.34 | 5.17 | 1.17 | 2.16E-04 | 2.62E-03 | 5UTR (NM_003380, exon 2 of 10, VIM) | VIM |
| chr5 | 37371035 | 37371209 | 175 | 5.64 | 4.04 | 1.60 | 2.43E-04 | 2.92E-03 | 5UTR (NM_004298, exon 1 of 35, NUP155) | NUP155 |
| chr7 | 26241353 | 26241484 | 132 | 6.64 | 5.62 | 1.02 | 2.56E-04 | 3.03E-03 | 5UTR (NM_016587, exon 1 of 6, CBX3) | CBX3 |
| chr7 | 10979645 | 10979813 | 169 | 7.35 | 6.55 | 0.80 | 2.58E-04 | 3.03E-03 | 5UTR (NM_002489, exon 1 of 4, NDUFA4) | NDUFA4 |
| chr2 | 98262509 | 98262590 | 82 | 6.08 | 4.78 | 1.29 | 2.66E-04 | 3.10E-03 | 5UTR (NM_001862, exon 1 of 4, COX5B) | COX5B |
| chr1 | 115259298 | 115259391 | 94 | 4.71 | 2.15 | 2.56 | 3.11E-04 | 3.56E-03 | 5UTR (NM_002524, exon 1 of 7, NRAS) | NRAS |
| chr10 | 102106867 | 102107263 | 397 | 7.10 | 6.22 | 0.88 | 3.10E-04 | 3.56E-03 | 5UTR (NM_005063, exon 1 of 6, SCD) | SCD |
| chr17 | 72199816 | 72199919 | 104 | 7.36 | 6.56 | 0.81 | 3.22E-04 | 3.65E-03 | 5UTR (NM_001035258, exon 1 of 5, RPL38) | RPL38 |
| chr6 | 36853702 | 36853820 | 119 | 4.34 | 1.15 | 3.19 | 3.40E-04 | 3.81E-03 | 5UTR (NM_001286635, exon 1 of 9, C6orf89) | C6orf89 |
| chr1 | 235291958 | 235292152 | 195 | 6.67 | 5.70 | 0.97 | 3.70E-04 | 4.11E-03 | 5UTR (NM_014765, exon 1 of 5, TOMM20) | TOMM20 |
| chr12 | 131356586 | 131356700 | 115 | 6.68 | 5.70 | 0.98 | 3.93E-04 | 4.33E-03 | 5UTR (NM_001300797, exon 1 of 6, RAN) | RAN |
| chr9 | 95055845 | 95055998 | 154 | 5.42 | 3.53 | 1.89 | 4.10E-04 | 4.47E-03 | 5UTR (NM_013417, exon 1 of 34, IARS) | IARS |
| chr12 | 6976694 | 6976762 | 69 | 5.78 | 4.32 | 1.46 | 4.15E-04 | 4.49E-03 | 5UTR (NM_000365, exon 1 of 7, TPI1) | TPI1 |
| chr22 | 45898740 | 45898841 | 102 | 5.59 | 4.04 | 1.56 | 4.30E-04 | 4.61E-03 | 5UTR (NM_001996, exon 1 of 15, FBLN1) | FBLN1 |
| chr8 | 99057690 | 99057784 | 95 | 6.92 | 6.04 | 0.88 | 4.34E-04 | 4.61E-03 | 5UTR (NM_000989, exon 1 of 5, RPL30) | RPL30 |
| chr11 | 63742061 | 63742216 | 156 | 7.10 | 6.26 | 0.84 | 4.43E-04 | 4.66E-03 | 5UTR (NM_004074, exon 1 of 2, COX8A) | COX8A |
| chr13 | 45915149 | 45915346 | 198 | 6.48 | 5.46 | 1.02 | 5.08E-04 | 5.31E-03 | 5UTR (NM_001286273, exon 1 of 5, TPT1) | TPT1 |
| chr3 | 23958620 | 23958744 | 125 | 6.78 | 5.86 | 0.92 | 5.39E-04 | 5.58E-03 | 5UTR (NM_002948, exon 1 of 4, RPL15) | RPL15 |
| chrX | 118602393 | 118602559 | 167 | 5.35 | 3.58 | 1.77 | 5.96E-04 | 6.07E-03 | 5UTR (NM_001152, exon 1 of 4, SLC25A5) | SLC25A5 |
| chr3 | 49066743 | 49066833 | 91 | 6.91 | 6.04 | 0.87 | 5.93E-04 | 6.07E-03 | 5UTR (NM_000884, exon 1 of 14, IMPDH2) | IMPDH2 |
| chr18 | 19192242 | 19192407 | 166 | 6.49 | 5.47 | 1.02 | 6.04E-04 | 6.10E-03 | 5UTR (NM_006938, exon 1 of 4, SNRPD1) | SNRPD1 |
| chr17 | 7210862 | 7211093 | 232 | 6.41 | 5.38 | 1.03 | 6.57E-04 | 6.59E-03 | 5UTR (NM_001970, exon 1 of 6, EIF5A) | EIF5A |
| chr2 | 24299076 | 24299323 | 248 | 6.36 | 5.33 | 1.03 | 7.19E-04 | 7.15E-03 | 5UTR (NM_016047, exon 1 of 4, SF3B6) | SF3B6 |
| chr12 | 21810646 | 21810776 | 131 | 6.69 | 5.73 | 0.95 | 7.33E-04 | 7.23E-03 | 5UTR (NM_001174097, exon 1 of 8, LDHB) | LDHB |
| chr1 | 152009378 | 152009496 | 119 | 5.62 | 4.18 | 1.44 | 7.53E-04 | 7.36E-03 | 5UTR (NM_005620, exon 1 of 3, S100A11) | S100A11 |
| chr8 | 125551355 | 125551538 | 184 | 6.33 | 5.27 | 1.06 | 7.69E-04 | 7.45E-03 | 5UTR (NM_001278645, exon 1 of 4, NDUFB9) | NDUFB9 |
| chr1 | 203830725 | 203830841 | 117 | 6.49 | 5.50 | 0.99 | 7.94E-04 | 7.63E-03 | 5UTR (NM_001304464, exon 1 of 5, SNRPE) | SNRPE |
| chr19 | 12912528 | 12912680 | 153 | 6.31 | 5.25 | 1.06 | 8.37E-04 | 7.98E-03 | 5UTR (NM_005809, exon 1 of 6, PRDX2) | PRDX2 |
| chr3 | 101405486 | 101405562 | 77 | 7.44 | 6.73 | 0.70 | 9.58E-04 | 9.07E-03 | 5UTR (NM_000986, exon 1 of 6, RPL24) | RPL24 |
| chr10 | 79793618 | 79793699 | 82 | 8.07 | 7.49 | 0.58 | 1.00E-03 | 9.41E-03 | 5UTR (NM_001142283, exon 1 of 7, RPS24) | RPS24 |
| chr16 | 56642494 | 56642630 | 137 | 6.25 | 5.22 | 1.03 | 1.11E-03 | 1.03E-02 | 5UTR (NM_005953, exon 1 of 3, MT2A) | MT2A |
| chr22 | 38071643 | 38071754 | 112 | 7.22 | 6.49 | 0.74 | 1.13E-03 | 1.05E-02 | 5UTR (NM_002305, exon 1 of 4, LGALS1) | LGALS1 |
| chr7 | 54826816 | 54826936 | 121 | 6.17 | 5.09 | 1.07 | 1.16E-03 | 1.06E-02 | 5UTR (NM_014302, exon 1 of 4, SEC61G) | SEC61G |
| chr1 | 46769334 | 46769534 | 201 | 6.34 | 5.34 | 1.01 | 1.15E-03 | 1.06E-02 | 5UTR (NM_001297566, exon 1 of 3, UQCRH) | UQCRH |
| chr5 | 179125945 | 179126121 | 177 | 7.43 | 6.74 | 0.69 | 1.20E-03 | 1.08E-02 | 5UTR (NM_001024649, exon 1 of 15, CANX) | CANX |
| chr16 | 28857545 | 28857666 | 122 | 6.20 | 5.17 | 1.03 | 1.31E-03 | 1.18E-02 | 5UTR (NM_003321, exon 1 of 10, TUFM) | TUFM |
| chr9 | 36190890 | 36191013 | 124 | 5.61 | 4.25 | 1.36 | 1.33E-03 | 1.19E-02 | 5UTR (NM_001311206, exon 1 of 4, CLTA) | CLTA |
| chr12 | 76478340 | 76478464 | 125 | 6.08 | 4.96 | 1.13 | 1.34E-03 | 1.19E-02 | 5UTR (NM_139207, exon 1 of 16, NAP1L1) | NAP1L1 |
| chr5 | 68665258 | 68665366 | 109 | 5.85 | 4.61 | 1.24 | 1.40E-03 | 1.20E-02 | 5UTR (NM_133343, exon 1 of 18, RAD17) | RAD17 |
| chr19 | 48828795 | 48828905 | 111 | 5.81 | 4.58 | 1.23 | 1.40E-03 | 1.20E-02 | 5UTR (NM_001425, exon 1 of 5, EMP3) | EMP3 |
| chr8 | 100905827 | 100905935 | 109 | 6.22 | 5.17 | 1.05 | 1.40E-03 | 1.20E-02 | 5UTR (NM_004374, exon 1 of 4, COX6C) | COX6C |
| chrX | 77359708 | 77359911 | 204 | 6.80 | 5.98 | 0.82 | 1.36E-03 | 1.20E-02 | 5UTR (NM_000291, exon 1 of 11, PGK1) | PGK1 |
| chr12 | 51632606 | 51632701 | 96 | 5.09 | 3.29 | 1.80 | 1.42E-03 | 1.21E-02 | 5UTR (NM_014764, exon 1 of 4, DAZAP2) | DAZAP2 |
| chr15 | 63449567 | 63449682 | 116 | 4.96 | 3.08 | 1.88 | 1.46E-03 | 1.23E-02 | 5UTR (NM_015920, exon 1 of 4, RPS27L) | RPS27L |
| chr6 | 35436153 | 35436252 | 100 | 6.83 | 6.02 | 0.82 | 1.46E-03 | 1.23E-02 | 5UTR (NM_007104, exon 1 of 6, RPL10A) | RPL10A |
| chrX | 135962768 | 135962894 | 127 | 5.46 | 4.00 | 1.46 | 1.49E-03 | 1.24E-02 | 5UTR (NM_001164803, exon 1 of 8, RBMX) | RBMX |
| chr17 | 16284405 | 16284518 | 114 | 6.72 | 5.87 | 0.84 | 1.49E-03 | 1.24E-02 | 5UTR (NM_018955, exon 1 of 2, UBB) | UBB |
| chr20 | 62152116 | 62152262 | 147 | 5.62 | 4.29 | 1.34 | 1.52E-03 | 1.25E-02 | 5UTR (NM_024299, exon 1 of 4, PPDPF) | PPDPF |
| chr2 | 242255316 | 242255433 | 118 | 4.81 | 2.83 | 1.99 | 1.60E-03 | 1.29E-02 | 5UTR (NM_001008492, exon 1 of 14, SEPT2) | SEPT2 |
| chr6 | 79944251 | 79944411 | 161 | 6.02 | 4.91 | 1.11 | 1.59E-03 | 1.29E-02 | 5UTR (NM_138730, exon 1 of 6, HMGN3) | HMGN3 |
| chr19 | 49122362 | 49122440 | 79 | 6.28 | 5.30 | 0.98 | 1.59E-03 | 1.29E-02 | 5UTR (NM_000979, exon 1 of 7, RPL18) | RPL18 |
| chrX | 119005790 | 119005939 | 150 | 5.39 | 3.87 | 1.52 | 1.65E-03 | 1.32E-02 | 5UTR (NM_006978, exon 1 of 1, RNF113A) | RNF113A |
| chr2 | 170681332 | 170681423 | 92 | 4.73 | 2.73 | 2.00 | 1.88E-03 | 1.50E-02 | 5UTR (NM_014168, exon 1 of 7, METTL5) | METTL5 |
| chr15 | 72523427 | 72523553 | 127 | 6.73 | 5.89 | 0.84 | 1.95E-03 | 1.54E-02 | 5UTR (NM_182470, exon 1 of 11, PKM) | PKM |
| chr1 | 156307907 | 156308090 | 184 | 6.13 | 5.11 | 1.02 | 1.97E-03 | 1.55E-02 | 5UTR (NM_001008800, exon 1 of 12, CCT3) | CCT3 |
| chr5 | 72794253 | 72794392 | 140 | 7.07 | 6.31 | 0.76 | 1.99E-03 | 1.56E-02 | 5UTR (NM_001037637, exon 1 of 6, BTF3) | BTF3 |
| chr7 | 5570134 | 5570232 | 99 | 7.80 | 7.20 | 0.60 | 2.02E-03 | 1.57E-02 | 5UTR (NM_001101, exon 1 of 6, ACTB) | ACTB |
| chr2 | 176046342 | 176046440 | 99 | 5.30 | 3.73 | 1.56 | 2.08E-03 | 1.59E-02 | 5UTR (NM_001002258, exon 1 of 4, ATP5G3) | ATP5G3 |
| chr6 | 170862277 | 170862398 | 122 | 5.49 | 4.11 | 1.38 | 2.09E-03 | 1.59E-02 | 5UTR (NM_002793, exon 1 of 6, PSMB1) | PSMB1 |
| chr17 | 73775746 | 73775861 | 116 | 5.55 | 4.22 | 1.33 | 2.08E-03 | 1.59E-02 | 5UTR (NM_005324, exon 1 of 4, H3F3B) | H3F3B |
| chr8 | 101964136 | 101964298 | 163 | 5.70 | 4.41 | 1.29 | 2.14E-03 | 1.62E-02 | 5UTR (NM_145690, exon 1 of 6, YWHAZ) | YWHAZ |
| chr5 | 132202329 | 132202410 | 82 | 5.44 | 4.04 | 1.40 | 2.20E-03 | 1.66E-02 | 5UTR (NM_014402, exon 1 of 3, UQCRQ) | UQCRQ |
| chr11 | 67798118 | 67798236 | 119 | 4.11 | 1.41 | 2.70 | 2.34E-03 | 1.74E-02 | 5UTR (NM_002496, exon 1 of 7, NDUFS8) | NDUFS8 |
| chr14 | 104387771 | 104387893 | 123 | 6.09 | 5.08 | 1.01 | 2.32E-03 | 1.74E-02 | 5UTR (NM_004894, exon 1 of 4, C14orf2) | C14orf2 |
| chr12 | 57119003 | 57119131 | 129 | 6.18 | 5.20 | 0.98 | 2.35E-03 | 1.74E-02 | 5UTR (NM_001113203, exon 1 of 11, NACA) | NACA |
| chrX | 23685591 | 23685746 | 156 | 5.82 | 4.69 | 1.14 | 2.44E-03 | 1.79E-02 | 5UTR (NM_006406, exon 1 of 7, PRDX4) | PRDX4 |
| chr17 | 42422641 | 42422729 | 89 | 4.07 | 1.41 | 2.65 | 2.47E-03 | 1.80E-02 | 5UTR (NM_002087, exon 1 of 13, GRN) | GRN |
| chr12 | 104680759 | 104680923 | 165 | 5.19 | 3.58 | 1.60 | 2.52E-03 | 1.83E-02 | 5UTR (NM_001261446, exon 1 of 14, TXNRD1) | TXNRD1 |
| chr7 | 93551353 | 93551536 | 184 | 5.39 | 4.00 | 1.39 | 2.75E-03 | 1.99E-02 | 5UTR (NM_004126, exon 1 of 2, GNG11) | GNG11 |
| chr20 | 57617766 | 57617876 | 111 | 5.55 | 4.18 | 1.36 | 2.83E-03 | 2.02E-02 | 5UTR (NM_016045, exon 1 of 6, PRELID3B) | PRELID3B |
| chr8 | 11660259 | 11660466 | 208 | 5.87 | 4.78 | 1.09 | 2.85E-03 | 2.02E-02 | 5UTR (NM_001287756, exon 1 of 5, FDFT1) | FDFT1 |
| chr12 | 16500579 | 16500680 | 102 | 5.95 | 4.89 | 1.05 | 2.83E-03 | 2.02E-02 | 5UTR (NM_001260511, exon 1 of 4, MGST1) | MGST1 |
| chr11 | 8704312 | 8704395 | 84 | 7.59 | 6.42 | 1.17 | 3.01E-03 | 2.12E-02 | 5UTR (NM_000990, exon 1 of 5, RPL27A) | RPL27A |
| chr2 | 207024061 | 207024246 | 186 | 4.98 | 3.29 | 1.69 | 3.04E-03 | 2.13E-02 | 5UTR (NM_001199981, exon 1 of 18, NDUFS1) | NDUFS1 |
| chr7 | 56174091 | 56174191 | 101 | 5.90 | 4.83 | 1.07 | 3.11E-03 | 2.15E-02 | 5UTR (NM_016139, exon 1 of 4, CHCHD2) | CHCHD2 |
| chr6 | 26158352 | 26158439 | 88 | 7.42 | 6.80 | 0.62 | 3.10E-03 | 2.15E-02 | 5UTR (NM_138720, exon 1 of 2, HIST1H2BD) | HIST1H2BD |
| chrX | 37706730 | 37706831 | 102 | 4.26 | 2.00 | 2.26 | 3.35E-03 | 2.27E-02 | 5UTR (NM_006520, exon 1 of 5, DYNLT3) | DYNLT3 |
| chr19 | 8455207 | 8455344 | 138 | 4.34 | 2.15 | 2.19 | 3.36E-03 | 2.27E-02 | 5UTR (NM_004218, exon 1 of 5, RAB11B) | RAB11B |
| chr16 | 74330699 | 74330819 | 121 | 4.86 | 3.07 | 1.79 | 3.37E-03 | 2.27E-02 | 5UTR (NM_002811, exon 1 of 7, PSMD7) | PSMD7 |

### Supplementary Table 2 (cont.)

| chr | start | end | width | Conc eFT508 | Conc PMA | log2 fold change eFT508/DMSO | p value | FDR | Deatiled Annotation | Gene |
|---|---|---|---|---|---|---|---|---|---|---|
| chr17 | 49230920 | 49231051 | 132 | 5.79 | 4.61 | 1.18 | 3.39E-03 | 2.27E-02 | 5UTR (NM_001018136, exon 1 of 8, NME1-NME2) | NME1-NME2 |
| chr2 | 201676626 | 201676745 | 120 | 6.29 | 5.40 | 0.90 | 3.30E-03 | 2.27E-02 | 5UTR (NM_001207067, exon 1 of 12, BZW1) | BZW1 |
| chr11 | 118888958 | 118889070 | 113 | 6.98 | 6.27 | 0.71 | 3.39E-03 | 2.27E-02 | 5UTR (NM_001028, exon 1 of 5, RPS25) | RPS25 |
| chr12 | 118573892 | 118574018 | 127 | 5.47 | 4.15 | 1.32 | 3.46E-03 | 2.30E-02 | 5UTR (NM_002567, exon 1 of 4, PEBP1) | PEBP1 |
| chr6 | 34204651 | 34204765 | 115 | 5.67 | 4.50 | 1.17 | 3.60E-03 | 2.38E-02 | 5UTR (NM_145903, exon 1 of 5, HMGA1) | HMGA1 |
| chr20 | 34287301 | 34287416 | 116 | 5.26 | 3.83 | 1.44 | 3.63E-03 | 2.39E-02 | 5UTR (NM_080748, exon 1 of 3, ROMO1) | ROMO1 |
| chrX | 134166366 | 134166443 | 78 | 4.88 | 3.15 | 1.74 | 3.68E-03 | 2.41E-02 | 5UTR (NM_001078171, exon 1 of 1, FAM127A) | FAM127A |
| chr2 | 27294439 | 27294539 | 101 | 5.23 | 3.73 | 1.50 | 3.75E-03 | 2.42E-02 | 5UTR (NM_001134693, exon 1 of 3, OST4) | OST4 |
| chr9 | 100745598 | 100745732 | 135 | 5.75 | 4.63 | 1.11 | 3.75E-03 | 2.42E-02 | 5UTR (NM_006401, exon 1 of 7, ANP32B) | ANP32B |
| chr6 | 74230692 | 74230757 | 66 | 6.07 | 5.07 | 0.99 | 3.75E-03 | 2.42E-02 | 5UTR (NM_001402, exon 1 of 8, EEF1A1) | EEF1A1 |
| chr12 | 16035321 | 16035587 | 267 | 5.90 | 4.87 | 1.03 | 3.94E-03 | 2.48E-02 | 5UTR (NM_007178, exon 1 of 10, STRAP) | STRAP |
| chr1 | 21835906 | 21836027 | 122 | 6.25 | 5.32 | 0.93 | 3.91E-03 | 2.48E-02 | 5UTR (NM_001127501, exon 1 of 11, ALPL) | ALPL |
| chr2 | 232329100 | 232329205 | 106 | 6.70 | 5.90 | 0.80 | 3.95E-03 | 2.48E-02 | 5UTR (NM_005381, exon 1 of 14, NCL) | NCL |
| chr6 | 26124400 | 26124517 | 118 | 6.64 | 5.86 | 0.78 | 3.87E-03 | 2.48E-02 | 5UTR (NM_003512, exon 1 of 1, HIST1H2AC) | HIST1H2AC |
| chr12 | 6643683 | 6643777 | 95 | 8.56 | 7.78 | 0.78 | 3.95E-03 | 2.48E-02 | 5UTR (NM_001289745, exon 1 of 9, GAPDH) | GAPDH |
| chr11 | 32605366 | 32605499 | 134 | 6.43 | 5.60 | 0.84 | 4.01E-03 | 2.51E-02 | 5UTR (NM_001307929, exon 1 of 8, EIF3M) | EIF3M |
| chr5 | 138089109 | 138089225 | 117 | 5.42 | 4.11 | 1.31 | 4.24E-03 | 2.63E-02 | 5UTR (NM_001903, exon 1 of 18, CTNNA1) | CTNNA1 |
| chr9 | 75766780 | 75766861 | 82 | 6.43 | 5.61 | 0.82 | 4.30E-03 | 2.66E-02 | 5UTR (NM_000700, exon 1 of 13, ANXA1) | ANXA1 |
| chr5 | 41870390 | 41870631 | 242 | 5.06 | 3.47 | 1.59 | 4.36E-03 | 2.68E-02 | 5UTR (NM_000436, exon 1 of 17, OXCT1) | OXCT1 |
| chr4 | 77997067 | 77997155 | 89 | 5.34 | 4.00 | 1.34 | 4.42E-03 | 2.69E-02 | 5UTR (NM_006835, exon 1 of 7, CCNI) | CCNI |
| chr9 | 21994263 | 21994418 | 156 | 6.17 | 5.25 | 0.92 | 4.40E-03 | 2.69E-02 | 5UTR (NM_058195, exon 1 of 3, CDKN2A) | CDKN2A |
| chr3 | 120169740 | 120169849 | 110 | 6.57 | 5.73 | 0.83 | 4.44E-03 | 2.69E-02 | 5UTR (NM_007085, exon 1 of 11, FSTL1) | FSTL1 |
| chr9 | 117350039 | 117350153 | 115 | 5.31 | 3.87 | 1.44 | 4.69E-03 | 2.83E-02 | 5UTR (NM_004888, exon 1 of 3, ATP6V1G1) | ATP6V1G1 |
| chr8 | 101734088 | 101734316 | 229 | 6.88 | 6.18 | 0.70 | 4.75E-03 | 2.85E-02 | 5UTR (NM_002568, exon 1 of 15, PABPC1) | PABPC1 |
| chr9 | 34637567 | 34637791 | 225 | 5.32 | 3.96 | 1.37 | 4.80E-03 | 2.86E-02 | 5UTR (NM_001282206, exon 1 of 4, SIGMAR1) | SIGMAR1 |
| chr14 | 69864918 | 69865051 | 134 | 5.87 | 4.85 | 1.03 | 4.81E-03 | 2.86E-02 | 5UTR (NM_004450, exon 1 of 4, ERH) | ERH |
| chr11 | 65430326 | 65430387 | 62 | 4.30 | 2.15 | 2.15 | 4.91E-03 | 2.87E-02 | 5UTR (NM_001243984, exon 1 of 11, RELA) | RELA |
| chr13 | 76111862 | 76111986 | 125 | 5.23 | 3.83 | 1.40 | 4.91E-03 | 2.87E-02 | 5UTR (NM_001287392, exon 1 of 4, COMMD6) | COMMD6 |
| chr3 | 99357438 | 99357528 | 91 | 6.07 | 5.13 | 0.94 | 4.89E-03 | 2.87E-02 | 5UTR (NM_020351, exon 1 of 4, COL8A1) | COL8A1 |
| chr4 | 83295011 | 83295160 | 150 | 7.08 | 6.42 | 0.66 | 4.92E-03 | 2.87E-02 | 5UTR (NM_001003810, exon 1 of 7, HNRNPD) | HNRNPD |
| chr6 | 38670817 | 38670921 | 105 | 5.05 | 3.53 | 1.52 | 5.08E-03 | 2.95E-02 | 5UTR (NM_006708, exon 1 of 6, GLO1) | GLO1 |
| chr13 | 100153679 | 100153842 | 164 | 6.09 | 5.17 | 0.92 | 5.13E-03 | 2.96E-02 | 5UTR (NM_004800, exon 1 of 17, TM9SF2) | TM9SF2 |
| chr17 | 79479719 | 79479828 | 110 | 7.99 | 7.50 | 0.50 | 5.35E-03 | 3.07E-02 | 5UTR (NM_001199954, exon 1 of 6, ACTG1) | ACTG1 |
| chr14 | 105941150 | 105941244 | 95 | 5.17 | 3.73 | 1.43 | 5.41E-03 | 3.09E-02 | 5UTR (NM_001312, exon 1 of 8, CRIP2) | CRIP2 |
| chr11 | 64889532 | 64889660 | 129 | 7.37 | 6.77 | 0.60 | 5.61E-03 | 3.19E-02 | 5UTR (NM_001997, exon 1 of 5, FAU) | FAU |
| chr2 | 9771028 | 9771144 | 117 | 4.63 | 2.83 | 1.80 | 5.75E-03 | 3.20E-02 | 5UTR (NM_006826, exon 1 of 6, YWHAQ) | YWHAQ |
| chr6 | 160183040 | 160183179 | 140 | 5.21 | 3.83 | 1.38 | 5.68E-03 | 3.20E-02 | 5UTR (NM_005891, exon 1 of 9, ACAT2) | ACAT2 |
| chr14 | 90863371 | 90863467 | 97 | 5.32 | 4.00 | 1.32 | 5.75E-03 | 3.20E-02 | 5UTR (NM_006888, exon 1 of 6, CALM1) | CALM1 |
| chr3 | 128902587 | 128902776 | 190 | 6.06 | 5.11 | 0.95 | 5.65E-03 | 3.20E-02 | 5UTR (NM_001127194, exon 1 of 5, CNBP) | CNBP |
| chr1 | 155990641 | 155990741 | 101 | 6.16 | 5.27 | 0.89 | 5.76E-03 | 3.20E-02 | 5UTR (NM_003145, exon 1 of 6, SSR2) | SSR2 |
| chr1 | 11072713 | 11072821 | 109 | 4.86 | 3.22 | 1.64 | 5.83E-03 | 3.21E-02 | 5UTR (NM_007375, exon 1 of 6, TARDBP) | TARDBP |
| chr1 | 225965525 | 225965597 | 73 | 6.75 | 5.97 | 0.78 | 5.84E-03 | 3.21E-02 | 5UTR (NM_001130440, exon 1 of 4, SRP9) | SRP9 |
| chr5 | 43313411 | 43313514 | 104 | 6.63 | 5.85 | 0.78 | 5.90E-03 | 3.23E-02 | 5UTR (NM_001098272, exon 1 of 11, HMGCS1) | HMGCS1 |
| chr14 | 54863701 | 54863816 | 116 | 4.07 | 1.83 | 2.24 | 6.06E-03 | 3.31E-02 | 5UTR (NM_005192, exon 1 of 8, CDKN3) | CDKN3 |
| chrX | 20159887 | 20159969 | 83 | 5.00 | 3.47 | 1.53 | 6.10E-03 | 3.32E-02 | 5UTR (NM_001412, exon 1 of 7, EIF1AX) | EIF1AX |
| chr19 | 39109730 | 39109938 | 209 | 5.23 | 3.87 | 1.35 | 6.45E-03 | 3.48E-02 | 5UTR (NM_001308393, exon 1 of 8, EIF3K) | EIF3K |
| chr14 | 23057989 | 23058139 | 151 | 5.46 | 4.25 | 1.20 | 6.46E-03 | 3.48E-02 | 5UTR (NM_001344, exon 1 of 3, DAD1) | DAD1 |
| chr5 | 85913746 | 85913887 | 142 | 6.40 | 5.60 | 0.81 | 6.61E-03 | 3.54E-02 | 5UTR (NM_001867, exon 1 of 3, COX7C) | COX7C |
| chr14 | 70233831 | 70233988 | 158 | 5.00 | 3.47 | 1.53 | 6.70E-03 | 3.56E-02 | 5UTR (NM_001039465, exon 1 of 8, SRSF5) | SRSF5 |
| chr17 | 56429431 | 56429563 | 133 | 5.25 | 3.91 | 1.33 | 6.70E-03 | 3.56E-02 | 5UTR (NM_003168, exon 1 of 5, SUPT4H1) | SUPT4H1 |
| chr16 | 67880810 | 67880924 | 115 | 4.53 | 2.73 | 1.80 | 6.79E-03 | 3.58E-02 | 5UTR (NM_005796, exon 1 of 5, NUTF2) | NUTF2 |
| chr12 | 6602237 | 6602432 | 196 | 5.42 | 4.22 | 1.20 | 6.77E-03 | 3.58E-02 | 5UTR (NM_016497, exon 1 of 3, MRPL51) | MRPL51 |
| chr1 | 19578289 | 19578369 | 81 | 5.05 | 3.58 | 1.47 | 6.88E-03 | 3.61E-02 | 5UTR (NM_016183, exon 1 of 8, MRTO4) | MRTO4 |
| chr7 | 155089588 | 155089736 | 149 | 5.68 | 4.61 | 1.07 | 7.07E-03 | 3.70E-02 | 5UTR (NM_198337, exon 1 of 4, INSIG1) | INSIG1 |
| chr15 | 85259228 | 85259388 | 161 | 5.87 | 4.91 | 0.96 | 7.22E-03 | 3.76E-02 | 5UTR (NM_001271918, exon 1 of 7, SEC11A) | SEC11A |
| chr1 | 36107013 | 36107154 | 142 | 5.75 | 4.73 | 1.01 | 7.32E-03 | 3.79E-02 | 5UTR (NM_002794, exon 1 of 6, PSMB2) | PSMB2 |
| chr7 | 91763615 | 91763795 | 181 | 6.89 | 6.23 | 0.66 | 7.50E-03 | 3.87E-02 | 5UTR (NM_000786, exon 1 of 10, CYP51A1) | CYP51A1 |
| chr13 | 48807287 | 48807419 | 133 | 4.56 | 2.83 | 1.74 | 7.63E-03 | 3.92E-02 | 5UTR (NM_021999, exon 1 of 6, ITM2B) | ITM2B |
| chr3 | 156272840 | 156272937 | 98 | 5.35 | 4.08 | 1.27 | 7.68E-03 | 3.93E-02 | 5UTR (NM_007107, exon 1 of 5, SSR3) | SSR3 |
| chr19 | 1095245 | 1095373 | 129 | 5.39 | 4.18 | 1.20 | 8.03E-03 | 4.09E-02 | 5UTR (NM_001316324, exon 1 of 8, POLR2E) | POLR2E |
| chrX | 102611399 | 102611563 | 165 | 4.34 | 2.41 | 1.93 | 8.29E-03 | 4.20E-02 | 5UTR (NM_001006612, exon 1 of 3, TCEAL9) | TCEAL9 |
| chr1 | 23885893 | 23886018 | 126 | 4.84 | 3.29 | 1.55 | 8.42E-03 | 4.23E-02 | 5UTR (NM_002167, exon 1 of 3, ID3) | ID3 |
| chr1 | 111992071 | 111992214 | 144 | 5.42 | 4.25 | 1.17 | 8.38E-03 | 4.23E-02 | 5UTR (NM_001688, exon 1 of 7, ATP5F1) | ATP5F1 |
| chr12 | 14923996 | 14924065 | 70 | 5.17 | 3.83 | 1.35 | 8.62E-03 | 4.32E-02 | 5UTR (NM_175054, exon 1 of 1, HIST4H4) | HIST4H4 |
| chr2 | 237994534 | 237994635 | 102 | 4.10 | 2.00 | 2.11 | 8.86E-03 | 4.37E-02 | 5UTR (NM_198189, exon 1 of 9, COPS8) | COPS8 |
| chr1 | 173793805 | 173794060 | 256 | 4.15 | 2.15 | 2.00 | 8.83E-03 | 4.37E-02 | 5UTR (NM_018122, exon 1 of 17, DARS2) | DARS2 |
| chr4 | 109571757 | 109571851 | 95 | 4.59 | 2.91 | 1.68 | 8.81E-03 | 4.37E-02 | 5UTR (NM_021227, exon 1 of 4, OSTC) | OSTC |
| chr14 | 50087247 | 50087354 | 108 | 5.06 | 3.58 | 1.48 | 8.75E-03 | 4.37E-02 | 5UTR (NM_001001, exon 1 of 2, RPL36AL) | RPL36AL |
| chr8 | 62626817 | 62627150 | 334 | 6.08 | 5.20 | 0.87 | 8.96E-03 | 4.40E-02 | 5UTR (NM_001164755, exon 1 of 13, ASPH) | ASPH |
| chr10 | 73975770 | 73975919 | 150 | 4.84 | 3.29 | 1.56 | 9.49E-03 | 4.64E-02 | 5UTR (NM_001198798, exon 1 of 10, ASCC1) | ASCC1 |
| chr1 | 17307044 | 17307154 | 111 | 5.02 | 3.58 | 1.44 | 9.64E-03 | 4.69E-02 | 5UTR (NM_001135248, exon 1 of 9, MFAP2) | MFAP2 |
| chr17 | 8152604 | 8152682 | 79 | 4.19 | 2.29 | 1.90 | 9.78E-03 | 4.72E-02 | 5UTR (NM_012393, exon 1 of 28, PFAS) | PFAS |
| chr22 | 42084771 | 42084899 | 129 | 5.22 | 3.96 | 1.27 | 9.76E-03 | 4.72E-02 | 5UTR (NM_005008, exon 1 of 4, SNU13) | SNU13 |
| chr4 | 122617984 | 122618156 | 173 | 5.73 | 4.73 | 1.00 | 9.86E-03 | 4.74E-02 | 5UTR (NM_001154, exon 1 of 13, ANXA5) | ANXA5 |
| chrX | 71497006 | 71497098 | 93 | 5.78 | 4.80 | 0.97 | 1.01E-02 | 4.83E-02 | 5UTR (NM_001007, exon 1 of 7, RPS4X) | RPS4X |
| chr17 | 5342322 | 5342484 | 163 | 5.91 | 5.02 | 0.89 | 1.02E-02 | 4.84E-02 | 5UTR (NM_001212, exon 1 of 6, C1QBP) | C1QBP |
| chr10 | 73610916 | 73611054 | 139 | 6.17 | 5.37 | 0.80 | 1.02E-02 | 4.85E-02 | 5UTR (NM_001042465, exon 1 of 15, PSAP) | PSAP |
| chr19 | 39897487 | 39897562 | 76 | 6.97 | 6.30 | 0.67 | 1.02E-02 | 4.85E-02 | 5UTR (NM_003407, exon 1 of 2, ZFP36) | ZFP36 |
| chr12 | 42719796 | 42719942 | 147 | 4.27 | 2.41 | 1.86 | 1.06E-02 | 4.99E-02 | 5UTR (NM_033114, exon 1 of 8, ZCRB1) | ZCRB1 |

#### capCLIP Reagent List

| enzyme/reagent | vendor | cat # |
|---|---|---|
| DNAse I | NEB | M0303S |
| RNAse inhibitor, recombinant | NEB | M0314S |
| T4 PNK | NEB | M0201S |
| T4 RNA ligase, high conc. | NEB | M047M |
| proteinase K | NEB | P8107S |
| NEBNext Ultra II Q5 Master Mix | NEB | M0544S |
| SuperScript IV RT | ThermoFisher | 18090050 |
| dNTP set, 100 mM | ThermoFisher | 10297018 |
| Bolt 4-12% Bis-Tris Plus Gels, 10 well | ThermoFisher | NW04120BOX |
| NuPAGE LDS Sample Buffer | ThermoFisher | NP0007 |
| NuPAGE MES SDS Run buf, 20X | ThermoFisher | NP0002 |
| NuPAGE reducing agent (10X) | ThermoFisher | NP0009 |
| NuPAGE transfer buf, 20X | ThermoFisher | NP0006 |
| Roche EDTA-free Complete protease tablets, 20 tablets | SigmaAldrich | 11873580001 |
| anti-FLAG mAb, clone M2 | SigmaAldrich | F3165 |
| GlycoBlue Coprecipitant, 15 mg/ml | SigmaAldrich | AM9515 |
| Sephadex G-25 spin columns | SigmaAldrich | 11273990001 |
| X-Cell SureLock Mini-Cell | ThermoFisher | EI0001 |
| X-cell II Blot Module | ThermoFisher | EI9051 |
| Ambion RNAse I | ThermoFisher | AM2294 |
| Dynabeads protein G | ThermoFisher | 10003D |
| Acid-Phenol pH 4.5 | ThermoFisher | AM9712 |
| phiX174 DNA/BsuRI (HaeIII) Marker | ThermoFisher | SM0251 |
| Corning® Costar® Spin-X centrifuge tube filters | ThermoFisher | CLS8160 |
| 21G syringe needles, 1.5 inch | BD | 305167 |
| bis-acrylamide, acrylamide, urea | SigmaAldrich | |
| Exo-SAP-IP | ThermoFisher | 78200 |
| MyONE Silane Dynabeads | ThermoFisher | 37002D |
| empty Bolt gel cassettes for homemade gel mixes | ThermoFisher | |
| SYBR gold stain | ThermoFisher | |

| other equipment + reagents |
|---|
| Stratagene Stratalinker 2400 |
| Eppendorf Thermomixer |
| Qubit and Qubit reagents |
| Agilent or Tapestation +disposables |
| Illlumina NextSeq +reagents |

**RNA and DNA oligonucleotides**

**SR_RNA** 3' 21 nt RNA adapter 5’-P-AGAUCGGAAGAGCACACGUCU-SpC3-3’

**SR-RT** 21 nt DNA RT primer. 3’-TCTAGCCTTCTCGTGTGCAGA-5’

**5’SRdeg_DNAv2** DNA adapter (blocked 3’ end)
5’P-HHNNNNNNNNGATCGTCGGACTGTAGAACTCTGAAC-3SpC-3’

**Illumina RP1 primer**
5’-AATGATACGGCGACCACCGAGATCTACACGTTCAGAGTTCTACAGTCCG**s**A -3’

**NEBNext index primers (index hexamer in blue)**
3’-TsCTAGCCTTCTCGTGTGCAGACTTGAGGTCAGTGNNNNNNTAGAGCATACGGCAGAAGACGAAC-5’

#### Buffers/Solutions:

**1X PXL**
1X PBS (TC grade; no $Mg^{2+}$, $Ca^{2+}$)
0.1% SDS
0.5% deoxycholate
0.5% NP-40 (Igepal)

**5X PXL (5X in Na+, not detergents)**
5X PBS (TC; no $Mg^{2+}$, no $Ca^{2+}$)
0.1% SDS
0.5% deoxycholate
0.5% NP-40 (Igepal)

**1X PNK Buffer**
50 mM Tris-Cl pH 7.5
10 mM $MgCl_2$
0.5% NP-40 (Igepal)

**1X PNK+EGTA**
50 mM Tris-Cl pH 7.5
20 mM EGTA
0.5% NP-40 (Igepal)

**2X PK buffer**
200 mM Tris-Cl pH 7.5
100 mM NaCl
20 mM EDTA

**Nucleic acid elution buffer**
1 M NaOAc pH 5.2
1 mM EDTA

**1X PBS (tissue culture grade; no $Mg^{2+}$, no $Ca^{2+}$)**

**1X PBS-Tw (1X PBS + 0.05% Tween-20)**

#### A: Preparation of cells, drug treatment, UV cross-linking and cell harvesting

For clarity, this protocol will describe the 'eFT-508 capCLIP' experiment from the paper. As future users will be applying the method to a diverse range of cells and/or tissues, some of the steps of the method will need to be adapted to the system under study. Fox example, UV crosslinking dose, and RNAse I concentration and/or time of digestion typically require adjustment for the individual cells or tissue being used. We have tried to document the steps that require individual adjustment, and how to determine what modification(s) need to be made.

##### A1. Cell growth and plating

Flag-eIF4E Hela cells were grown in 10% FBS and DMEM and allowed to grow to ~85% confluency in 150 mm diameter tissue culture plates. The cells were then serum starved overnight (~15 hours) in 9 ml of DMEM.

##### A2. Drug treatments

In the morning, aliquots with 3 µl of a 1 mM solution of eFT-508 in DMSO and 997 µl of DMEM were prepared for each plate being treated with drug. Aliquots with 3 µl of DMSO and 997 µl of DMEM were prepared for each control plate.

After addition of a single 1ml aliquot per plate, plates were placed back in the TC incubator for 1 hour. Aliquots of 5.5 µl of a 1 mM stock solution of PMA (phorbol 12-myristate 13-acetate) and 994.5 µl of DMEM were prepared for each plate, and at the end of pre-treatment, each 1ml aliquot was added per plate and the plates returned to the TC incubator for another 30 mins.

##### A3. UV irradiation and harvesting

Pre-treatment with eFT-508 or DMSO only, and subsequent stimulation with PMA was sequenced so that 2 150 mm plates are ready approximately every 20 minutes. For this experiment, each 'replicate' sample represents the cells from 2 150 mm plates, and thus the two plates for each sample were processed together.

The 2 paired plates are brought out of the incubator and placed in a tray filled with an ice and water slurry so that the plates are essentially floating. ***This is to ensure that the plates are as level as possible during irradiation, and that the cells remain covered in PBS.***

The media on each plate is aspirated and 6 ml of ice-cold 1X PBS then added back to each plate- just enough volume to cover the cells.

Plates are irradiated for a total of 175 mJ/cm2, then rotated 180˚ and irradiated for an additional 175 mJ/cm2. ***This is to ensure that the irradiation from the individual UV bulbs irradiates the cells as evenly as possible.*** Plates are then transferred to the benchtop (blotted dry on the bottom) and are then are very gently scraped with a Teflon scraper to allow collection of the detached cells.

This initial cell suspension from both plates is transferred to a single chilled 50 ml tube. An additional 10 ml of ice cold 1X PBS is then added to each plate, briefly scraped again, and these volumes also added to the same 50 ml tube.

The 50ml tube is spun at 350 x G for 2 minutes at 4˚C, the supernatant gently removed (poured if possible), and the cell pellet resuspended in 1ml of 1X PBS. This cell suspension is then transferred to an Eppendorf tube, spun at 400 x G for 1 min, and the supernatant again removed by pipette.

The cell pellet volume is recorded for each sample and pellets stored at -80˚C.

#### B. Immunoprecipitation

There are 4 samples to be worked up, each consisting of cell pellets collected from 2 150 mm plates of flag-eIF4E Hela cells:

| | |
|---|---|
| PMA stimulation only (1) | PMA 1 |
| PMA stimulation only (2) | PMA 2 |
| eFT508 inhibitor (1) | eFT508 1 |
| eFT508 inhibitor (2) | eFT508 2 |

##### B1. Antibodies

Sigma anti-FLAG mouse mAb clone M2 IgG1
(stock concentration is 3.85 mg/ml)

##### B2. Bead prep

**Note:** all washes are performed with ice-cold buffers, 1mL per wash unless otherwise indicated

For each IP we use 50 µl of protein G Dynabeads

The binding capacity is ~8 µg Ab per mg beads. 30 mg beads/ml is stock, so 50 µl of beads should bind to 12 µg of antibody.

1. There are a total of 4 IPs:
2. Prepare beads; aliquot 220 µl of bead stock to fresh Eppie (50 µl beads/IP + 10% for pipetting errors)

3. Immobilise beads on magnetic rack; take off storage buffer
4. Take out of magnetic rack; resuspend beads 1X PBS-Tw
5. Wash a total of 3X in 1X PBS-Tw
6. After final wash, resuspend beads in 200 µl of PBS-Tw
7. For FLAG mAb, use 12 µg per 50 µl of beads, so 52.8 µg/220 µl beads; at stock concentration of 3.85 µg/µl, add 13.7 µl of FLAG Ab to washed beads and gently mix
8. Place tube on rotating rack at room temp for ~45 min.
9. Wash beads 3x with 1 PXL, with beads in final wash; split 1ml volume across 4 new Eppies
10. Keep beads in final wash until ready to perform the IP

#### B3. Crosslinked lysate workup

1. Resuspend each cell pellet (representing 2 x 150 mm plates of crosslinked cells in 500 µl 1X PXL + protease inhibitors. (Prepare beforehand 10 ml of 1X PXL supplemented with EDTA-free Complete protease inhibitors) ***Depending on the cell type, sometimes a volume closer to 800-1000 µl of 1X PXL is more appropriate for lysis.***
2. Take out 20 µl from each sample and retain on ice for Bradford analysis, and for possible future analysis or troubleshooting.
3. Let samples sit on ice for 15 min. (Vortex occasionally) ***Samples should get quite viscous from the lysis of nuclei and the release of genomic DNA. If this is not apparent at this point, you may need to let samples sit on ice a bit longer.***
4. Triturate samples by passing them through a 21G needle 5 times.
5. Add 40 µl of DNAse I (NEB M0303) to each tube.
6. Incubate at 37° and 750 rpm in the Thermomixer for 10 min, then return samples to ice for at least 1 min. ***Make sure the viscosity of the sample has been significantly reduced. Strangely, sometimes full lysis of nuclei takes a long time and the viscosity of the genomic DNA can take a while to fully develop.*** If still viscous, add additional DNAse.
7. ; For the RNase step, make a 1:50 dilution of the RNase 1 stock (AM2295) in 1X PBS.
8. Add 5 µl of **diluted** RNase to each sample (1 µl per 100 µl of lysate)
9. Incubate at 37° and 750 rpm for 5 min in the Thermomixer, then return samples to ice for at least 1 min. Spin lysates in pre-chilled micro-centrifuge; full speed for 30 min at 4°.
10. After the spin, carefully take the cleared supernatant of each sample to a fresh Eppie. ***We generally store the insoluble pellets at -20˚ or -80˚ for possible troubleshooting of any failed experiments.***
11. While lysates are spinning, do Bradfords (or other protein concentration assay) on 2 µl of the 20 µl samples. The ideal concentration for IP is ~1.5 µg/µl. If the concentration is above this amount, calculate the amount of 1X PXL to add to obtain a final protein concentration of 1.5 µg/µl, and then add this amount to the lysate. If your concentration is less than 1.5 µg/µl, then go with what you have.

#### B4. Immunoprecipitation

1. Once you are ready for the IP, immobilise the FLAG mAb coated beads you prepared in **Step 10 of Section B2** in the magnetic rack, and take off the 1X PXL wash buffer.
2. Add the concentration-adjusted, cleared supernatant from **Step 11 of Section B3** to the FLAG mAb coated beads and gently mix.
3. Place the IP samples on a rotating rack for 2 hours at 4°.

4. Immobilise the beads from the IP samples on the magnetic rack.
5. Collect the bead supernatants and save them in fresh Eppies. Store at -80˚C. ***These post-IP supernatants are generally analysed by western to determine the efficiency of the IP step.***
6. Wash the IP beads 2X with 1X PXL
7. Then wash the IP beads 2X with 5X PXL
8. Then wash the IP beads 2X with 1X PNK
9. Immobilise the beads in the magnetic rack and emove this final wash from the beads.

**B5. 3’ end dephosphorylation**

***This step is to resolve the possible 2’-3’ cyclic phosphate that is present on the 3’ end of the RNA fragments into 2’ and 3’ -OH. The cyclic phosphate is created by the addition of the RNase I enzyme. (Fragments may also be produced by endogenous RNases.)***

Resuspend each set of IP beads in the following mix:

8 µl 10x T4 PNK buffer
4 µl T4 PNK
2 µl NEB RNAse inhibitor
66 µl water
80 µl

***Generally, we make a master mix of these components, and then distribute 80 µl of the mix to each sample.***

Incubate in thermomixer at 37° and 750 rpm for 20 min.

Wash each set of IP beads:
- 1X with 1X PNK
- 1X with PNK+EGTA
- 2X with 1X PNK

After last wash, immobilise the beads again in the magnetic rack, take off all excess wash buffer and proceed to **Section B7**.

**B6. 3’ linker labelling**

***This step is usually done during the 2-hour IP step.***

Set up the following reaction:

65 µl water
5 µl NEB RNAse inhibitor
10 µl 10X T4PNK buffer
5 µl 3' **SR_RNA(-P) adapter** (20 µM; 100 pmol); final [adapter] is 1 µM
5 µl $^{32}$P-gamma-ATP (150 mCi/ml, 25 µM; 300 pmol); final [ATP] is 10 µM
10 µl T4 PNK
100 µl total

1. Incubate at 37° for 30 min.
2. Add 5 µl 3 mM ‘cold’ (non-radioactive) ATP, and then incubate a further 1 min at 37°
3. Prepare a mini G-25 column. Resuspend the resin in the G-25 column by vortexing, then break off the bottom seal and loosen the cap.
4. Pre-centrifuge the column in a collection tube 1 min at 735 x g at room temperature (so ~3,000 rpm in a typical Eppendorf centrifuge).

5. Transfer the G-25 column to a fresh, RNase-free micro-centrifuge tube.
6. Add 100 µl of 1X PNK buffer to the top of the column, and then spin again at 735 x g for 1 min.
7. Move the column to a fresh collection tube
8. Carefully apply the phosphorylated linker sample to the top of the column and spin the column for 2 min at 735g
9. Use your P200 pipet to estimate the volume of the eluate and bring the volume up to 105 µl with MQ water if needed.

**B7. 3' linker ligation (on-bead)**

***The SR_RNA primer now has a phosphate (hot or cold) at its 5' end, so it can now be ligated on to the 3' end of the RNA fragments pulled down by the flag IP. The 3' end of this RNA adapter is 'blocked' so that this 3' end is not a substrate for adapter ligation itself.***

Resuspend each set of IP beads from **Section B7** in a mix of:

| | | |
|---|---|---|
| 4 | µl | 10X RNA ligase buffer |
| 9 | µl | 50% PEG 8000 |
| 10 | µl | $^{32}$P- **SR_RNA(now +$^{32}$P) RNA adapter** (@ 1 pmol/µl) |
| 0.5 | µl | RNAse inhibitor |
| 4 | µl | 10 mM ATP |
| 1 | µl | T4 RNA ligase HC |
| 11.5 | µl | water |
| 40 | µl | total |

***Again, we would generally make a master mix of this ligation mix for all samples and then aliquot 40 µl per set of beads.***

Incubate in the Thermomixer at room temp and at 750 rpm for approximate **9.5 hours (overnight)**

#### C. SDS-PAGE and nitrocellulose transfer

**C1. Preparation of samples for gel; SDS-PAGE**

1. Place the beads + ligation mix Eppies on the magnetic rack and immobilise the beads. Carefully remove the radioactive reaction mix and dispose in the hot waste.
2. Wash the beads 1X with 1X PXL
3. Wash the beads 1X with 5X PXL
4. Wash the beads 1X with 1X PNK
5. Wash the beads 3X with 1X PNK+EGTA
6. In the last wash step, transfer beads to a new tube. ***As the next step will use LDS and a reducing agent to pull the flag-eIF4E and RNA off the beads, we don't want to also solubilise any material that has precipitated overnight on to the tube itself.***
7. Resuspend each set of beads in 40 µl 1X "LDS/β-me sample buffer"

   1ml of 1X buffer = 250 µl 4X Bolt LDS Novex sample buffer
   10 µl β-mercapto-ethanol, neat
   740 µl MQ water

8. Incubate at 70° at 1200 rpm for 10 min in Thermomixer

9. Immobilise beads and move the eluted sample to a fresh Eppie
10. Set up a X-Cell SureLock Mini-Cell (or Bolt gel apparatus) with a 10-well Bolt 4-12% Bis-Tris Plus gel. We generally use the MES/SDS running buffer, not the MOPS/SDS buffer.
11. Load entire 40 µl sample in a well of the gel
12. Load an appropriate pre-stained protein molecular weight marker into the outside lanes of the gel
13. Run gel @ 150 V constant until all dyes and salt fronts have left the gel

**C2. Nitrocellulose transfer of Bolt gel**

1. After the gel run, carefully dispose of the radioactive running buffer and give the gel itself a rinse with MQ water.
2. Crack open the gel cassette, hopefully retaining the gel intact on one of the plastic surfaces of the cassette.
3. Carefully cut off the wells at top and the lip and extreme lower region of the gel. ***This is best done with a blunter edge; we like to use the thinner of the glass sheets that is used in the Biorad mini-gel system. You can make a clean 'chop' across the entire gel. We find that when using scalpel blades or straight razor blades it is difficult to achieve clean cuts.***
4. Make up ~500 ml of 1X NuPage transfer buffer (with 10% methanol) to use with the X-cell II Blot Module.
5. Pre-cut Whatman paper and a single nitrocellulose membrane to fit the X-cell II Blot Module. ***Make sure the blotting pad/sponges use are using are completely clean.***
6. Assemble the blot on the deep plate of the blot module in the following order:

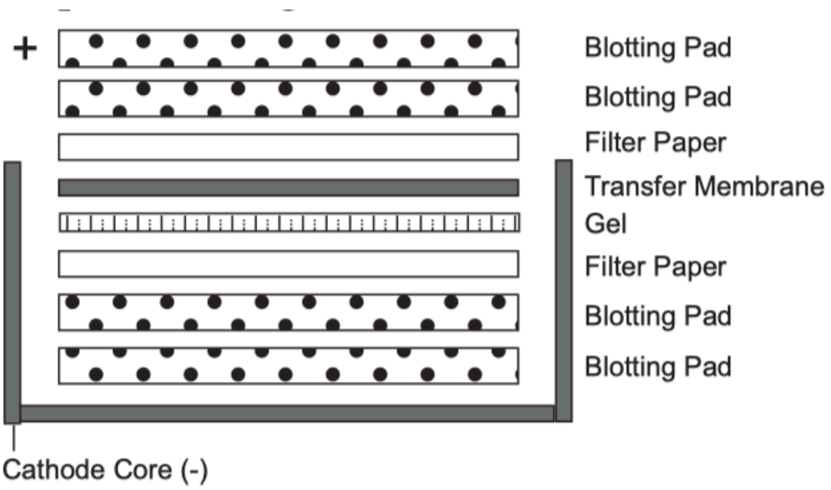

7. Place the anode plate on top and assemble the module into the X-Cell SureLock Mini-Cell tank and fill the X-cell with transfer buffer (no higher than the blot/sponges). Then add DI water to the tank itself, to dissipate heat from the X-cell itself.
8. Run the transfer for 1 hour at 30 V constant; amperage is ~ 170 mA starting
9. After 30 min take apart the apparatus and carefully unstack the sponges, etc. Check transfer by just peeling back the membrane from the gel; make sure that the pre-stained marker bands have transferred properly
10. Let the membrane dry a bit on top of a Kimwipe, then carefully attach the membrane to an A-5 sized piece of Whatman paper using with a tiny bit of tape. ***As you will want to be able to align the radioactive bands you will see with the phosphorimager with the membrane itself, we generally use some source of dilute $^{32}P$ (It is a good idea to save the $1^{st}$ wash collected after ligation for this purpose) to make a pattern of radioactive 'spots' on the***

***Whatman paper. As you need a way to see these spots on the Whatman paper itself, 1st make a pattern of spots with a sharpie, and then spot 1-2 µl of the dilute $^{32}P$ solution precisely on top of the Sharpie marks. Getting the radioactive intensity of the spots in the same range and the radioactive bands on the membrane takes a bit of practice, but you are likely to expose the membrane for 24-48 hours on the phosphorimager plate.***

11. Carefully cover the Whatman sheet assembly with glad wrap and secure the assembly inside an imaging cassette with tape. Add in a freshly blanked phosphorimager screen and 1st do an overnight 'test' exposure to gauge how long you will need to your 'real' exposure.
12. While a sensitive Geiger counter should be able to detect a signal from the membrane, it is fairly unusual that the membrane has sufficient radioactivity still present to require shielding. Generally one can now handle the membrane without any sort of protective screen, which makes the excision of bands in the next section of the protocol much easier to do.

#### C3. Phosphorimaging of the nitrocellulose membrane

Process the 'overnight' phosphorimaging screen. Ideally, one can already see in the 12-16 hour scan the presence of a radioactive band or 'smear' in the region of 30 kDa – 50 kDa for flag-eIF4E (this range will of course depend on the molecular weight of the cap-binding protein under study, and the precise amount of RNase used).

Also, when setting up capCLIP it is critical to do the following controls:

- a no UV crosslinking sample
- high and low concentrations of RNase
- a mock IP with an irrelevant antibody
- a sample from cells with (ideally) no epitope-tagged eIF4E (or other cap-binding protein) [this sample should be UV crosslinked and treated with RNase]

Such controls will allow you to confirm that the radioactive bands you see are UV-dependent, dependent on the presence of the epitope-tagged eIF4E protein, and dependent on the presence of the correct anti-epitope antibody on the protein G Dynabeads

Once you have determined the least amount of time you require to produce a phosphorimage with good band detail, expose the screen for this amount of time and process the image, making sure that you can also resolve your $^{32}P$ 'marker spots' with good detail too.

Finally, you will need to determine the print settings that allow you to get a 1:1-sized version of your phosphorimage (that is the print needs to be exactly the same size as the original nitrocellulose membrane). Once you determine the proper settings, you will need to print the image on a sheet of "transparency film" (or what used to be called 'overhead transparencies' as they were used to project lecture notes, etc. using overhead projectors). You can still find this material at most office supply stores, and these sheets should all be compatible with modern multifunction copy-printer devices. Add a sheet of the transparency film to the bypass tray of your multifunction printer and send your image there to produce the 'X-ray film equivalent' you need to cut out RNA-protein bands from the membrane. ***Of course, if you still use X-ray film to do $^{32}P$ imaging, you can skip all of this. Indeed, you can even skip the 'spotting' steps and just use something like a Promega luminescent maker ladder.***

#### D. Cutting of RNA-protein bands from nitrocellulose; RNA isolation & purification

##### D1. Cutting out eIF4E:RNA bands from the nitrocellulose membrane

If everything has gone OK you have your membrane still attached to a piece of Whatman paper that has the Sharpie/$^{32}$P marks that will allow you to align the transparency precisely to the membrane. It is usually a good idea to tape the entire Whatman/membrane sheet to the bench, and then once you have positioned the transparency on top, you can tape this sheet in place too.

As the transparency sheeting is usually fairly thin, we like to leave the sheet in place as we cut the membrane; this more or less ruins the transparency as a record of the experiment, but allows you cut the membrane in exactly the right place.

While there is no hard or fast rule, we are looking to excise eIF4E:RNA material corresponding to crosslinked RNAs from ~30-80 nt. As the material on the gel that you care about (and can see because of the 32P-labeling of the 3' adapter) is actually eIF4E+the original crosslinked RNA+the 3' adapter RNA. The 3' RNA adapter is 21 nt (approximately 7 kDa) and the flag-eIF4E protein is approximately 29 kDa, for a total of ~36 kDa. We recommend that the crosslinked RNA fragments be no less than about 25 nt, and preferably at least 30 nt, so at a minimum you want to take material that is at least about 45 kDa.

As RNAs of up to 80-100 nt are OK, you should aim to cut material in the 45 kDa to ~65 kDa range.

Given this is the MW range you prefer, you want the membrane's main 'smear' of radioactivity to also correspond to this MW range. If the radioactive signal is mostly much lower than this, then it is likely you have over-digested with RNase. If the signal is a lot higher than this range, then you should try to increase your concentration (or incubation time) on the RNase step. Getting the signal 'aligned' to the MW 'sweet spot' is the best way to insure you get the best library coverage.

I generally use a clean scalpel blade to cut out each individual piece of membrane, as you do not want to cross-contaminate your individual capCLIP samples. Mark the boundaries of the membrane piece you want to cut with a Sharpie on the transparency material, and then cut right through the transparency and membrane. Carefully remove the transparency material and put the membrane piece in a fresh Eppie.

Membrane pieces can be stored dry (typically at -80˚C if not immediately processing the material).

The remaining membrane should be wrapped up in plastic wrap and stored @ -80 until the experiment is complete.

##### D2. Elution of protein:RNA from the membrane; digestion of the eIF4E protein; purification and precipitation of the isolated RNA.

Solutions needed:

2X PK buffer
200 mM Tris-Cl pH 7.5
100mM NaCl
20 mM EDTA

- ‘RNA’ phenol, pH 4.5
- “$CHCl_3$” is chloroform; no isoamyl alcohol

Proteinase K buffer:

100 µl 2X PK buffer
20 µl proteinase K (20 mg/ml)
2 µl 10% SDS
78 µl $H_2O$
200 µl total

1. Make a 2mg/ml proteinase K solution in 1X final PK buffer + 0.2% SDS;
2. pre-incubate this stock at 50° for 10 min to kill any RNAses
3. Add 200 µl of protK solution to each tube of isolated NC pieces;
4. incubate at 50°, 1200 rpm in Thermomixer for 60 min
5. Add an additional 200 µl 1X PK buffer per tube
6. Add 400 µl “RNA phenol” and 130 µl of $CHCl_3$ to solution;
7. Incubate at 37°, 1200 rpm for 5 min
8. Spin tubes at full speed in microcentrifuge (16°, 10min)
9. Carefully remove the aqueous phase to new tube
10. To this aqueous phase, add:
    60 µl 3M NaOAc pH 5.2
    2 µl of GlycoBlue (15 mg/ml)

    and then 1 ml of 1:1 EtOH:isopropanol

11. Precipitate the isolated RNA O/N at -20°

The next morning pre-chill the microfuge to 4˚C and then spin the precipitate samples for 30 min at full speed at 4˚C.

Take off the supernatants (and save each in a fresh tube!)

Wash pellets in 150 µl of 80% EtOH.

Spin full speed again for 5 minutes at 4˚C

Remove the wash with a pipette; first with P200, and then the last remaining liquid with a P20.

Invert the tubes on the bench and let each dry for about 5 minutes, being careful no to dislodge the pellets.

Return the upright tubes to a rack for another 5 minutes.

#### E. cDNA synthesis; single-stranded DNA linker-ligation; size-selection of cDNA

##### E1. Anneal primer & RT:

Make the following MQ water:SR primer mix so that each pellet can be resuspended in 4 µl volume that already includes the SR RT primer

***The SR-RT primer is used to prime cDNA synthesis from SR_RNA adapter-ligated RNA. The SR_RT DNA primer will be at a final concentration of 0.1 µM in the 10 µl total RT reaction***

***The cDNA reaction will include $^{32}$P dATP, so that the cDNA itself is radioactive. It is necessary to gel purify the final cDNA + 3'DNA adapter product, to remove cDNA products which are the result of merely ligating the 5'SRdeg_DNAv2 on to the SR-RT cDNA primer. These 'no insert' DNA products will amplify in the subsequent PCR reaction, and as they are a significant fraction of the cDNA 3' adapter ligation step, they will dominate in the PCR. We also want to remove any authentic cDNAs that are too short (~25 nt and smaller). These products cannot be reliably aligned to the genome.***

Add (per tube) on ice:
0.5 µl **SR_RT** primer @ 2 µM
3.5 µl water
4.0 µl

This mixture is generally prepared as a master mix, and the 4 µl aliquoted to each dry RNA pellet

- Resuspend each RNA pellet in 4 µl of mix.
- Heat 65°C for 5 min, the cool on ice at least 1min; quick spin to collect condensate

RT reactions:

Add (per tube) on ice:
2.0 µl 5x SuperScript IV buffer
0.25 µl RNase inhibitor
0.5 µl DTT stock
0.25 µl dNTPs (@ 20mM dCTP, dGTP ,dTTP; 8 mM dATP)
0.25 µl SuperScript IV `enzyme
1.0 µl $^{32}$P dATP
1.75 µl water
6.0 µl total volume

This is also generally made as a master mix.

Distribute 6.0 µl to each sample, mix, incubate 55° C, 10 min

***The hot dATP used is a 3.33 µM stock***
***@ 1 µl in 10 µl reaction, final hot dATP will be 0.33 µM***
***Thus the hot:cold raio is 200:0.33 or 606:1.***

***If the average cDNA is 50 nt, and avg # of A nucleotides/transcript is ~12, then ~ 1 in 50 cDNAs has a hot dATP***

**E2. Clean up cDNA:**

1. Add 2 µl ExoSAP-IT to each sample, vortex, spin down in microcentrifuge
2. Incubate 37°C for 15 mins
3. Add 1 µl 0.5 M EDTA pH 8.0
4. Add 1.5 µl of 1M NaOH, pipette-mix (make 50 ml of 1N NaOH)
5. Incubate 70°C, 12 min on heat block
6. Add 1.52 µl of 1M HCl, pipette-mix (make 50 ml of 1N HCl)
7. Put the tubes back on ice

**E3. Silane bead isolation of cDNA:**

1. Aliquot 10 µl of MyONE Silane beads per sample in a fresh Eppie, place on magnet and remove storage buffer
2. Wash each bead aliquot 1x with 250 µl RLT buffer/tube (***Qiagen #79216; Make sure to add 10 ⊠l ß-ME per ml RLT before use***)
3. Isolate beads and take off wash
4. Resuspend beads in 93 µL RLT buffer
5. Add a cleaned up cDNA sample to each set of beads:RLT buffer, mix
6. Add 111.6 µl 100% EtOH to each reaction
7. Pipette mix, let sit for 5 min, and then mix a $2^{nd}$ time 1/2 way through the 5 min
8. Place tubes on magnet, remove supernatant (save the supe in new tubes)
9. Add 500 µl 80% EtOH, pipette to resuspend beads and move all to new tube
10. After 30 sec, place tubes on magnet, remove supernatant
11. Wash 2x with 500 µl 80% EtOH (let sit 30 sec)
12. Spin briefly in picoFuge to collect wash liquid from Eppie lid
13. Place tubes back on magnet, remove residual liquid from tube with a pipette tip
14. Air-dry 5 min
15. Resuspend each dried sample in 5 µl 5 mM tris-Cl pH 8.5, let sit for 5 min (keep the bead/buffer mix as-is for the next step)

**E4. 5' linker ligate cDNA ('on-bead/with-bead'):**

- add 0.8 µl of gel purified **5'SRdeg_DNAv2** 5' DNA adapter @ 40 µM to the buffer/bead mix
- add 1 µl 100% DMSO (use tissue culture grade stock)
- heat at 75°C, 2 min, place immediately on ice for at least 1 min; quick spin

Prepare ligation master mix (on ice):
2.0 µl 10x NEB RNA Ligase Buffer (with DTT)
0.2 µl 0.1M ATP
9.0 µl 50% PEG 8000
1.0 µl RNA ligase (high conc.)
1.0 µl $H_2O$
13.2 µl

Again, make a master mix and aliquot the 13.2 µl to each tube

- add enzyme last and gently mix the master mix to homogeneity
- add 13.2 µl to each sample: alternate stirring sample with pipette tip and gently pipetting up and down, make sure solution is homogeneous
- Incubate tubes on the rotating wheel at room temp, slow rotation and near-vertical orientation of the wheel
- Incubation proceeds overnight

- NOTE: each sample is just slightly hot; easy to detect on the handheld counter, but no more than just moving the needle slightly from its resting place on the left.

**F. cDNA cleanup; size slection of cDNA on denaturing PAGE**

***The cDNA strand has a fixed sequence on its 5' end, and the cDNA product extends to (hopefully) the authentic end of the isolated RNA fragment. This fragment should still be capped, and we are aiming to get a cDNA product right to the end.***

***Regardless of the extension issue, we need to add a fixed sequence to the 3' end of the cDNA, so that we can amplify the isolated RNA fragments with PCR. This DNA adapter has a UMI of 10 nt on its 5' end, and then has a fixed sequence for amplification with the Illumina RP1 primer. The 3' end of the primer is 'blocked' so that addition DNA fragments cannot be ligated to this end. (The cDNA itself, which is primed with the SR-RT primer, does not have a phosphate at its 5' end, so it is not competent to participate in any ligation reaction, such as circularizing of the cDNA itself).***

**F1. Silane cleanup linker-ligated cDNA**

1. Aliquot 5 µl MyONE Silane Dynadbeads per sample to new Eppies, place on magnet and remove supernatant
2. Wash 1x with 250 µl RLT buffer (Qiagen buffer; remember to add fresh β-mercapto-ethanol)
3. Resuspend beads in 60 µl RLT buffer/tube
4. Add this bead/buffer miix to each overnight ligation sample, mix
5. Add 60 µl 100% EtOH to each tube
6. Pipette mix, let sit on the bench for 5 min,; then mix a second time 1/2 way through
7. Place tubes on magnet, remove supernatant (save supe in fresh tubes)
8. Add 500 µl 80% EtOH (make fresh), pipette resuspend and move to new tube
9. After 30 sec, place on magnet and remove supernatant
10. Wash another 2x with 500 µl 80% EtOH (30 sec/wash)
11. Spin briefly in picoFuge, place on magnet, remove residual liquid with fine tip
12. Air-dry 5 min
13. Resuspend in 20 µl 10 mM Tris-Cl pH 8.5
14. Let sit for 5 min on bench; then move the tubes to ice

DNA ladder labelling reaction:

***This step is***

| | |
|---|---|
| 1 µl | 10X T4PNK buffer |
| 1 µl | DNA ladder |
| 1 µl | $^{32}$P-gamma-ATP (150 mCi/ml, 25 µM; 25 pmol) |
| 6 µl | MQ water |
| 1 µl | T4 PNK |
| 10 µl | total |

1. Incubate at 37˚C for 30 min
2. Bring up rnx volume to 40 µl with MQ water

3. Set up a G25 spin column; spin out storage buffer; 736x g for 1 min
4. Add the reaction to top of G25 column, spin 735 x g for 2 min
5. Check eluate vol, bring up to 10 µl if necessary with MQ water
6. Mke some serial dilutions in MQ water of the marker to get the right abount of signal for the radioactivity of the hot cDNA

The $10^{-3}$ and/or the $10^{-4}$ dilution of ladder appears to be the best for loading. You probably should be conservative on the ladder, rather than too much (as too much signal will 'shadow' the cDNA lanes on the phosphorimager)

**F2. Gel mix and setup:**

Want to run a 7M urea, 15% acrylamide gels, with 10 lanes. Use empty Novex/Bolt casting cassettes.

**gel mix:**

| | | |
|---|---|---|
| 3 | ml | 10X TBE buffer |
| 10.66 | ml | 40% acrylamide |
| 11.85 | ml | 2% bis-acrylamide |
| 12.61 | g | urea |

Mix, add stir bar and stir w/ heat until urea dissolved

Bring volume to ~36 ml total with by adding another 600 µl of 10X TBE

So, the final gel percentage is 12.5% acrylamide

Add 400 µl 10% APS + 20 µl of TEMED to gel mix; swirl to mix and quickly pour 2 Bolt gels w/ 10-well combs using this volume of mix.

After polymerization, prep the bolt gels for running and purge the wells of excess urea;

**gel loading:**

Set up Novex apparatus with the 1-2 gels, depending on the # of samples to run, filling both reservoirs with 1X TBE.

Pre-run for 10 min at 180 V constant.

DO NOT preheat the marker; if you do, then the marker will not run to size.

Add a urea loading dye to each sample; run the entire 20 µl of resuspended cDNA + DNA linker in one lane (DO NOT pre-heat these samples either)

Run the gel at 200 V constant, and run until the BB maker has just left the gel (and the XC is at about 1/2-way down the gel).

Crack the cassettes and carefully transfer the gel to a small piece of overhead transparency. Cover in glad wrap.

Prepare strips of Whatman for spotting of diluted $^{32}P$ marker (as indicated on the strips themselves).

Arrange with the gels on a larger piece of Whatman paper and then secure all together with tape. Make sure that gels and the co-registration strips all fit on the phosphorimage film size and place in cassette overnight at room temp.

DO NOT freeze the gels!

**F3. Gel slice cutting and cDNA elution from the gel:**

1. In general the phosphorimage scans of hot cDNA gels will give a weak signal, but the hot material should be exactly at the size you would predict from the MW of the eIF4E:RNA material that you excised from the nitrocellulose blots, plus the added MW of the 5' DNA linker

2. Just as you did for the initial nitrocellulose phosphorimages, make overhead transparency copies of the gel images, and cut out bands from ~75 to ~200 base pairs. This is a bit wide on both the short and long ends, but it is better to take too much material here than too little. In general, the gel purification is most important for removing "no insert" cDNA products from subsequent PCR amplification of the libraries.

3. Take the gel slices to fresh eppies—we usually use screw cap eppies for this step. Add 350 µl nucleic acid elution buffer to each tube.

4. Using a 1 ml syringe plunger, crush each acrylamide slices into a slurry

5. Let the slurry sit in the tube overnight. Place tubes on Thermomixer at 37˚ C and 800 rpm.

**F4. Spin column filtration of the eluted cDNA; EtOH precipitation of the cDNA**

1. Set up Costar columns in fresh collection eppies with Millipore glass filters placed on top of the column matrix to prevent clogging of the Costar filter with the acrylamide slurry
2. Add the slurry to the column with a blunt P1000 tip; spin full speed for 5 min; measure eluate volume in collection eppies (volumes ~360 µl)
3. Transfer eluate to new Eppies and add 2 µl of glycoblue + 1000 µl 100% Ethanol
4. Precipitate the cDNA @ -20˚C for at least overnight

**F5. Further work-up of the gel purified, precipitated cDNA**

1. Spin ppt samples at top speed for 30 min in 4˚C microfuge
2. Wash with 100 µl 80% EtOH
3. Spin full speed 10 mins, 4° C
4. Take off wash; air dry pellet on bench
5. Resuspend each dried pellet in 40 µl MQ water

**G. PCR amplification of cDNA using index primers**

**G1. Estimating the # of PCR cycles to correctly amplify the cDNA**

***A critical step of capCLIP is to do enough rounds of PCR to have just enough DNA for the sequencer, without overcycling the library and loosing library complexity at the expense of more moles of PCR product.***

***In general, you want about 20 fmol of DNA per 80-base Nextseq 500 run. Thus, if you want to sequence 10 individual sample libraries, you want to have 2 fmol of each library for the run.***

***The best approach to demining the proper # of amplification cycles for each library is to do test PCRs for each and every sample, using just a small fraction of the cDNA you have just prepared.***

***Step F5 has the cDNA of each sample resuspended in 40 µl of MQ water, so we recommend to always try at least 1 set of test PCRs with ~10% to 20% of the total cDNA***

From experience, the best way to preserve library complexity is to a **single step of PCR amplification**, meaning that you never want to do a PCR reaction, find that you don't have enough for sequencing, and then attempt to do a few more rounds of PCR on the already amplified material. This re-PCR step never works predictably, and you want to never be in the position of having used up all of your original cDNA, and having to do more cycles of PCR on previously amplified material

It is critical to do test PCRs,and get the proper # of cycles worked out.

We have also had the best luck with doing one more gel purification after the PCR step, as there will still be some "no insert" PCR product in your PCR reactions, despite the cDNA gel purification step. So you also need to factor in the amount of material that will be lost with this final purification step.

Thus, if you have a really valuable sample, do a test PCR and also do a test gel purification, and then use a Qubit + Agilent to see how you are doing.

**G2. PCR for library amplification:**

This 'test' reaction will use 8 µl of each cDNA prep (so 20%).

Reaction mix with Ultra II Q5 master mix: (per 25 µl reaction)
8 µl gel purified cDNA
1.56 µl SR (aka RP1) @ 20 µM
1.25 µl **NEBNext index primer** @ 25 µM
1.69 µl MQ water
12.50 µl Q5 high fidelity master mix
25 µl total

***As each individual sample requires a separate NEBNext index primer, you will need to set up each PCR reaction individually.***

Cycle # for the PCR reactions:

If one were doing an experiment very similar to what is described here, we would recommend that the initial PCR should be done for somewhere between 12-14 cycles, but this will vary a lot with the experimental system.

**Cycle parameters:**

98° C for 30 sec

---

98° C for 10 sec
72° C for 30 sec
65° C for 45 sec

---

65° C for 5 min final extension
hold @ 4°C

If you don't want to work up the samples, store samples@ -20°C

Set up another empty Bolt cassette for a 7M urea, 12.5% acrylamide gel, using a 10-well cassette

**G3. Gel purification of PCR products**

Gel mix for PCR product purification gels:
3 ml 10X TBE buffer
10.66 ml 40% acrylamide
11.85 ml 2% bis-acrylamide
12.61 g urea

-mix, add stir bar and stir w/ heat until urea dissolved

-volume should be ~35+ ish ml, and 600 µl of 10X TBE for a total of 36 ml of gel mix and a final acrylamide of 12.5%

-add 400 µl '10%' APS and 20 µl of TEMED (both 'fresh') and pour gel quickly (the gel mix volume is sufficient for 2 Bolt gels

Gel loading:

1. To each PCR tube, add 25 µl of NEB RNA loading dye/buffer (Load entire sample/well on gel)
2. run 40 ng of the pBR322/MspI digest on each side of each gel (2 µl of 20 ng/µl pBR ladder dilution in 1X Cut Smart buffer, plus 5 µl of NEB RNA load dye)
3. If you have any 'blank' lanes, make an Eppie tube of 1:1 1X Cut Smart buffer:1X NEB RNA load buffer and load 30 µl/lane (so need 6 x 30 µl)

Unlike for the cDNA gels, **pre-heat samples and ladder at 70 °C for 5 min**, then place them back on ice to chill. Do a quick spin with the tubes and load samples at room temp.

Run the gel at 160 V until the XC band is 3/4 or farther down the gel

For visualisation of the DNA, make up 1:10,000 SYBR gold dilution in 50 ml of 1X TBE; filter the mix (particulates in the stain are very bright on gel imaging)

Carefully open each cassette with **gloved hands**, do not use powdered gloves

Chop off the wedge wells and the bottom lip of gel; make sure orientation is clear on each naked gel

Carefully place the gel into the new staining tray that is pre-filled with the 1X

TBE/SYPR gold stain (below); stain for ~30 minutes

Image gel on a BioRad GelDoc

As for previous gel images, you will want to transfer the life-sized gel image to transparency film so that you can cut bands on the gel. [While you may be able to see the bands on a UV light box, we do not recommend this approach, both for the additional UV exposure and for the usually contaminated surfaces of such equipment.

***Never freeze a gel before imaging. While there used to be a product call GelBond that would allow you to store a gels @ -80˚C and then thaw if without the gel falling apart, if you do so with transparency film as the substrate you will get back a destroyed gel upon thawing. It is possible to put a gel @ 4˚C for a bit, but best to do cut bands immediately.***

<u>Cutting bands:</u>
This PCR step adds primer flanking sequence of 128 base pairs, so you will want to cut PCR products that run at 128 nt + the range of RNA inset sizes you desired from your cDNA sizing gel.

Generally, what we see at this point looks like this:

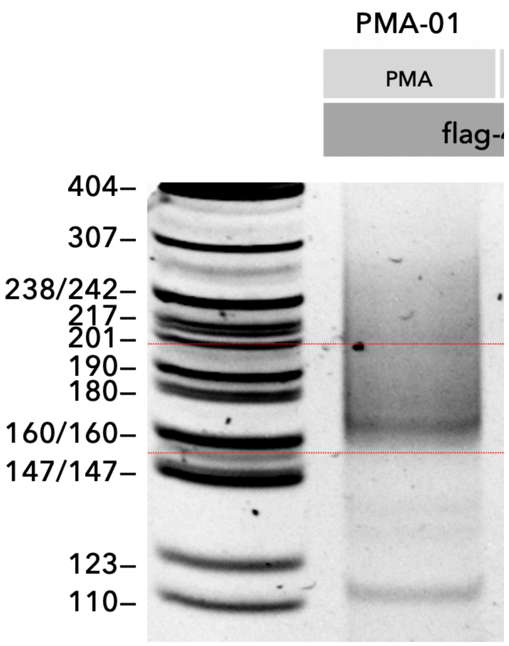

There is a smear of PCR product from about ~160 nt up to just under 300 nt. The red lines indicate the range of product that we isolated for sequencing (160-200 nt). This corresponds to insert sizes of ~30 to 70 nt. You can see a prominent band at ~120 nt – this is almost certainly the “no insert” PCR product (which is 128 nt).

<u>Gel slice isolation</u>

1. Use clean scalpel blades to remove your gel slices, and add each to a screw-cap Eppie.
2. Add 350 µl nucleic acid elution buffer and use 1 ml string plunger to crush

each acrylamide slice to a slurry; elute at 37˚ C and 800 rpm in thermomixer overnight (at least 18 hours and up to 24 hour)

Spin column filtration of PCR product; EtOH ppt.

- Around 18-24 hours post-incubation transfer the gel slurry from each tube to ‘Costar’ columns with glass filters on top (as before)
- Spin full speed for 5 min; measure eluate volume (volume should be ~360 µl)
- Transfer eluate to a new Eppie and add 2 µl of glycoblue + 1000 µl 100% Ethanol
- Precipitate overnight at -20˚C

The next day:

- Spin ppt samples at top speed for 30 min in 4˚C microfuge
- Wash with 100 µl 80% EtOH
- Spin full speed 10 mins, 4° C
- Air dry pellet.
- Resuspend each pellet in 20 µl MQ water for QC analysis on the Qubit and Tapestation/Agilent

**G4. Qubit quantification and Tapestation QC:**

Qubit: Pepare the Qubit working solution by diluting the Qubit dsDNA HS Reagent 1:200 in Qubit dsDNA HS Buffer. Measure 2 µl of each DNA sample; into a fresh thin walled 500 µl PCR tube add 2 µl of DNA sample + 198 µl of Qubit working solution. Mix, quick centrifuge, and then place samples in dark for at least 2 minutes. Then read on Qubit.

Tapestation: Get ScreenTape and loading reagents out of fridge and equilibrate to room temp (~30 min.) Mix 2 µl of each DNA sample + 2 µl of loading buffer; vortex; quick centrifuge; load tape. Do the same with 2 µl of the appropriate DNA marker

As the sequencing service you use is likely to get involved in the process at this point, we won’t say a whole lot further about library QC.

It is important that you use the Agilent or Tapestation to verify the size of your purified PCR products, and to use the Qubit to get an accurate concentration.

Depending on the # of individual libraries, the sequencing machine you use, etc. how you go from here is too variable to provide useful commentary—talk to your sequencing people!
